# CD4^+^ T-cells sensitize quasi-mesenchymal breast tumors lacking CD73 to anti-CTLA4 immune checkpoint blockade therapy

**DOI:** 10.1101/2025.05.12.653467

**Authors:** Shiney Chandraganti, Caitlyn Sams, Sarthak Sahoo, Brian Feng, Isabel O’Connell, Lynna Li, Sunita Nepal, Siddhartha Pulukuri, Kimaya Bakhle, Prathapan Thiru, Corina Simian, George W. Bell, Mohit Kumar Jolly, Anushka Dongre

## Abstract

Although immune checkpoint blockade therapy has generated dramatic responses in certain cancer types, breast tumors are largely unresponsive. Epithelial-mesenchymal plasticity leads to the assembly of an immunosuppressive tumor microenvironment and drives resistance of breast tumors to immunotherapies. Importantly, targeting CD73 completely sensitizes quasi-mesenchymal breast tumors to anti-CTLA4 immune checkpoint blockade therapy. However, the mechanism(s) of sensitization remained unknown. We demonstrate that targeting CD73 in quasi-mesenchymal breast tumors sensitizes them to anti-CTLA4 immune checkpoint blockade therapy in a CD4^+^ T-cell dependent manner. Moreover, epithelial-mesenchymal plasticity results in elevated expression of cancer cell-intrinsic CD73 in human triple negative breast cancers. Given the ability of quasi-mesenchymal cancer cells to metastasize and resist multiple therapies, these findings can instruct the formation of novel translational strategies for the treatment of human breast cancers. These findings also bring to the forefront the attractive possibility of utilizing the phenotypic plasticity of cancer cells along with CD73 and CD4^+^ T-cells as a predictive criterion for immunotherapy responsiveness.

**Teaser:** Targeting CD73 sensitizes breast cancer cells with mesenchymal traits to anti-CTLA4 therapy in a CD4^+^ T-cell dependent manner.

## Introduction

The use of immune checkpoint blockade therapies (ICB), which harness the immune system to kill cancer cells, has revolutionized cancer treatment by creating durable clinical responses (*1*). However, while melanomas and lung cancers mount proficient responses to these therapies, certain other cancer types, such as breast carcinomas, are still largely unresponsive (*2–5*). It is therefore critical that we drastically improve the curative potential of these therapies by understanding the mechanisms by which breast cancer cells mount resistance to immunotherapy.

Epithelial-Mesenchymal Plasticity (EMP), which converts epithelial cells to more-mesenchymal derivatives, endows cancer cells with many traits associated with high grade malignancies. This includes their ability to metastasize to distant organ sites, acquire tumor-initiating abilities, and mount resistance to chemotherapies (*6–9*). In fact, EMP is a highly dynamic process which often gives rise to a spectrum of partial or quasi-mesenchymal (qM) states which can co-express both, more-epithelial and more-mesenchymal markers (*10–12*). Importantly, EMP is a reversible process in which more-mesenchymal cancer cells can lapse back into a more-epithelial state by undergoing a mesenchymal-to-epithelial transition or MET (*13*). In addition to these well-documented features of this program, we have shown that EMP can lead to the assembly of an immunosuppressive tumor microenvironment (TME) and render breast tumors unresponsive to ICB therapies (*14–17*).

In previous work, we established epithelial or qM cell lines from tumors arising in the MMTV-PyMT autochthonous murine model of breast cancer (*18*). Some of these mice contained IRES-YFP reporter constructs that labelled cells expressing the Snail EMT transcription factor enabling us to isolate Snail^HI^ qM cancer cells which differed from their Epcam-expressing epithelial counterparts (*18, 19*). By implanting these cell lines into immunocompetent, syngeneic hosts, we established novel, pre-clinical murine models of epithelial or qM breast tumors and observed that epithelial tumors recruit CD8^+^ T-cells to the tumor microenvironment and are highly responsive to anti-CTLA4 ICB. In sharp contrast, qM tumors recruit immunosuppressive cells such as T-regulatory cells and M2-like macrophages instead and are resistant to the same therapy (*18*). Most strikingly, in mixed tumors comprised of both epithelial and qM cancer cells, a minority population (10%) of more-mesenchymal cells can cross-protect the vast majority (90%) of their epithelial neighbors from immune attack (*18*). These observations alone, are of great consequence clinically, since majority of human carcinomas contain minority populations of more-mesenchymal cells that could dictate the outcome of the entire tumor to immune attack.

This ability of qM cancer cells to resist being eliminated by the immune system is in part due to their secretion of multiple immune-suppressive factors (*20*). Of particular importance, is the observation that qM cancer cells that lack the expression of a specific immune-suppressive factor called CD73 (an ectoenzyme that produces immunosuppressive adenosine), were completely sensitized to anti-CTLA4 ICB (*20*). Additionally, targeting the adenosinergic signaling pathway in qM tumors with either anti-CD73 or an adenosine receptor antagonist generated synergistic responses with anti-CTLA4 leading to a significant reduction in primary tumor size as well as distal metastases. Additionally, these strategies sensitized qM breast tumors specifically to anti-CTLA4 but not anti-PD1 ICB therapy (*20*). Our previous work demonstrated for the very first time that disruption of certain cancer cell-intrinsic, EMP-regulated immune-suppressive signaling channels, notably CD73, could lead to a near complete eradication of qM cancer cells.

Given that more-mesenchymal cells enable metastasis and are notoriously resistant to multiple treatment regimens, strategies to eliminate them altogether could revolutionize cancer treatment. However, the mechanism(s) underlying such sensitization remains unknown. In other words, the identity of immune cell subsets that mediate the elimination of qM cancer cells lacking CD73 in response to anti-CTLA4 ICB treatment is yet to be determined. We present our findings demonstrating that CD4^+^ T-cells sensitize qM breast tumors that lack the expression of CD73 to anti-CTLA4 ICB. Importantly, CD73 expression is associated with a partial and more-mesenchymal state in human breast cancer cell lines and patient samples. Our work brings to the forefront the attractive possibility of utilizing CD4^+^ T-cells, CD73, and EMP as predictive criterion for ICB responsiveness of breast tumors.

## Results

### Presence of T-cells in responding tumors

In previous work, we established novel E and qM breast cancer cell lines by sorting cells from tumors arising in the autochthonous MMTV-PyMT murine model bearing Snail-IRES-YFP knock-in constructs (*18*). We have demonstrated that qM cancer cells lacking the expression of CD73 (hereafter referred to as sgCD73) are completely eliminated after treatment with anti-CTLA4 ICB relative to qM control tumors which are resistant (*20*) (fig. S1A, B). However, precisely which immune cells mediate this elimination was unknown. We first determined how the absence of CD73 from qM cancer cells alters the tumor microenvironment even before the administration of anti-CTLA4 ICB, by performing multiplexed scRNA-Seq analysis of qM control and sgCD73 tumors. We identified eleven clusters representing B-cells, CD4^+^ T-cells, CD8^+^ T-cells, two subsets of macrophages, neutrophils, fibroblasts, endothelial cells, mesenchymal progenitor cells and two types of cancer cells (Fig. 1, A and B and Supplementary Table 1). Of these various cell types, sgCD73 tumors contained elevated numbers of CD4^+^ T-cells, followed by B-cells and a subset of cancer cells relative to qM control tumors even prior to ICB treatment (Fig. 1C).

**Figure 1:**
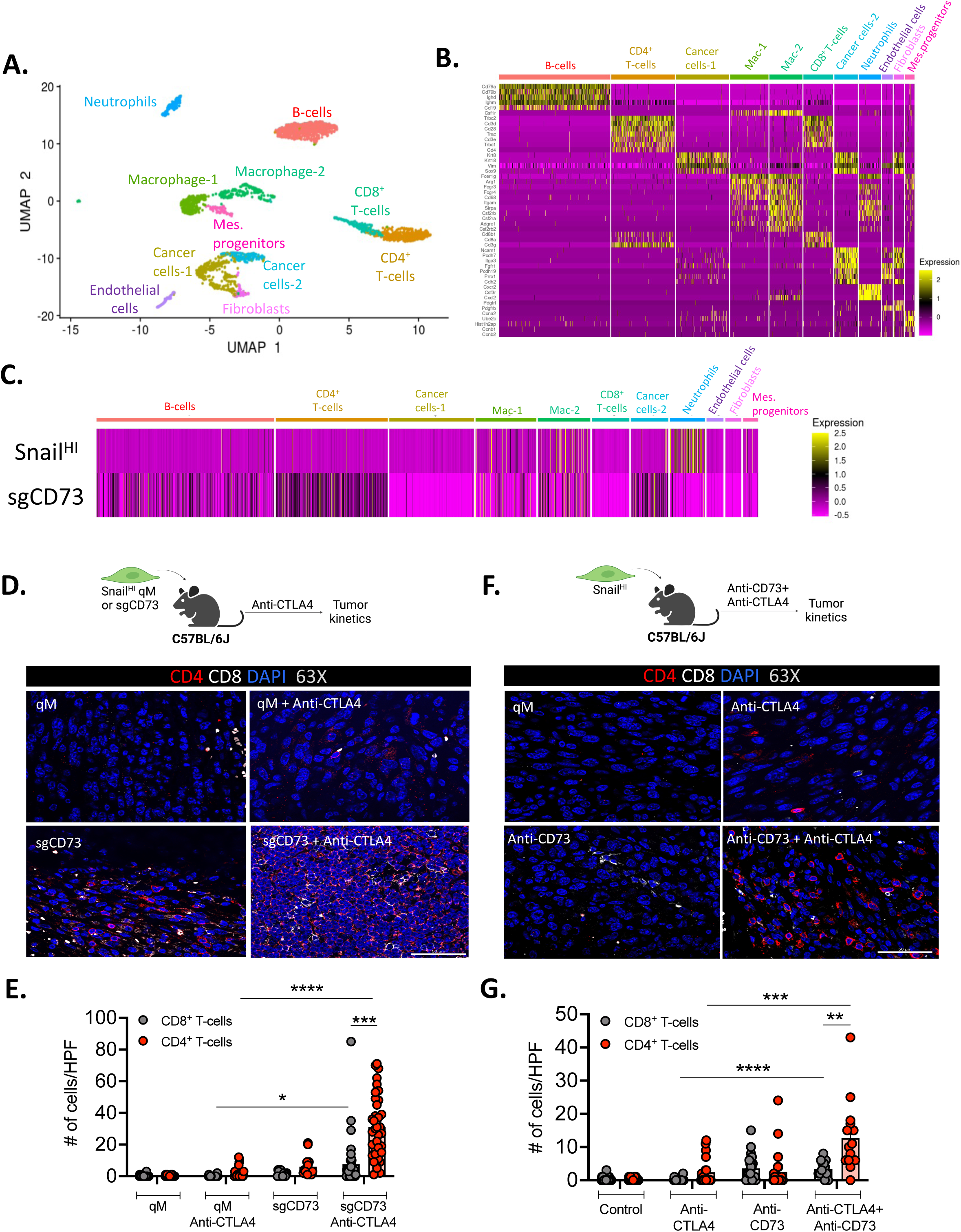
Presence of T-cells in responding tumors: **(A)** UMAP plot of unbiased clustering of Snail^HI^ qM and sgCD73 tumors, where each color-coded cluster represents a specific cell type or state. **(B)** Genes representing each cluster depicted in the UMAP plot in (A). See additional details in Supplementary Table 1 **(C)** Representation of each cluster in Snail^HI^ qM and sgCD73 tumors. Expression levels represent log transformed values. **(D)** Immune-fluorescence analysis of primary tumor sections obtained from Snail^HI^ qM or sgCD73 tumor-bearing mice receiving Control or anti-CTLA4 antibodies stained for CD4 (red), CD8 (white) and DAPI (blue) at 63X magnification. (**E**) Quantification of CD8^+^ T-cells and CD4^+^ T-cells in each high-power field (HPF) at 63X magnification from (D). Data represent three independent experiments, with n=3-5 mice in each group. 3-5 fields of view from the tumor interior were obtained for each group at 63X magnification. **(F)** Immune-fluorescence analysis of primary tumor sections obtained from Snail^HI^ qM tumor-bearing mice receiving Control, anti-CD73, anti-CTLA4 or combinations of anti-CD73 and anti-CTLA4 antibodies stained for CD4 (red), CD8 (white) and DAPI (blue) at 63X magnification. (**G**) Quantification of CD8^+^ T-cells and CD4^+^ T-cells in each high-power field (HPF) at 63X magnification from (**F**). Data represent three independent experiments, with n=3-5 mice in each group. 3-5 fields of view from the tumor interior were obtained for each group at 63X magnification. (**E, G**) Data represent SEM, two-tailed unpaired t-test, *\*, p<0.05, **, p<0.01, ***, p<0.001, ****, p<0.0001*.

Given the ability of anti-CTLA4 ICB to regulate the recruitment and function of both, CD8^+^ and CD4^+^ T-cells, we focused our analyses on T-cells and asked whether the proportion of both these subsets was altered in sgCD73 and qM tumors after treatment with anti-CTLA4 ICB. Immunofluorescence staining of tumor sections revealed that responding tumors (sgCD73 treated with anti-CTLA4) recruited significantly higher numbers of both, CD8^+^ and CD4^+^ T-cells to the tumor core relative to control qM tumors that were unresponsive to ICB therapy (Fig. 1, D and E and fig. S1A, B). We have previously determined that treating Snail^HI^ qM tumor-bearing mice with anti-CD73 in combination with anti-CTLA4 ICB generates synergistic responses resulting in significantly smaller tumors relative to mice receiving each antibody individually (*20*) (fig. S1C and S1D). Accordingly, Snail^HI^ qM tumor-bearing mice that received combinations of anti-CD73 and anti-CTLA4, also demonstrated an increased influx of both CD8^+^ and CD4^+^ T-cells relative to untreated tumors or those that received each therapy individually (Fig. 1F and G). What was particularly striking in both models was that although responding tumors recruited both T-cell types, the number of CD4^+^ T-cells in the tumor core was significantly higher than the number of CD8^+^ T-cells (Fig. 1E and G).

### CD8^+^ T-cells are partially important for regulating responses of qM tumors lacking CD73 to anti-CTLA4 ICB

To understand which T-cell subset was functionally important in regulating responses of sgCD73 tumors to anti-CTLA4 ICB, we first depleted CD8^+^ T-cells using subset-specific antibodies. Depletion of CD8^+^ T-cells from sgCD73 tumor-bearing mice only partially reversed sensitivity to anti-CTLA4 ICB (Fig. 2A and fig. S1E)(*20*). Moreover, sgCD73 tumor-bearing mice continued to recruit CD4^+^ T-cells to their primary tumors in response to anti-CTLA4 ICB, even in the absence of CD8^+^ T-cells (Fig. 2B). To validate these findings further, we implanted sgCD73 cells into genetically modified mice that lacked all CD8^+^ T-cells (CD8-KO) or alternatively, in mice which lacked B2M (B2M-KO). B2M is required for the stable cell surface expression of MHC-I, which is critical for antigen presentation to CD8^+^ T-cells. Thus, in this later scenario, CD8^+^ T-cells while still present, are functionally compromised due to the absence of priming. Strikingly, sgCD73 tumors propagated in both, CD8-KO and B2M-KO mice continued to respond to anti-CTLA4 ICB just as well as those propagated in Wild Type (WT) mice that also received ICB treatment (Fig. 2, C and E and fig. S1F and S1G). Additionally, a large number of CD4^+^ T-cells were found within the tumor core of responding tumors that grew in both, CD8-KO and B2M-KO mice upon treatment with anti-CTLA4 ICB, suggesting that the absence or functional impairment of CD8^+^ T-cells did not affect the recruitment of CD4^+^ T-cells to responding tumors (Fig. 2, D and F).

**Figure 2:**
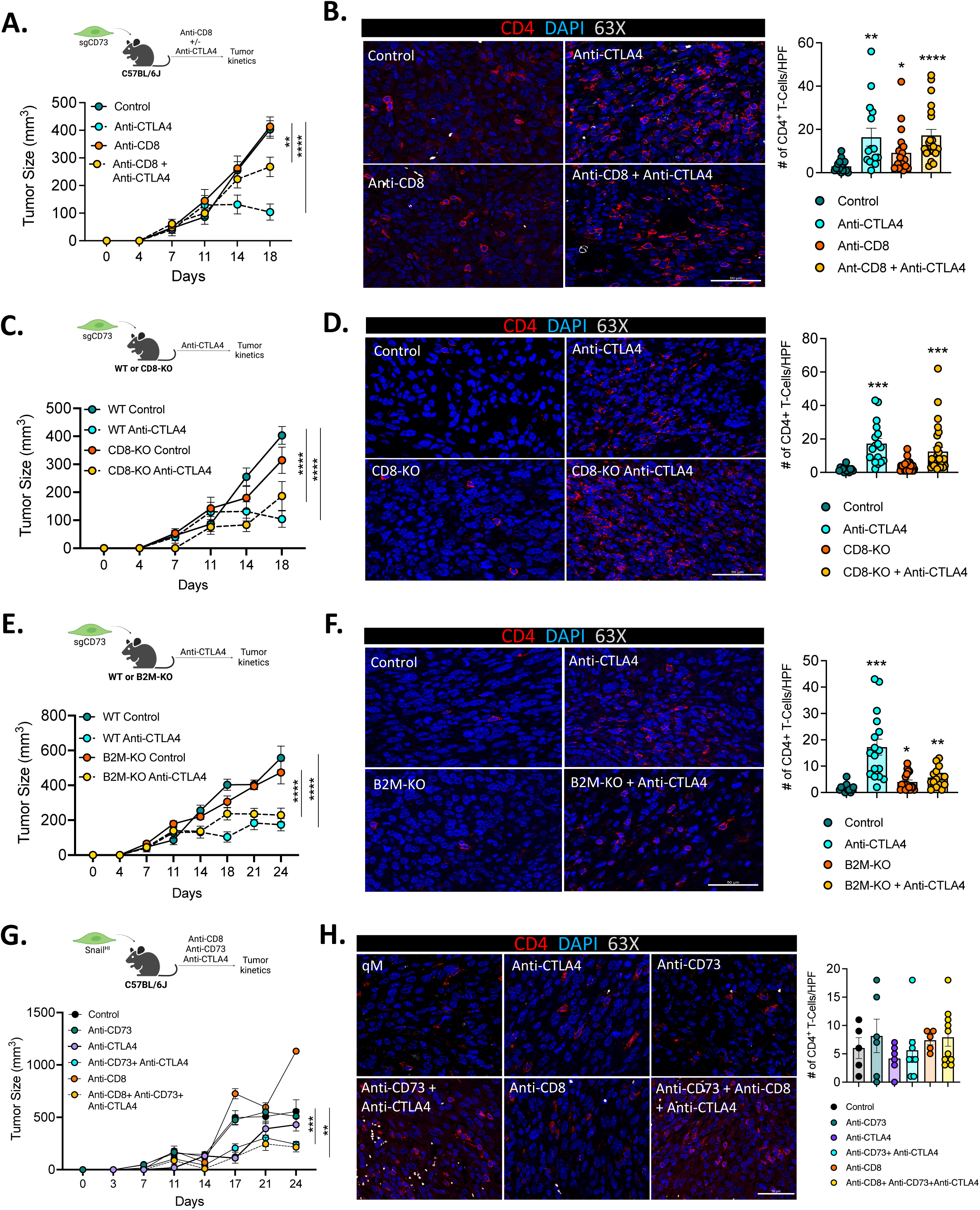
CD8^+^ T-cells are partially important for sensitizing qM tumors lacking CD73 to anti-CTLA4 ICB: **(A)** Schema and tumor kinetics for sgCD73 tumor-bearing mice treated with the indicated antibodies. Data represent three independent experiments where n=3-4 for each group **(B)** Representative immune-fluorescence images of primary tumor sections obtained from (A) stained for CD4 (red) and DAPI (blue) at 63X magnification. Bar graph on the right represents quantification of CD4^+^ T-cells in each high-power field (HPF) at 63X magnification for the indicated treatment groups. 3-5 fields of view from the tumor interior were obtained for each tumor. Data represent three independent experiments, with n=3-5 mice in each group. **(C)** Schema and tumor kinetics for sgCD73 tumors propagated in Wild Type (WT) or CD8 knock-out (CD8-KO) mice treated with the indicated antibodies. Data represent three independent experiments where n=3-4 for each group **(D)** Representative immune-fluorescence images of primary tumor sections obtained from (C) stained for CD4 (red) and DAPI (blue) at 63X magnification. Bar graph on the right represents quantification of CD4^+^ T-cells in each high-power field (HPF) at 63X magnification for the indicated treatment groups. 3-5 fields of view from the tumor interior were obtained for each tumor. Data represent three independent experiments, with n=3-5 mice in each group. (**E**) Schema and tumor kinetics for sgCD73 tumors propagated in Wild Type (WT) or B2M knock-out (B2M-KO) mice treated with the indicated antibodies. Data represent three independent experiments where n=3-5 for each group **(F)** Representative immune-fluorescence images of primary tumor sections obtained from (E) stained for CD4 (red) and DAPI (blue) at 63X magnification. Bar graph on the right represents quantification of CD4^+^ T-cells in each high-power field (HPF) at 63X magnification for the indicated treatment groups. 3-5 fields of view from the tumor interior were obtained for each tumor. Data represent three independent experiments, with n=3-5 mice in each group. (**G**) Schema and tumor kinetics for Snail^HI^ qM tumor-bearing mice treated with the indicated antibodies. Data represent two independent experiments where n=3-4 for each group. (**H**) Representative immune-fluorescence images of primary tumor sections obtained from (G) stained for CD4 (red) and DAPI (blue) at 63X magnification. Bar graph on the right represents quantification of CD4^+^ T-cells in each high-power field (HPF) at 63X magnification for the indicated treatment groups. 3-5 fields of view from the tumor interior were obtained for each tumor. Data represent three independent experiments, with n=3-5 mice in each group. (**A, C, E, G**) Data represent SEM, two-way ANOVA, *\*\*, p<0.01, ***, p<0.001, ****, p<0.0001.* (**B, D, F, H**) Bar graph data represent SEM, two-tailed unpaired t-test, *\*, p<0.05, **, p<0.01, ***, p<0.001, ****, p<0.0001.* Scale bars are 100um.

Finally, we asked whether synergistic responses observed by using anti-CD73 in combination with anti-CTLA4, were also dependent on CD8^+^ T-cells. Antibody based depletion of CD8^+^ T-cells from Snail^HI^ qM tumor-bearing mice did not alter their responses to combination therapy (anti-CD73 and anti-CTLA4). In other words, these tumor-bearing mice mounted synergistic responses to combination therapy and recruited CD4^+^ T-cells to the tumor core just as efficiently in the presence and absence of CD8^+^ T-cells (Fig. 2, G and H and fig. S1H). Taken together, our data demonstrate that targeting CD73 sensitizes qM tumors to anti-CTLA4 in a manner that is only partially dependent on CD8^+^ T-cells. Moreover, the absence of CD8^+^ T-cells does not impact the recruitment of CD4^+^ T-cells to responding tumors.

### CD4^+^ T-cells drive sensitivity of qM tumors lacking CD73 to anti-CTLA4 ICB

While CD8^+^ T-cells have been ascribed as the key players of the adaptive immune system in driving anti-tumor immune responses, the functional importance of anti-tumor CD4^+^ T-cells is only beginning to emerge. Given the large influx of CD4^+^ T-cells in both responding tumor models, we asked whether they were functionally important for sensitizing qM tumors lacking CD73 to anti-CTLA4 ICB. Antibody-based depletion of CD4^+^ T-cells from sgCD73 tumor-bearing mice completely reversed their responsiveness to anti-CTLA4 ICB (Fig. 3A and fig. S2A)(*20*).

**Figure 3:**
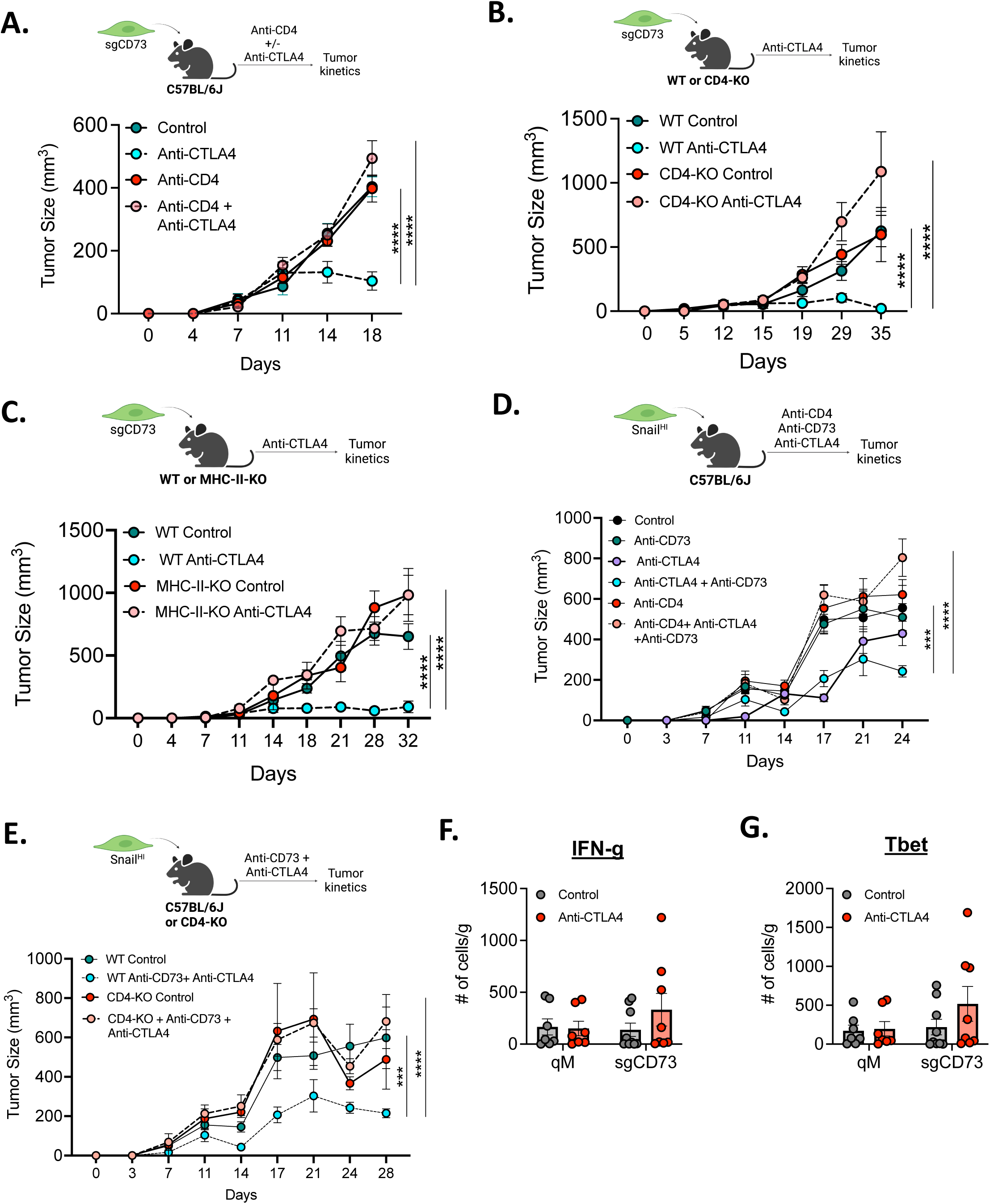
CD4^+^ T-cells drive sensitization of qM tumors lacking CD73 to anti-CTLA4 ICB: (A) Schema and tumor kinetics for sgCD73 tumor-bearing mice treated with the indicated antibodies. Data represent 3 independent experiments where n=3-7 for each group **(B)** Schema and tumor kinetics for sgCD73 tumors propagated in Wild Type (WT) or CD4 knock-out (CD4-KO) mice treated with the indicated antibodies. Data represent 3 independent experiments where n=3-7 for each group (**C**) Schema and tumor kinetics for sgCD73 tumors propagated in Wild Type (WT) or MHC-II knock-out (MHC-II-KO) mice treated with the indicated antibodies. Data represent 3 independent experiments where n=3-7 for each group (**D**) Schema and tumor kinetics for Snail^HI^ qM tumor-bearing mice treated with the indicated antibodies. Data represent 3 independent experiments where n=3-7 for each group (**E**) Schema and tumor kinetics for Snail^HI^ qM tumors propagated in Wild Type (WT) or CD4 knock-out (CD4-KO) mice treated with the indicated antibodies. Data represent 3 independent experiments where n=3-7 for each group (**F, G**) Flow cytometry analysis for (**F**) IFN-gamma and (**G**) T-BET from Snail^HI^ qM control or sgCD73 tumor-bearing mice treated with the indicated antibodies. Data represent 3 independent experiments where n=3-7 for each group (**A-E**) Data represent SEM, two-way ANOVA, *\*\*, p<0.01, ***, p<0.001, ****, p<0.0001.* (**F, G**) Bar graph data represent SEM, two-tailed unpaired t-test, *\*, p<0.05, **, p<0.01, ***, p<0.001, ****, p<0.0001*.

Similarly, sgCD73 tumors failed to respond to ICB when grown in genetically modified mice that lacked all CD4^+^ T-cells (CD4-KO) (Fig. 3B and fig. S2B) relative to WT control mice. CD4^+^ T-cells are activated when their T-cell receptor recognizes antigens presented by MHC-II molecules. Along these lines, genetically modified mice lacking the expression of MHC-II molecules have impaired CD4^+^ T-cell responses. To further confirm the functional importance of CD4^+^ T-cells in driving sensitization of sgCD73 tumors to anti-CTLA4 ICB, we propagated sgCD73 cancer cells in WT or MHC-II-KO mice. Strikingly, sgCD73 tumors failed to respond to anti-CTLA4 ICB even when propagated in MHC-II-KO mice, in sharp contrast to WT mice, where they mounted proficient responses to the same treatment (Fig. 3C and fig. S2C).

Finally, Snail^HI^ qM tumor-bearing mice failed to respond to combination therapy of anti-CD73 and anti-CTLA4 ICB when CD4^+^ T-cells were depleted (Fig. 3D and fig. S2D). Similarly, Snail^HI^ qM tumors failed to respond to combinations of anti-CD73 and anti-CTLA4 when orthotopically implanted in CD4-KO mice (Fig. 3E and fig. S2E), relative to WT mice where they responded proficiently to combination therapy. Taken together, these findings from multiple models enabled us to determine that CD4^+^ T-cells are necessary for sensitizing qM breast tumors lacking CD73 to anti-CTLA4 ICB.

CD4^+^ T-cells are heterogeneous and can differentiate into multiple subsets, including anti-tumor, immune-stimulatory T_H_1 and T_H_17 cells or pro-tumor, immune-suppressive T_H_2 and T-regulatory cells (*21*). Whether the abrogation of CD73 from qM cancer cells altered the representation of one or more T-cell subsets remained unknown. This is particularly important as the polarization status of CD4^+^ T-cells can directly influence breast tumor progression and could have profound consequences on their subsequent responsiveness to ICB therapies. Thus, we determined the identities of various CD4^+^ T-cell subsets in control qM and sgCD73 tumors pre and post treatment with anti-CTLA4 ICB. sgCD73 tumors demonstrated an increase (albeit not significant) in the absolute numbers of IFN-gamma and T-BET-expressing CD4^+^ T-cells only in response to anti-CTLA4 ICB relative to untreated tumors or qM tumors that were unresponsive to ICB treatment (Fig. 3, F and G). Surprisingly, the representation of other T-cell subsets T_H_2, T_H_17, Tregs, T-follicular helper cells, as well as CD4^+^ T-cells expressing Granzyme A, B, or TNF-alpha was unaltered in responders and non-responders (fig. S3A-I) pre and post anti-CTLA4 ICB therapy. Moreover, Snail^HI^ qM and sgCD73 tumors did not show statistically significant differences in the proportion of CD4^+^ T-cells expressing markers associated with proliferation or dysfunction such as KLRG1, Ki67, CTLA-4, PD-1, TIGIT, TIM3 and 4-1BB (fig. S4A-H). Thus, sgCD73 tumors demonstrate elevated numbers of T_H_1-like CD4^+^ T-cells in response to anti-CTLA4 ICB.

### Abrogation of MHC-I from qM tumors lacking CD73 drives sensitivity to anti-CTLA4 ICB in a CD4^+^ T-cell dependent manner

We and others have previously demonstrated that qM cancer cells significantly reduce their cell surface expression of MHC-I as a consequence of activating the EMP program, which could in turn render sgCD73 tumors vulnerable to NK cell-mediated cytotoxicity (*20*). Moreover, myeloid cells are also capable of engulfing MHC-deficient tumors. Thus, sensitization of sgCD73 tumors to anti-CTLA4 ICB could be driven by NK and/or myeloid cells, in addition to CD4^+^ T-cells.

To functionally validate the relevance of these other immune cells in driving sensitization of sgCD73 tumors to anti-CTLA4 ICB, we abrogated the expression of MHC-I from sgCD73 cancer cells (via CRISPR/Cas9 mediated deletion of B2M, which is required for the stable cell-surface expression of MHC-I) (fig. S5A). While control sgCD73 cancer cells upregulated MHC-I and B2M in response to IFN-gamma treatment *in vitro*, double knockout cells (DKO) which lacked both, CD73 and B2M, failed to do so (fig. S5B and S5C). Accordingly, such a strategy would render DKO cells unresponsive to elimination by CD8^+^ T-cells while retaining their susceptibility to macrophages, NK cells, and CD4^+^ T-cells. These DKO cells were implanted orthotopically into syngeneic immune-competent hosts and treated with control antibodies or anti-CTLA4 ICB. Strikingly, tumor-bearing mice that contained DKO cells which lacked both CD73 and B2M, were completely sensitized to anti-CTLA4 ICB (Fig. 4A and fig. S5D).

**Figure 4:**
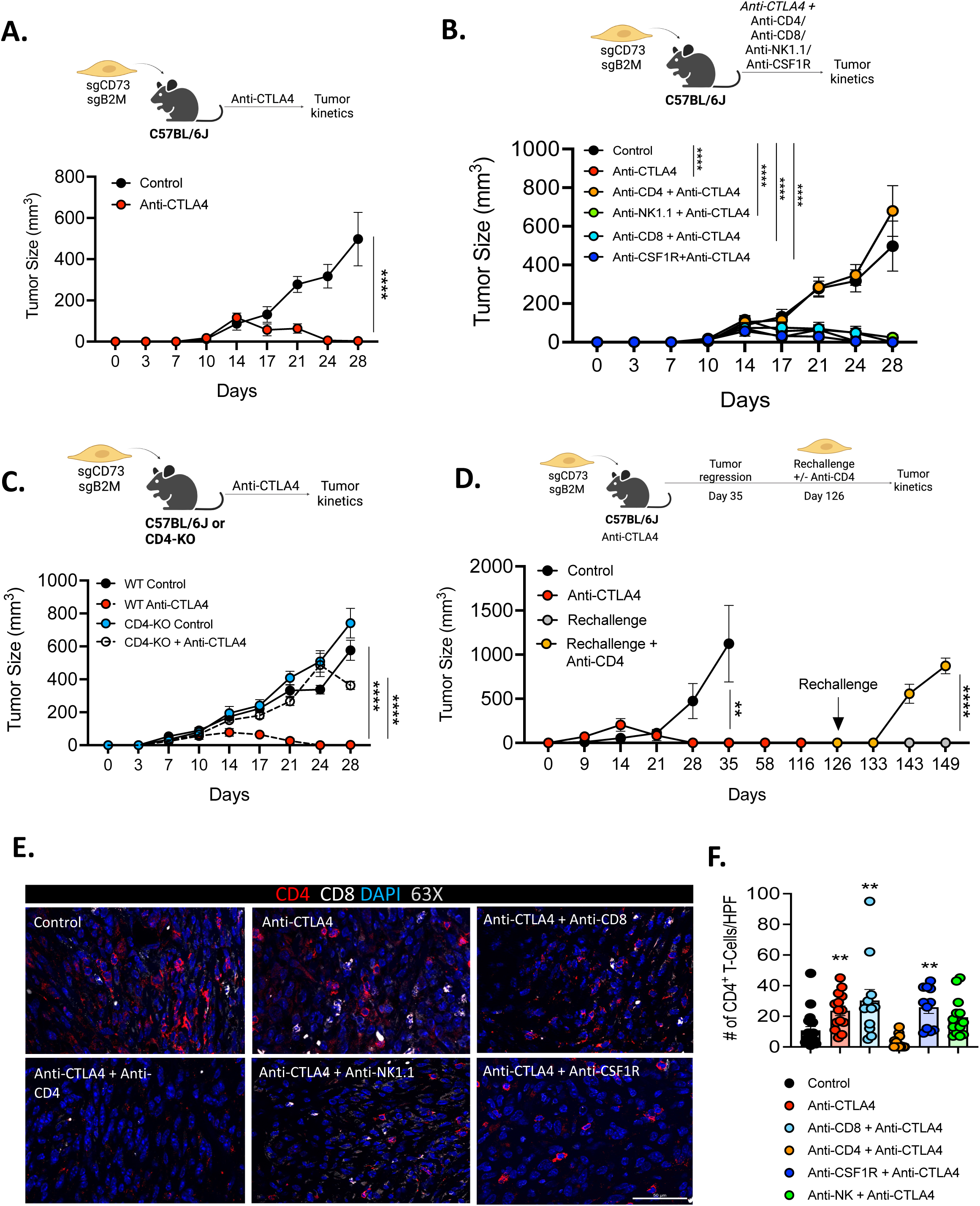
CD4^+^ T-cells sensitize qM tumors lacking CD73 and MHC-I to anti-CTLA4 ICB: **(A)** Schema and tumor kinetics for CD73 and B2M double knock-out (DKO) tumor-bearing mice treated with the indicated antibodies. Data represent 3 independent experiments where n=3-7 for each group **(B)** Schema and tumor kinetics for CD73 and B2M double knock-out (DKO) tumor-bearing mice treated with the indicated antibodies. Data represent 3 independent experiments where n=3-7 for each group. (**C**) Schema and tumor kinetics for CD73 and B2M double knock-out (DKO) tumors propagated in Wild Type (WT) or CD4 knock-out (CD4-KO) mice treated with the indicated antibodies. Data represent 3 independent experiments where n=3-7 for each group (**D**) Schema and tumor kinetics for CD73 and B2M double knock-out (DKO) tumor-bearing mice treated with the indicated antibodies. Responders were rechallenged with the same cell line as indicated with or without treatment with anti-CD4. Data represent 3 independent experiments where n=3-7 for each group. (**E, F**) Representative immune-fluorescence images of primary tumor sections obtained from (B) stained for CD4 (red), CD8 (white) and DAPI (blue) at 63X magnification. Bar graph on the right represents quantification of CD4^+^ T-cells in each high-power field (HPF) at 63X magnification for the indicated treatment groups. 3-5 fields of view from the tumor interior were obtained for each tumor. Data represent three independent experiments, with n=3-5 mice in each group. (**A-D**) Data represent SEM, two-way ANOVA, *\*\*, p<0.01, ***, p<0.001, ****, p<0.0001.* (**F**) Data represent SEM, two-tailed unpaired t-test, *\*, p<0.05, **, p<0.01, ***, p<0.001, ****, p<0.0001.* Scale bars are 100um.

To determine the functional importance of CD8^+^ T-cells, NK cells, or myeloid cells in driving this sensitization, we depleted each of these cells using subset-specific antibodies. Antibody-based depletion of each of these immune cells failed to reverse sensitization (Fig. 4B and fig. S5E). Moreover, responding tumor-bearing mice recruited elevated numbers of CD4^+^ T-cells to their tumors in response to anti-CTLA4 therapy even in the absence of CD8^+^ T-cells, NK-cells, and macrophages (Fig. 4E and F). In sharp contrast, antibody mediated depletion of CD4^+^ T-cells completely reversed the response of DKO tumor-bearing mice to anti-CTLA4 ICB (Fig. 4B and fig. S5E). Similarly, DKO cancer cells failed to respond to anti-CTLA4 ICB when implanted orthotopically in CD4-KO mice relative to WT mice (Fig. 4C and fig. S5F) once again, underscoring the functional importance of CD4^+^ T-cells in sensitizing qM breast tumors lacking CD73 to anti-CTLA4 ICB.

To determine the efficacy of the aforementioned complete responses, DKO tumor-bearing mice that had responded to anti-CTLA4 ICB were rechallenged with the same tumor cells. Strikingly, these tumor-bearing mice remained tumor-free upon rechallenge indicating the generation of anti-tumor memory (Fig. 4D). More importantly, these protective effects were lost when CD4^+^ T-cells were depleted prior to rechallenge. Thus, CD4^+^ T-cells not only drive the sensitization of qM tumors lacking CD73 to ICB, but are also required for long-term memory responses (Fig. 4D).

### EMP is associated with CD73 expression in human breast cancers

Our novel, preclinical murine models of epithelial and qM tumors have enabled us to identify the EMP program and cancer cell-intrinsic expression of CD73 as important determinants of responsiveness to anti-CTLA4 ICB therapy. To assess the translational potential of our findings, we first asked whether and which types of human breast cancers express CD73. To that end, we analyzed publicly available bulk and single cell RNA-seq datasets of different breast cancer cell lines (*22*). We observed that CD73 was expressed in a subtype specific manner with the highest levels of CD73 being expressed by triple negative breast cancers (TNBC) (Fig. 5A and fig. S6A). In sharp contrast, cell lines belonging to the luminal A/B subtypes expressed little to no CD73 (Fig. 5A and fig. S6B). Most strikingly, analysis of RNA-Seq transcriptomic datasets of breast cancer cell lines from the Cancer Cell Line Encyclopedia (CCLE) revealed that CD73 expression was correlated strongly with EMP as well as partial EMP pathways (*23*) (Fig. S6B-D). To further assess whether CD73 expression by more-mesenchymal cancer cells is specifically associated with luminal or basal characteristics, we projected transcriptomics data obtained from single cell RNA seq of breast cancer cell lines on a two dimensional epithelial-mesenchymal and luminal-basal plot (*22*). This analysis revealed that only cells that were enriched for a basal signature and were intermediate or high in mesenchymal signatures were more likely to express *NT5E* (the gene name for CD73) compared to luminal cells which were largely epithelial in nature (Fig. 5B). Taken together, these analyses suggest a strong correlation between CD73 expression and TNBCs that residence in a more-mesenchymal, basal-like state.

**Figure 5:**
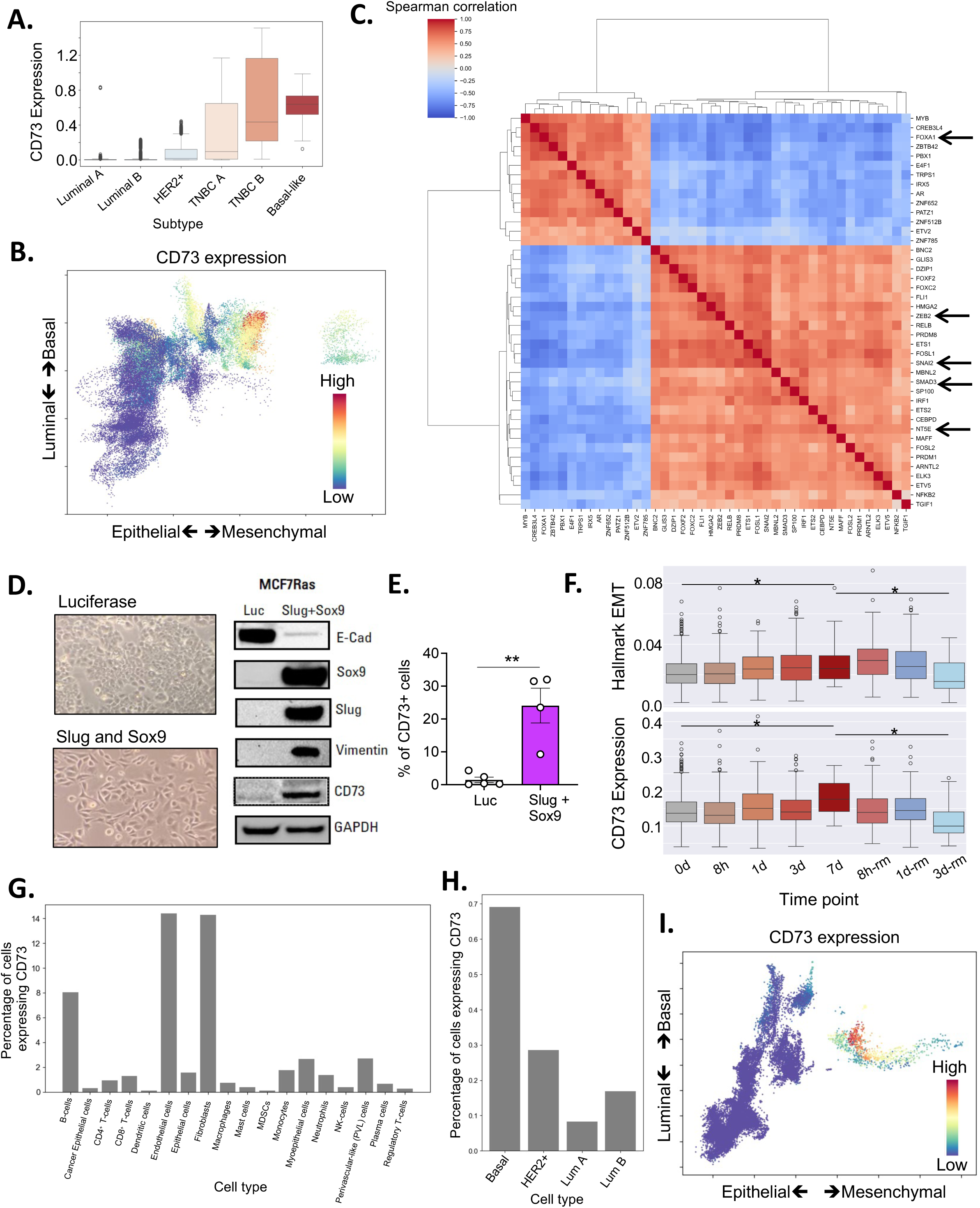
EMP regulates CD73 expression in human breast cancer: (**A**) Box plot showing the expression level of CD73 in breast cancer cell lines belonging to different breast cancer subtypes. (**B**) Scatter plot showing the relative position of single cells from different breast cancer cell lines on a two-dimensional luminal-basal and epithelial-mesenchymal plane. Individual cells are colored by CD73 expression levels with red denoting a higher expression. (**C**) A clustered heatmap showing top transcription factors that are correlated with CD73 expression in the CCLE breast cancer cell lines. (**D, E**) MCF7RAS cells expressing doxycycline-controlled control Luciferase or Slug and Sox9 co-expressing constructs were treated with doxycycline for 4 days to induce EMP. (**D**) Phase contrast images and western blots for the indicated markers. Data represent four independent experiments. (**E**) Bar graph representing percentage of cells expressing CD73. Data represent SEM, two-tailed unpaired t-test, *\*\*, p<0.01.* (**F**) Boxplot showing Hallmark EMT AUCell score and CD73 expression of MCF7 cells treated with TGF-ß followed by the removal of the same over a total period of 10 days. Students t-test was performed to assess the significance of difference of means in expression between day 0 with day 7 and day 7 with day 10 (3^rd^ day of TGF-ß removal). (**G**) Barplot showing the percentage of cells in which non-zero counts of reads were detected from single cell RNA seq of human breast cancer derived cells from different breast cancer subtypes. (**H**) Barplot showing the percentage of cells with non-zero counts of reads that were detected in tumor cells belonging to different breast cancer subtypes from human breast cancer samples. (**I**) Scatterplot showing the relative positions of tumor cells from human breast cancer samples on a two dimensional luminal-basal and epithelial-mesenchymal plane. Individual cells are colored by CD73 expression levels with red denoting a higher expression.

To understand the contribution of causal factors that could control cancer cell-intrinsic CD73 expression, we performed a correlation analysis of CD73 expression with all transcription factors using transcriptomic data of breast cancer cell lines from the CCLE. We observed that transcription factors expressed by luminal breast cancer cell lines, specifically PBX2 and FOXA1, were negatively correlated with *NT5E* expression. In sharp contrast, transcription factors that are known to activate EMP, specifically SNAI2(Slug), ZEB2, SMAD3 and FOXC2 were positively correlated with *NT5E* expression (Fig. 5C). To experimentally assess a causal connection between EMP and CD73 expression, we activated this program in MCF7RAS human breast cancer cells by doxycycline-controlled expression of two different EMT-inducing transcription factors SNAI2 and SOX9 (*20, 24*). Co-expression of these two EMT-TFs led to a robust activation of EMP as observed by the adoption of a more-mesenchymal morphology, loss of E-cadherin and gain of Vimentin relative to cells expressing a doxycycline-controlled luciferase construct. Most importantly, the acquisition of more-mesenchymal features was also accompanied by a concomitant increase in CD73 expression indicating a causal connection between EMP and CD73. (Fig. 5, D and E and fig S6E). To determine whether other EMT-TFs could also regulate the expression of CD73, we induced the expression of either ZEB1, TWIST or SLUG in MCF7RAS cells. The expression of these EMT-TFs resulted in only a partial-EMT as the cells retained varying levels of E-cadherin (fig S6F). However, residence in this partial state also resulted in increased CD73 expression albeit to a lesser extent compared to cells that co-expressed SLUG and SOX9 and underwent a more-complete transition (fig S6G).

To further assess causality, we analyzed single cell RNA-seq data from the MCF7 cell line that was induced to undergo an EMP by treatment with TGF-β for 7 days followed by a subsequent reversal of EMP by withdrawing TGF-β for 3 days (*25*). We observed that both, the Hallmark EMT signature as well as *NT5E* expression were significantly upregulated by day 7 of TGF-β treatment. Strikingly, the expression of both, EMT pathways and *NT5E* expression was reversibly lost as the signal for EMP was removed over the course of the next three days (Fig. 5F). Taken together, these findings establish the fact that the induction of cancer cell-intrinsic CD73 expression is EMP-dependent and that transcription factors can causally induce the expression of CD73 in a reversible manner.

Given the known immunosuppressive function of CD73, we sought to delineate its expression on various cell types within the tumor micro-environment of human breast tumors. Upon investigating a recently curated single cell RNA-seq atlas of human breast tumors (*26*), we found that several immune cells expressed varying levels of CD73 with the highest expression being present on B-cells, where it is known to generate adenosine, induce immunosuppressive effects and likely regulate class-switching (*27, 28*). Intriguingly, CD73 in the tumor microenvironment was also largely expressed by more-mesenchymal cells such as fibroblasts and endothelial cells, indicating a more general association of CD73 expression with the mesenchymal state even in non-cancer cells (Fig. 5G). Most importantly, within the epithelial, cancer-cell population, basal-like cancer cells had the highest expression of CD73 followed by HER2+ and Luminal B cancer cells (Fig. 5H). Finally, we analyzed tumor cells from a subset of breast cancer patients to determine which molecular phenotype of cancer cells express the highest levels of CD73 (*29*). Once again, we observed that patient derived breast cancer cells that expressed a basal and a partial EMP (pEMP)/full EMP phenotype were more likely to express CD73 in comparison to either luminal-epithelial or basal-epithelial cells (Fig. 5I). This was further exemplified by the fact that the same cells that expressed CD73 also expressed mesenchymal markers such as ZEB1, SLUG, FN1 and were low in CDH1 expression (Fig. S7A). In conclusion, analysis of multiple transcriptomic data sets of human breast cancer cell lines or scRNA-Seq data-sets obtained from breast cancer patients demonstrate a strong relationship between EMP and CD73 expression.

## Discussion

Epithelial-Mesenchymal Plasticity has long been studied as a process that potentiates metastasis and drives resistance of breast tumors to multiple forms of therapies, including targeted therapies, chemotherapies and more recently, immunotherapies (*6, 14, 30, 31*). Our previous and current findings have underscored the importance of the adenosine-generating ectoenzyme, CD73, in driving such resistance of quasi-mesenchymal breast tumors to anti-CTLA4 immune checkpoint blockade therapy(*20*). More specifically, inhibiting the expression of cancer cell-intrinsic CD73 can completely sensitize quasi-mesenchymal breast tumors to anti-CTLA4 ICB (*20*). While our previous work uncovered the importance of targeting the cancer cell-intrinsic adenosinergic pathway for eliminating more-mesenchymal cancer cells, the precise identity of immune cells that enabled such eradication remained unknown. In this study, we present findings that identify the functional importance of CD4^+^ T-cells in eliminating quasi-mesenchymal breast tumors lacking CD73 in response to anti-CTLA4 ICB.

We observed that the susceptibility of qM tumors lacking CD73 to anti-CTLA4 ICB was only partially dependent on CD8^+^ T-cells. Given the ability of more-mesenchymal cancer cells to downregulate MHC-I expression as a consequence of activating the EMT program, this finding is perhaps not that surprising (*18, 32*). Moreover, in previous studies we have determined that CD8^+^ T-cells infiltrating sgCD73 tumors do indeed, exhibit greater cytolytic function, relative to those present in control qM tumors even prior to administering ICB (*20*). However, despite retaining effector function, CD8^+^ T-cells were dispensable for eliminating sgCD73 tumors in response to anti-CTLA4 ICB. This is likely because CD8^+^ T-cells are outnumbered by their CD4^+^ T-cell counterparts in responding tumors as described in this study. Precisely why sgCD73 tumors recruit elevated numbers of CD4^+^ T-cells but not CD8^+^ T-cells in response to ICB treatment remains to be determined. Whether perturbation of CD73 in qM tumors promotes the release of cytokines and chemo-attractants that are specific for CD4^+^ T-cells is one possibility. Alternatively, anti-CTLA4 ICB treatment of sgCD73 tumors could result in more efficient priming and recruitment of peripheral CD4^+^ T-cells relative to CD8^+^ T-cells.

While CD4^+^ T-cells in the tumor microenvironment have primarily been studied as immunosuppressive Tregs, their anti-tumor functions in eliminating cancer cells, ostensibly akin to their CD8^+^ T-cell counterparts, is only beginning to emerge (*21, 33*). We observed marginally elevated numbers of T_H_1-like cells in sgCD73 tumors in response to anti-CTLA4 ICB treatment relative to those present in non-responding qM control tumors and no changes in other helper T-cell subsets. Previous studies have shown that both, anti-CTLA4 treatment or perturbation of CD73 signaling can specifically deplete Tregs and increase T_H_1-like cells (*34, 35*). The precise mechanism utilized by these CD4^+^ T-cells in eliminating more-mesenchymal breast cancer cells remains to be established. It is plausible that IFN-gamma derived from these T_H_1-like cells could exert anti-tumor effects ultimately leading to eradication of qM tumors.

We demonstrated that sgCD73 cancer cells lacking MHC-I were also sensitized to anti-CTLA4 ICB in a CD4^+^ T-cell dependent manner eliminating the contribution of NK-cells or myeloid cells in tumor clearance. Moreover, CD4^+^ T-cells were important not only for acute responses to anti-CTLA4 ICB, but also for long-term memory responses, underscoring their biological importance in controlling more-mesenchymal breast tumors. A few studies have implicated the ability of CD4^+^ T-cells to directly eliminate cancer cells that are MHC-II proficient or deficient by releasing pro-inflammatory cytokines (*36–38*). Other studies have identified alternative mechanisms where CD4^+^ T-cells enable tumoricidal myeloid cells to eliminate cancer cells regardless of MHC-II expression (*39, 40*). CD4^+^ T-cells can also eliminate cancer cells by increasing the cytolytic functions of CD8^+^ T-cells via dendritic cell licensing or via the formation of intra-tumoral triads (*41–43*). Determining precisely how CD4^+^ T-cells eliminate qM tumors lacking CD73 in response to anti-CTLA4 ICB will be critical for gleaning mechanistic insights for the results presented in this study.

By performing transcriptomic analyses of various published data sets from human breast tumors we observed that the expression of cancer cell-intrinsic CD73 was associated specifically with more basal-like, TNBCs. The ability of various EMT-TFs to induce the expression of CD73 also suggests a causal relationship between CD73 and the gain of more-mesenchymal properties by cancer cells. TNBCs activate components of the EMP program relative to other breast cancer subtypes and concomitantly resist ICB therapies (*44–46*). Given the ability of EMP to drive metastasis and resistance to therapies, our findings can spur translational efforts to (i) utilize the phenotypic plasticity of cancer cells along with CD73 expression and CD4^+^ T-cells as predictive criteria for ICB responsiveness and; (ii) therapeutically target these parameters in combination to potentiate the response of highly refractory TNBCs to anti-tumor immunity. Accordingly, such translational strategies hold the potential to be transformative for the treatment of human breast cancers.

We have previously demonstrated that qM tumors lacking CD73 are specifically, sensitized to anti-CTLA4 but not anti-PD1 ICB (*20*). The underlying reason(s) for this difference is unknown. Recognizing the importance of this observation, the goal of these studies was to first identify how disrupting CD73 signaling potentiates responses to anti-CTLA4 ICB and then apply the lessons learnt to anti-PD1 blockade in subsequent work. Several studies have outlined the ability of anti-PD1 blockade to reinvigorate stem-like CD8^+^ T-cell progenitors (*47, 48*). However, given the higher numbers and biological relevance of CD4^+^ T-cells, but not CD8^+^ T-cells in our models, this finding alone could explain the lack of response to anti-PD1 blockade. Future studies are aimed at understanding whether (and which) differential mechanisms are utilized by sgCD73 tumors in response to anti-CTLA4 versus anti-PD1 therapies. While 15-30% of breast cancer patients do respond to anti-PDL1 blockade, a vast majority of them remain unresponsive to single agent anti-CTLA4 or anti-CD73 ICB (*5, 49, 50*). Thus, the translational appeal of our work lies in determining mechanistically, how combining both agents together (anti-CTLA4 and anti-CD73 ICB) generates synergistic responses to specifically eliminate qM tumors. Indeed, the adenosine antagonist ciforadenant synergizes with anti-CTLA4 ICB in preclinical mouse models and is currently being used in phase 1b/2 clinical trials in combination with Ipilimumab (anti-CTLA4) or Nivolumab (anti-PD1) for metastatic renal cancer (*51*). We propose that similar strategies can also be utilized for the treatment of refractory, more-mesenchymal TNBC tumors.

## Materials and Methods Mice

C57BL/6J female mice, aged 6–8-weeks, were obtained from The Jackson Laboratory. Mice were age matched and randomly assigned to treatment or control groups in all experiments. All animal procedures were carried out in compliance with guidelines and protocols approved by the Institutional Animal Care and Use Committee (IACUC) and maintained by the Center for Animal Resources and Education (CARE) at Cornell University.

### Cell lines and tissue culture

All murine cell lines sgCD73, Snail^HI^ qM, CD73 and B2M double knock-out (DKO) and MCF7RAS human breast cancer cells containing doxycycline-inducible Luciferase or SLUG and SOX9 constructs were a kind gift from Weinberg Lab and established and maintained as previously described (*18–20*). Murine cell lines were cultured in a 1:1 mixture of Dulbecco’s modified Eagle’s medium (DMEM) and Ham’s F12 medium supplemented with 5% Bovine adult serum, 1x penicillin-streptomycin and 1x nonessential amino acids maintained at 37°C incubator containing 5% CO2. MCF7RAS cells were maintained in DMEM with 10% Bovine fetal serum and 1x penicillin-streptomycin as previously described (*18–20*). All cell lines were routinely tested for mycoplasma (from 2019-2025) using the MycoAlert Mycoplasma Detection Kit (Lonza). All cell lines are negative for Mycoplasma and not authenticated since they were first acquired.

### Generation of cell lines using CRISPR/Cas9

The sgCD73 cell line was established and maintained as previously described (*20*). In order to generate the CD73 and B2M double knock out (DKO) cell line, B2M was knocked out from the sgCD73 murine cell line via transient transfection using CRISPR/Cas9 plasmids obtained from Santa Cruz Biotechnology. Cells were initially seeded at 0.5 X 10^6^ cells/well in a 6-well plate for 12 hours and transfected with the plasmid of interest according to the manufacturer’s protocol. After 48 hours of incubation, cells expressing GFP were sorted using (BD Biosciences FACSMelody) into each well of a 96-well plate to obtain single cell clones. These single cell clones were then expanded and screened for presence or absence of B2M using Western blotting and for surface MHC-I by flow cytometry. B2M and MHC-I expression was measured on cell lines before and after treatment with 100ng/ml of IFN-gamma for 48 hrs.

### *In vivo* mouse models and tumor dissociation

For orthotopic tumor implantations, 1 × 10^6^ cells were counted and resuspended in 30 ul of media containing 20% Matrigel. Cells were then implanted orthotopically into the mammary fat pads of C57BL/6J mice. Tumor volume was calculated using the modified ellipsoid formula tumor volume (mm^3^) = *(L x W x W)/*2, where *L* represents the largest tumor diameter and *W* represents the perpendicular measurement. After the tumors reach 2000mm^3^, mice were sacrificed and tumors were collected. A small section of the tumors was saved for fixing while the rest was used for making single-cell suspensions for flow cytometry. For tumor digestions, tumors were cut and finely minced with a razor blade and digested in RPMI containing 2mg/ml Collagenase A (Krackeler Scientific) and 100 U/ml Hyaluronidase (Krackeler Scientific). The suspension was then incubated in a rotator at 37°C for 40 mins. After digestion, the single-cell suspension was filtered first through a 70 micron and then through a 40 micron pore sized strainer, followed by centrifugation at 1250rpm for 10 mins at 4°C. The pellets were then resuspended in RPMI containing Monensin GolgiStop (BD Sciences) and incubated for 3-5 hours at 37°C. After incubation, the cells were centrifuged at 1250rpm for 10 mins at 4°C and processed for flow cytometry as described below.

### Flow cytometry analysis

The following steps were performed in 96-well V-bottom microwell plates using single cell suspensions obtained after tumor processing. First, cells were centrifuged at 1250rpm for 5 mins at 4°C and the pellets were resuspended in 200ul of FACS wash buffer with a master mix containing surface markers each diluted 1:100: CD3 FITC (17A2; BioLegend), PD1 APC (J43, Invitrogen), CD45 BUV805 (30-F11, Invitrogen), CD4 eFluor 450 (RM4-5, Invitrogen), TIGIT PECy7 (1G9, BioLegend), CD25 BV650 (PC61.5, BioLegend), CD45 BUV563 (30-F11, BD Biosciences), CD3 PE (145-2c11, Invitrogen) for 30mins in dark on ice or at 4°C. Live/Dead staining was performed for 15-20 mins in dark on ice or at 4°C using the Invitrogen LIVE/DEAD Fixable Near-IR Dead Cell Stain Kit with APC-CY7 dye. Cells were then fixed for 30-60 mins using the Intracellular Fixation and Permeabilization Buffer Set (Thermo Fisher). Intracellular staining was performed for 30-60 mins in dark on ice or at 4°C using the FOXP3/Transcription Factor Staining Buffer Set (Thermo Fisher) for the following antibodies each diluted at 1:50 - FOXP3 FITC (FJK16s; BioLegend), CTLA4 PerCP-eFluor 710 (UC10-4 B9; BioLegend), TIM3 BV711 (RMT3-23; BioLegend), Ki-67 BUV737 (SolA15; Invitrogen), 41BB PE (TKS-1, BioLegend), RORgt PerCP-eFluor 710 (B2D, Invitrogen), IFN-gamma BV785(XMG1.2, BioLegend), IL-17a BUV395 (TC11-18H10, BD BioSciences) and T-BET PECY7 (4B10, BioLegend). The single stain controls were made using UltraComp eBeads Compensation Beads (Invitrogen). Flow cytometry data were acquired on a BD Biosciences FACSymphony A3, and data were analyzed using the FlowJo (TreeStar) software.

### Antibody depletion & Immune Checkpoint Blockade (ICB) treatment

For single or combinatorial treatment with immunotherapy, mice received 200 ug of anti-CTLA4 (clone 9H10, BioXCell) and/or 100 ug of anti-CD73 (clone TY/23, BioXCell) diluted in 200 ul of PBS administered intraperitoneally every other day for 6 doses followed by 100 ug weekly once injections for 7^th^ and 8^th^ dose. Other depletion antibodies were administered as follows: 200 ug anti-CD4 (clone GK1.5, BioXCell), 200ug anti-CD8a (clone 53-6.7, BioXCell), 200 ug of anti-NK1.1 (clone PK136, BioXCell) and 200 ug of anti-CSF1R (clone AFS98, BioXCell) administered once a week after 1-3 days of implantation until control tumors reached approximately 2000mm^3^ in size.

### Immunofluorescence staining

Tumor sections were fixed in 10% neutral buffered formalin for 24-48 hours and transferred to freshly made 70% ethanol, followed by embedding and sectioning. Formalin-fixed paraffin-embedded (FFPE) tumor sections were deparaffinized in Histoclear and rehydrated through a series of decreasing ethanol concentrations (100%, 95%, 75%, Milli-Q water and 1x DAKO wash buffer; 5 mins each). For antigen retrieval, the section slides were immersed in 1X DAKO antigen retrieval buffer pH 6.0 (Agilent Technologies) and microwaved. Following this, the slides were washed twice with 1x DAKO wash Buffer (Agilent Technologies). To block non-specific binding, sections were blocked with phosphate-buffered saline containing 0.3% TritonX 100 (Millipore Sigma) and 1% Normal donkey serum (Jackson Immunoresearch Lab) for 20 mins at Room temperature (RT). After blocking, the sections were stained overnight at 4°C with anti-Rabbit CD4 mAb (1:100 Abcam EPR19514) and anti-Mouse CD8a (1:100 Thermo Fisher). The sections were then washed twice with 1X DAKO wash buffer and tagged with Biotium CF555 Donkey anti-Rabbit IgG (1:500) and Biotium CF488A Donkey anti-Mouse IgG (1:500) for 2 hrs. at room temperature. After three washes, the sections were stained with DAPI (1:1000 from 10mg/ml stock-Millipore Sigma) for 5 mins at room temperature. Slides were washed for 5 mins in distilled water and mounted with Prolong Gold anti-Fade mounting media (Cell Signaling Technologies 9071S).

### Microscopy

The stained sections were viewed under a Zeiss AxioObserver inverted microscope with 63X oil objective and the images were analyzed using the ZEN imaging software.

### Western blots

Lysates were made in aqueous 1X RIPA lysis buffer (EMD Millipore Corp) and the concentration of extracted protein was measured using the Pierce BCA protein assay (Thermo Scientific, 23227) and the optical density was measured on a BioTek Synergy 2 Microplate Reader at 562nm and analyzed data on Gen5 1.11 software. 40ug of protein was resolved using the NuPAGE Bis-Tris mini protein gels, 12%,1.0mm (Invitrogen) and transferred to 0.2-micron immobilon-P PVDF membrane (Millipore Sigma) using a wet transfer. The blots were then blocked in 5% Blotto (Santa Cruz) containing 0.2% Tween-20 (Sigma Aldrich) in PBS. Membranes were then probed with primary antibodies (1:1000) overnight, washed and incubated with horseradish peroxidase-labeled secondary antibodies (1:5000) and developed using Dura substrate (Thermo Scientific). Primary antibodies: β2-microglobulin (Abcam), CD73 Monoclonal antibody (Invitrogen), GAPDH, E-Cadherin, Sox9, Slug, Vimentin (Cell Signaling Technologies) and secondary antibodies: anti-Rabbit IgG, HRP and anti-Mouse IgG, HRP (Cell Signaling Technologies)

### Processing tumors for single-cell RNA sequencing

Tumors were processed into a single cell suspension as described in the preceding sections and resuspended in 1X cold phosphate buffered saline. 2 million cells were then resuspended in 100ul of FACS wash buffer (0.05% bovine serum albumin in PBS). Each tumor sample was then incubated with the Total Seq antibody cocktail (Total Seq TM A0301 anti-mouse Hashtag 1 for Snail^HI^ qM tumors and Hashtag-2 for sgCD73 tumors; BioLegend Clone M1/42; 30-F11) at a final concentration of 10ug/ml. This was followed by a 30-minute incubation on ice, two washes with cold FACs buffer by centrifugation at 1250rpm for 10 mins, followed by resuspension in PBS for a final count of 1 million cells. Each sample was filtered through a 40-micron filter and pooled. Libraries were generated using the 10X Genomics Chromium Single Cell 3 prime library Gel Bead Kit V2 followed by purification using solid-phase reversible immobilization (SPRI) beads and sequencing on an Illumina NextSeq system. After sequencing, the first step in processing the raw data was the demultiplexing of the samples based on the unique hashtag antibodies using Cell Ranger. Quality control measures were implemented to focus on biologically relevant, high-quality single cells. We removed cells with fewer than 500 features (e.g., the number of genes detected in a cell) and cells with more than 7500 features, thereby excluding low-quality or non-viable cells and doublets. After quality control, the estimated number of labeled cells was 6332. Additionally, the percentage of mitochondrial gene expression in each cell was determined as part of the quality control process. We used Loupe to identify the number of cells from each cluster. LogNormalize was applied to normalize the feature expression measurements in Seurat. Subsequent cell clustering was performed using the Seurat 5.0.1 package in R, utilizing the Louvain algorithm based on the elbow method. We selected 10 principal components for the analysis, which led to the identification of eleven clusters representing B-cells, CD4^+^ T-cells, CD8^+^ T-cells, two subsets of macrophages, neutrophils, fibroblasts, endothelial cells, mesenchymal progenitor cells, and two subsets of cancer cells. Cluster identities were assigned based on known marker genes specific to each cell type. For each cell cluster, we identified 100 markers that defined the clusters through differential expression. A list of differentially expressed genes (DEGs) was compiled for each cluster, resulting in a heatmap that shows how the expression of specific clusters differed across samples (Fig 1B). For example, genes such as CD79a, CD79b, and CD19 showed high expression in cluster 0, suggesting that this cluster was associated with B-cells. Uniform Manifold Approximation and Projection (UMAP) was used to display the data.

### Computational analysis of breast cancer cell line and patient data

Pathway scores for all pathways were computed using AUCell scoring on different gene lists for different biological pathways considered (*52*). Spearman correlation was performed to assess the degree of correlation between CD73 expression and the different pathway scores calculated for the bulk RNA seq data from CCLE. Luminal and Basal gene expression signatures were obtained from (*53*). Epithelial and mesenchymal signatures were obtained from (*54*). Hallmark EMT signatures were obtained from MSigDB (*23*). pEMT signature was obtained from (*55*). The scores along the luminal-basal and epithelial-mesenchymal axis were calculated as described in (*53*).

### Statistical analysis

All statistics were performed using the GraphPad Prism V10 software. All data represent standard error of the mean (SEM) using either two-tailed unpaired t-tests (Fig 1E, G; Fig 2B, D, F, H; Fig 3F, G; Fig 4F) or a regular two-way ANOVA (Fig 2A, C, E, G; Fig 3A-E, Fig 4A-D). Asterisks indicate statistical significance where *\*, p<0.05, **, p<0.01, ***, p<0.001, ****, p<0.0001*.

## Acknowledgements

We thank all members of the Dongre Lab for helpful discussions and review of the manuscript. We thank Dr. Praveen Sethupathy, Dr. Paul Soloway, Dr. Andrew C. White, Dr. Glenn Simmons Jr., Dr. Barbara A. Osborne, Chia-Hsin Hsu, and Nicole Andre for critical reading of the manuscript. We are grateful for the Judith Appleton, PhD Early Career Excellence Award that has funded part of this work. We thank all staff in the Flow Cytometry Facility, Cornell Institute for Biotechnology, for assistance with flow cytometry. We thank all staff members at the Cornell Center for Animal Resources and Education (CARE) for mouse colony maintenance. We thank Scott Butler for assistance with mouse surgeries. We thank the Animal Health Diagnostic Center for assistance with processing slides for histology.

## Funding

A.D was supported by funds from K22 Transition Career Development Award, National Cancer Institute, National Institutes of Health (K22CA255420)

Breast Cancer Coalition of Rochester

Judith Appleton PhD Early Career Excellence in Research Award

MKJ was supported by Param Hansa Philanthropies.

SS was supported by Prime Ministers’ Research Fellowship (PMRF), Government of India.

## Author Contributions

Conceptualization: SC, AD

Methodology: SC, CS, SS, BF, IOC, LL, SN, SP, KB, PT, GWB, CS, MKJ, AD

Investigation: SC, CS, SS, BF, IOC, LL, SN, SP, KB, PT, GWB, CS, MKJ, AD

Visualization: SC, AD Supervision: MKJ, AD

Writing—original draft: SC, AD

Writing—review & editing: SC, SS, CS, MKJ, AD

## Data and Materials Availability

All data are made available in the manuscript figures and supplementary figures

## References

1. P. Sharma, J. P. Allison, The future of immune checkpoint therapy. Science 348, 56–61 (2015).

2. I. O’Connell, A. Dongre, Immune Checkpoint Blockade Therapy for Breast Cancer: Lessons from Epithelial-Mesenchymal Transition. Mol Diagn Ther 27, 433–444 (2023).

3. J. Larkin, F. S. Hodi, J. D. Wolchok, Combined Nivolumab and Ipilimumab or Monotherapy in Untreated Melanoma. N Engl J Med 373, 1270–1271 (2015).

4. W. Hugo et al., Genomic and Transcriptomic Features of Response to Anti-PD-1 Therapy in Metastatic Melanoma. Cell 165, 35–44 (2016).

5. L. A. Emens et al., First-line atezolizumab plus nab-paclitaxel for unresectable, locally advanced, or metastatic triple-negative breast cancer: IMpassion130 final overall survival analysis. Ann Oncol 32, 983–993 (2021).

6. S. Lamouille, J. Xu, R. Derynck, Molecular mechanisms of epithelial-mesenchymal transition. Nat Rev Mol Cell Biol 15, 178–196 (2014).

7. S. A. Mani et al., The epithelial-mesenchymal transition generates cells with properties of stem cells. Cell 133, 704–715 (2008).

8. A. P. Morel et al., Generation of breast cancer stem cells through epithelial-mesenchymal transition. PLoS One 3, e2888 (2008).

9. A. Singh, J. Settleman, EMT, cancer stem cells and drug resistance: an emerging axis of evil in the war on cancer. Oncogene 29, 4741–4751 (2010).

10. I. Pastushenko et al., Identification of the tumour transition states occurring during EMT. Nature 556, 463–468 (2018).

11. B. Bierie et al., Integrin-beta4 identifies cancer stem cell-enriched populations of partially mesenchymal carcinoma cells. Proc Natl Acad Sci U S A 114, E2337–E2346 (2017).

12. J. Yang et al., Guidelines and definitions for research on epithelial-mesenchymal transition. Nat Rev Mol Cell Biol 21, 341–352 (2020).

13. D. R. Pattabiraman et al., Activation of PKA leads to mesenchymal-to-epithelial transition and loss of tumor-initiating ability. Science 351, aad3680 (2016).

14. A. Dongre, R. A. Weinberg, New insights into the mechanisms of epithelial-mesenchymal transition and implications for cancer. Nat Rev Mol Cell Biol 20, 69–84 (2019).

15. M. Plaschka et al., ZEB1 transcription factor promotes immune escape in melanoma. J Immunother Cancer 10, (2022).

16. C. Kudo-Saito, H. Shirako, T. Takeuchi, Y. Kawakami, Cancer Metastasis Is Accelerated through Immunosuppression during Snail-induced EMT of Cancer Cells. Cancer Cell 15, 195–206 (2009).

17. M. P. Mak et al., A Patient-Derived, Pan-Cancer EMT Signature Identifies Global Molecular Alterations and Immune Target Enrichment Following Epithelial-to-Mesenchymal Transition. Clin Cancer Res 22, 609–620 (2016).

18. A. Dongre et al., Epithelial-to-Mesenchymal Transition Contributes to Immunosuppression in Breast Carcinomas. Cancer Res 77, 3982–3989 (2017).

19. X. Ye et al., Distinct EMT programs control normal mammary stem cells and tumour-initiating cells. Nature 525, 256-+ (2015).

20. A. Dongre et al., Direct and Indirect Regulators of Epithelial-Mesenchymal Transition-Mediated Immunosuppression in Breast Carcinomas. Cancer Discov 11, 1286–1305 (2021).

21. D. E. Speiser, O. Chijioke, K. Schaeuble, C. Munz, CD4(+) T cells in cancer. Nat Cancer 4, 317–329 (2023).

22. G. Gambardella et al., A single-cell analysis of breast cancer cell lines to study tumour heterogeneity and drug response. Nat Commun 13, 1714 (2022).

23. A. Liberzon et al., Molecular signatures database (MSigDB) 3.0. Bioinformatics 27, 1739–1740 (2011).

24. W. Guo et al., Slug and Sox9 cooperatively determine the mammary stem cell state. Cell 148, 1015–1028 (2012).

25. D. P. Cook, B. C. Vanderhyden, Context specificity of the EMT transcriptional response. Nat Commun 11, 2142 (2020).

26. L. Xu et al., A comprehensive single-cell breast tumor atlas defines epithelial and immune heterogeneity and interactions predicting anti-PD-1 therapy response. Cell Rep Med 5, 101511 (2024).

27. F. Schena et al., Dependence of immunoglobulin class switch recombination in B cells on vesicular release of ATP and CD73 ectonucleotidase activity. Cell Rep 3, 1824–1831 (2013).

28. R. A. Miller et al., Anti-CD73 antibody activates human B cells, enhances humoral responses and induces redistribution of B cells in patients with cancer. J Immunother Cancer 10, (2022).

29. S. Z. Wu et al., A single-cell and spatially resolved atlas of human breast cancers. Nat Genet 53, 1334–1347 (2021).

30. R. D. Z. Mullins, A. Pal, T. F. Barrett, M. E. Heft Neal, S. V. Puram, Epithelial-Mesenchymal Plasticity in Tumor Immune Evasion. Cancer Res 82, 2329–2343 (2022).

31. S. Terry et al., New insights into the role of EMT in tumor immune escape. Mol Oncol 11, 824–846 (2017).

32. I. Akalay et al., EMT impairs breast carcinoma cell susceptibility to CTL-mediated lysis through autophagy induction. Autophagy 9, 1104–1106 (2013).

33. J. Borst, T. Ahrends, N. Babala, C. J. M. Melief, W. Kastenmuller, CD4(+) T cell help in cancer immunology and immunotherapy. Nat Rev Immunol 18, 635–647 (2018).

34. S. C. Wei et al., Distinct Cellular Mechanisms Underlie Anti-CTLA-4 and Anti-PD-1 Checkpoint Blockade. Cell 170, 1120–1133 e1117 (2017).

35. S. C. Wei et al., Combination anti-CTLA-4 plus anti-PD-1 checkpoint blockade utilizes cellular mechanisms partially distinct from monotherapies. Proc Natl Acad Sci U S A 116, 22699–22709 (2019).

36. J. Matsuzaki et al., Nonclassical antigen-processing pathways are required for MHC class II-restricted direct tumor recognition by NY-ESO-1-specific CD4(+) T cells. Cancer Immunol Res 2, 341–350 (2014).

37. A. Cachot et al., Tumor-specific cytolytic CD4 T cells mediate immunity against human cancer. Sci Adv 7, (2021).

38. D. Mumberg et al., CD4(+) T cells eliminate MHC class II-negative cancer cells in vivo by indirect effects of IFN-gamma. Proc Natl Acad Sci U S A 96, 8633–8638 (1999).

39. B. Kruse et al., CD4(+) T cell-induced inflammatory cell death controls immune-evasive tumours. Nature 618, 1033–1040 (2023).

40 E. G. Bawden et al., CD4(+) T cell immunity against cutaneous melanoma encompasses multifaceted MHC II-dependent responses. Sci Immunol 9, eadi9517 (2024).

41. A. Magen et al., Intratumoral dendritic cell-CD4(+) T helper cell niches enable CD8(+) T cell differentiation following PD-1 blockade in hepatocellular carcinoma. Nat Med 29, 1389–1399 (2023).

42. G. Espinosa-Carrasco et al., Intratumoral immune triads are required for adoptive T cell therapy-mediated elimination of solid tumors. bioRxiv, (2023).

43. M. Binnewies et al., Unleashing Type-2 Dendritic Cells to Drive Protective Antitumor CD4(+) T Cell Immunity. Cell 177, 556–571 e516 (2019).

44. M. Segovia-Mendoza, S. Romero-Garcia, C. Lemini, H. Prado-Garcia, Determining Factors in the Therapeutic Success of Checkpoint Immunotherapies against PD-L1 in Breast Cancer: A Focus on Epithelial-Mesenchymal Transition Activation. J Immunol Res 2021, 6668573 (2021).

45. L. A. Emens, Immunotherapy in Triple-Negative Breast Cancer. Cancer J 27, 59–66 (2021).

46. Y. K. Bae, J. E. Choi, S. H. Kang, S. J. Lee, Epithelial-Mesenchymal Transition Phenotype Is Associated with Clinicopathological Factors That Indicate Aggressive Biological Behavior and Poor Clinical Outcomes in Invasive Breast Cancer. J Breast Cancer 18, 256–263 (2015).

47. I. Siddiqui et al., Intratumoral Tcf1(+)PD-1(+)CD8(+) T Cells with Stem-like Properties Promote Tumor Control in Response to Vaccination and Checkpoint Blockade Immunotherapy. Immunity 50, 195–211 e110 (2019).

48. B. C. Miller et al., Subsets of exhausted CD8(+) T cells differentially mediate tumor control and respond to checkpoint blockade. Nat Immunol 20, 326–336 (2019).

49. L. A. Emens et al., Long-term Clinical Outcomes and Biomarker Analyses of Atezolizumab Therapy for Patients With Metastatic Triple-Negative Breast Cancer: A Phase 1 Study. JAMA Oncol 5, 74–82 (2019).

50. T. E. Keenan, S. M. Tolaney, Role of Immunotherapy in Triple-Negative Breast Cancer. J Natl Compr Canc Netw 18, 479–489 (2020).

51. S. B. Willingham et al., A2AR Antagonism with CPI-444 Induces Antitumor Responses and Augments Efficacy to Anti-PD-(L)1 and Anti-CTLA-4 in Preclinical Models. Cancer Immunol Res 6, 1136–1149 (2018).

52. S. Aibar et al., SCENIC: single-cell regulatory network inference and clustering. Nat Methods 14, 1083–1086 (2017).

53. S. Sahoo et al., Increased prevalence of hybrid epithelial/mesenchymal state and enhanced phenotypic heterogeneity in basal breast cancer. iScience 27, 110116 (2024).

54. T. Z. Tan et al., Epithelial-mesenchymal transition spectrum quantification and its efficacy in deciphering survival and drug responses of cancer patients. EMBO Mol Med 6, 1279–1293 (2014).

55. S. V. Puram et al., Single-Cell Transcriptomic Analysis of Primary and Metastatic Tumor Ecosystems in Head and Neck Cancer. Cell 171, 1611–1624 e1624 (2017).

