## Supplementary Information for "CD4^+^ T-cells sensitize quasi-mesenchymal breast tumors lacking CD73 to anti-CTLA4 immune checkpoint blockade therapy"

### Supplementary Fig 1

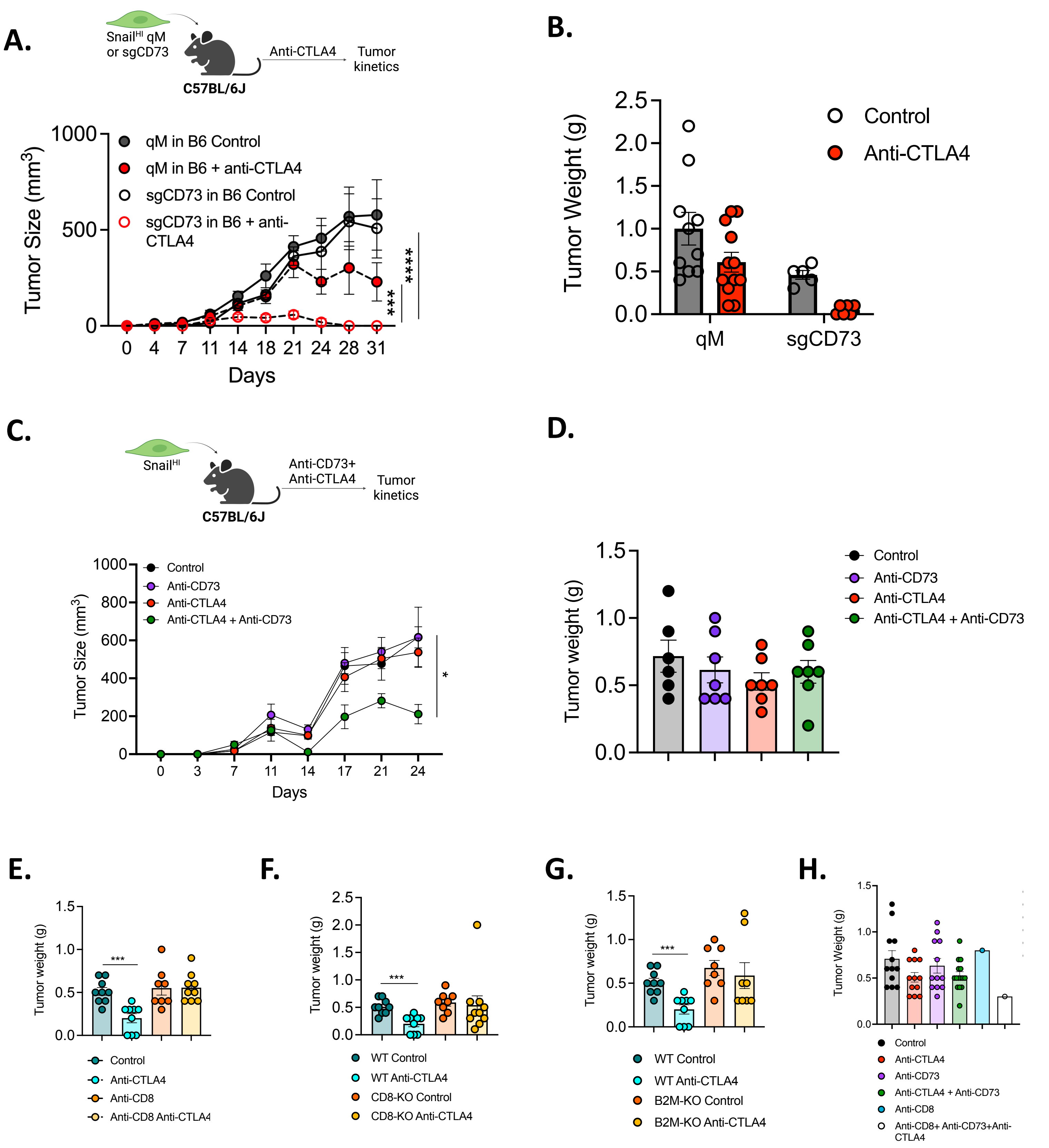

### Supplementary Fig 2

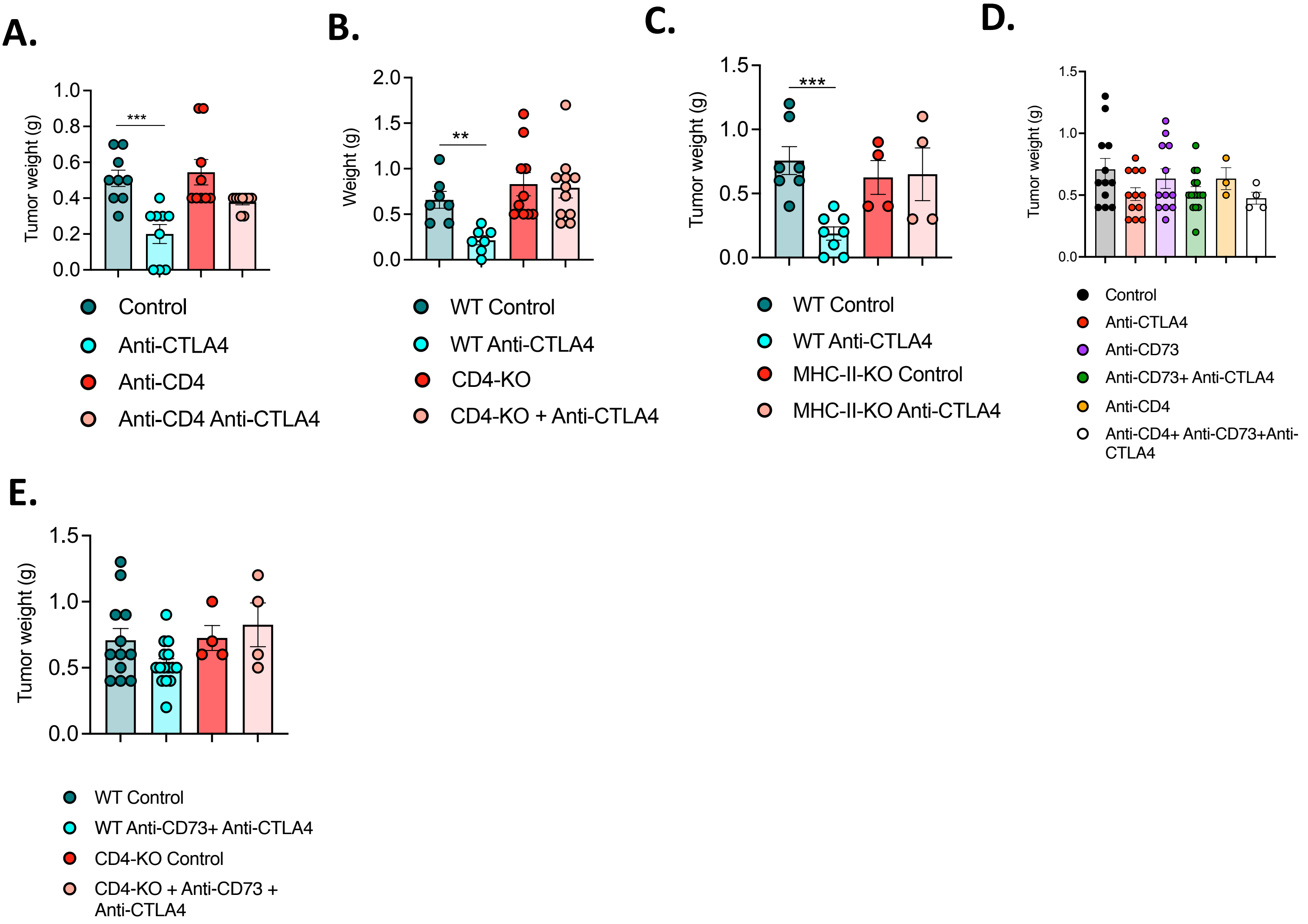

Supplementary Fig 3

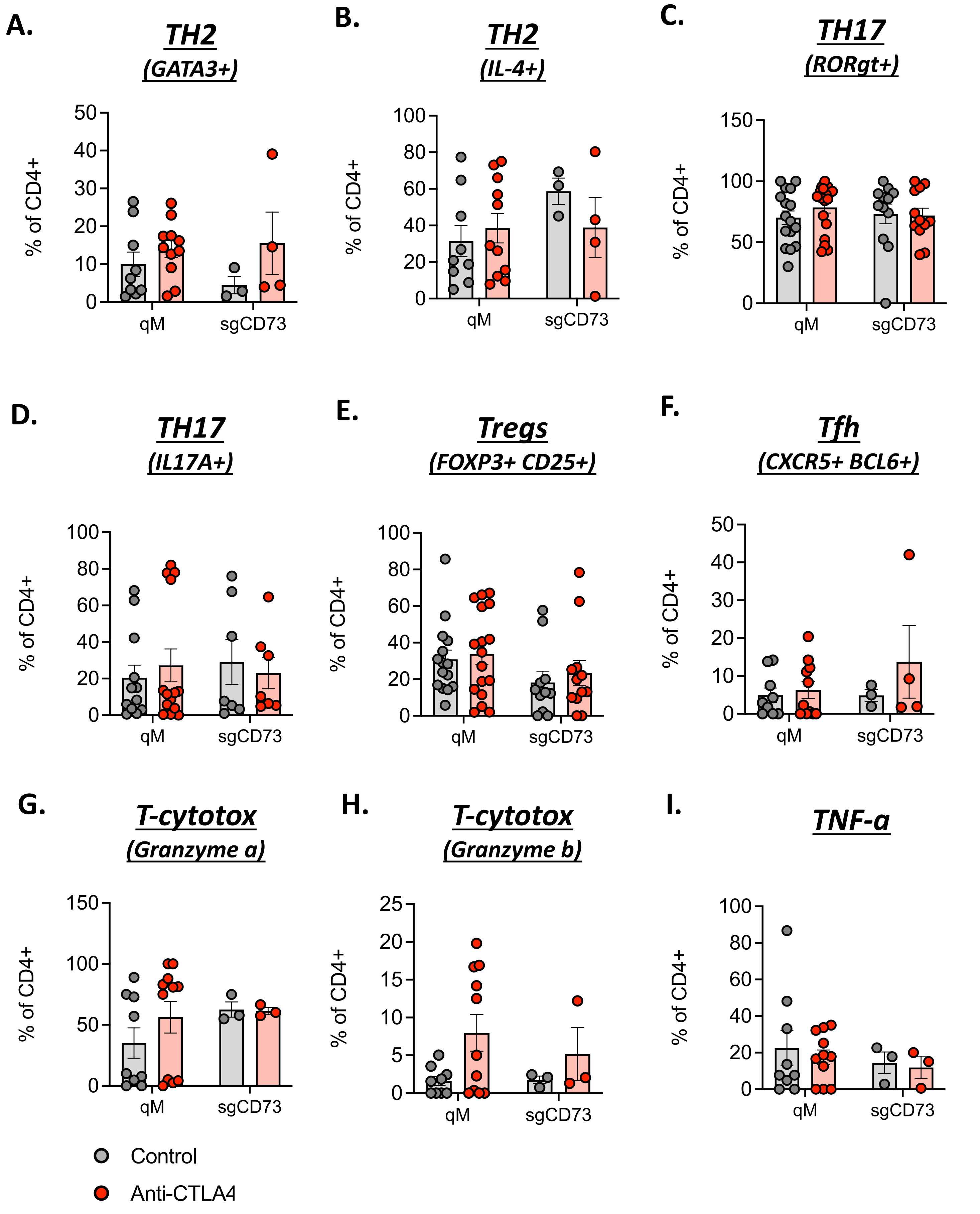

#### Supplementary Fig 4

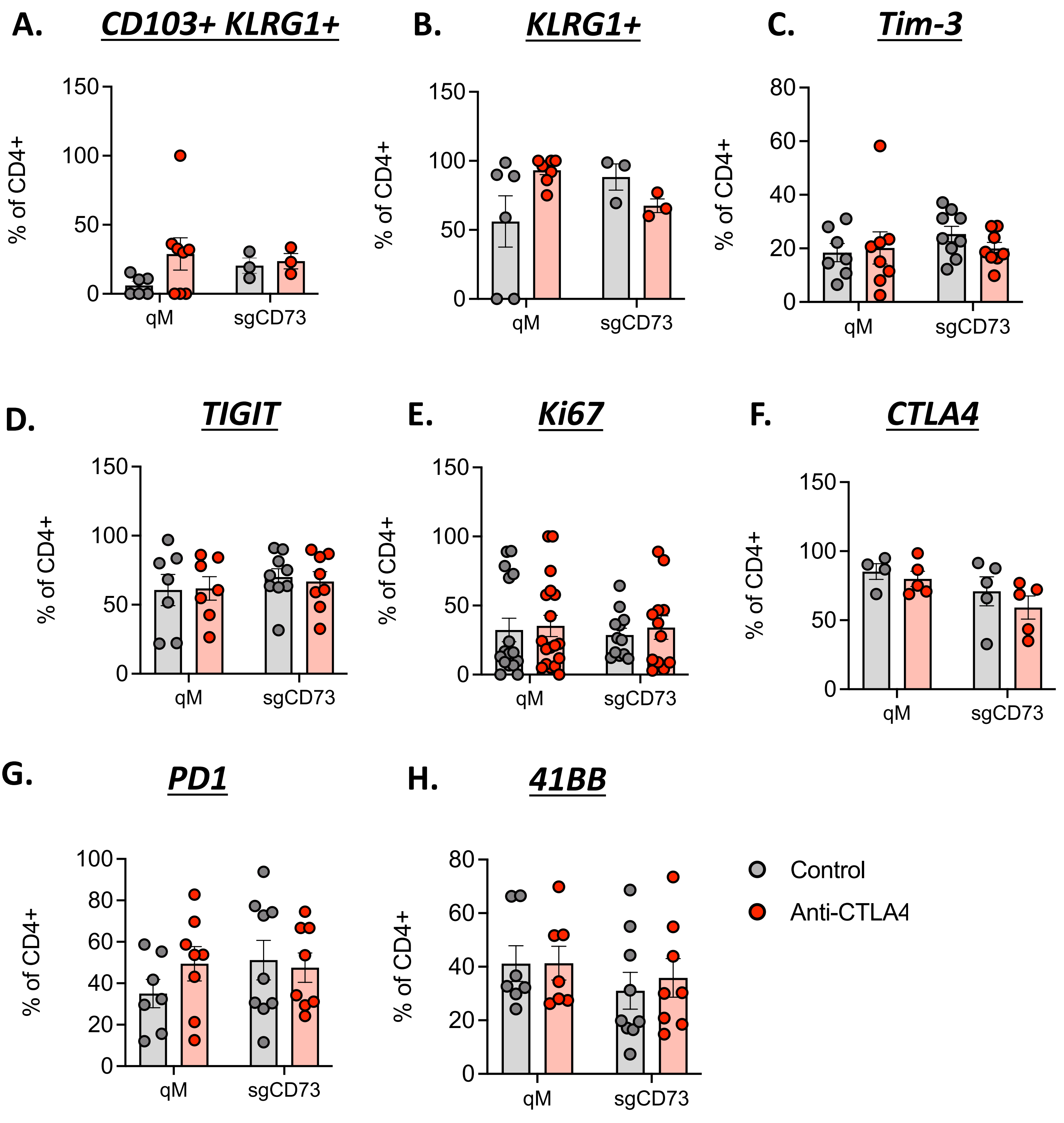

### Supplementary Fig 5

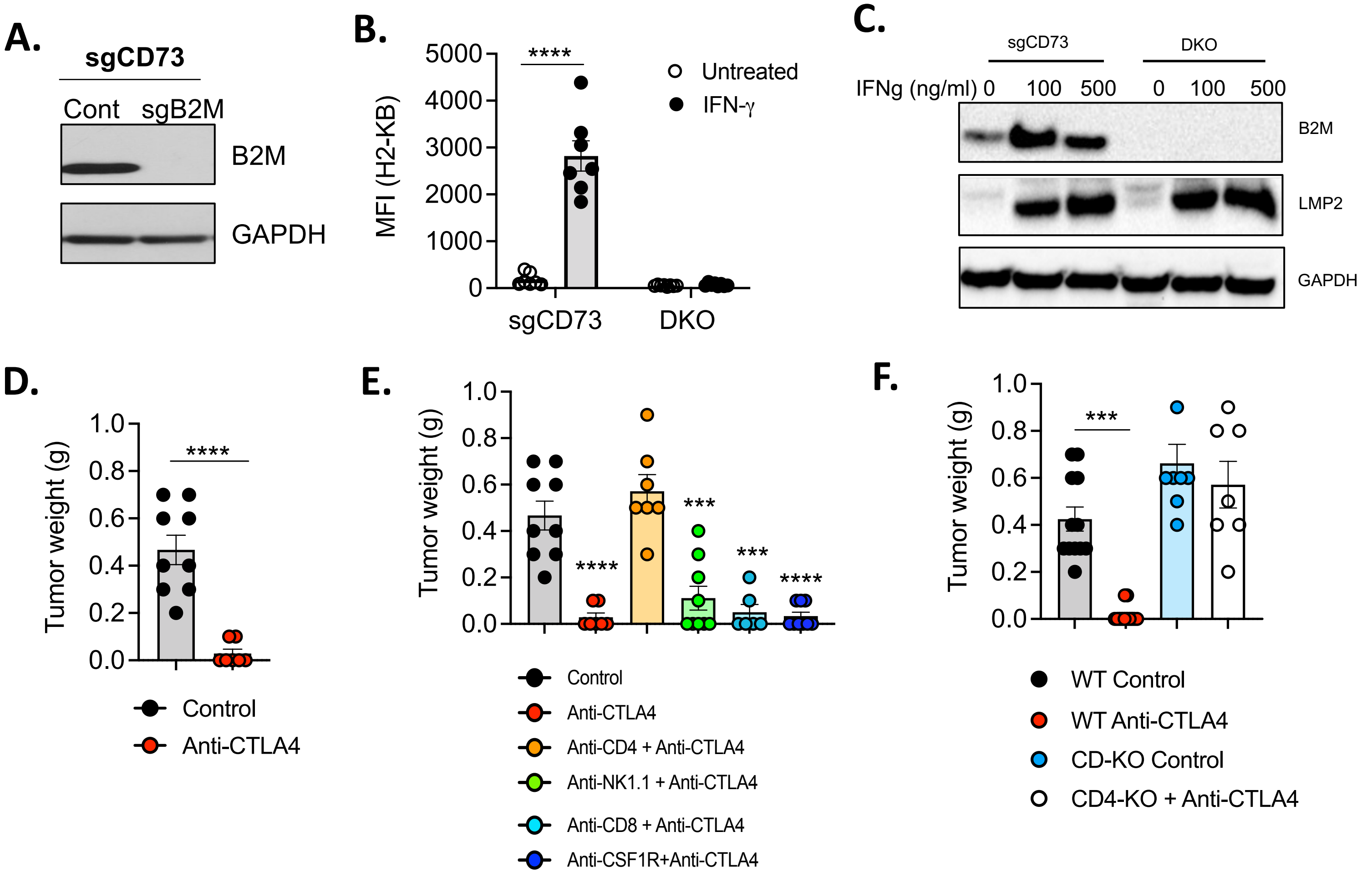

#### Supplementary Fig 6

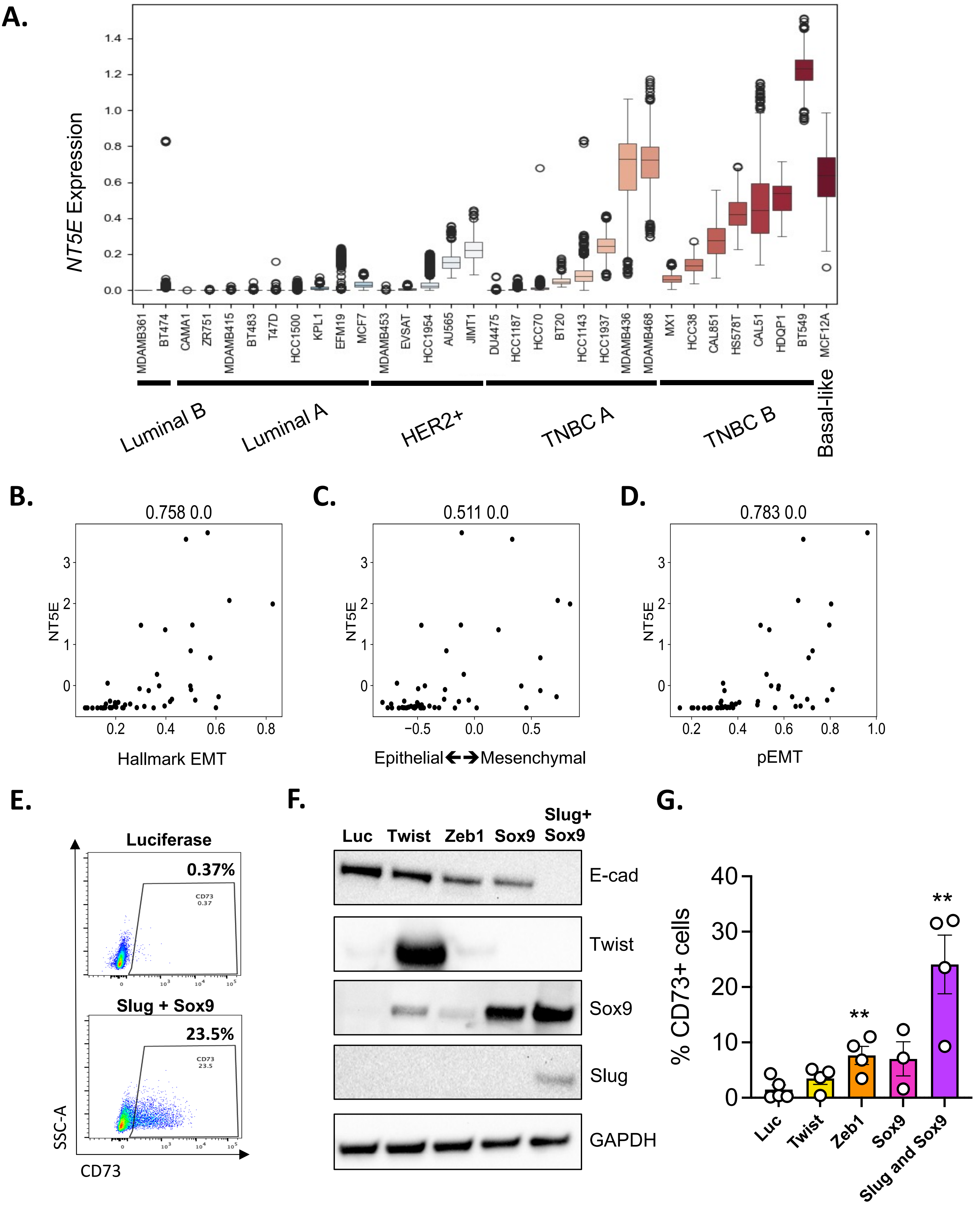

### Supplementary Fig 7

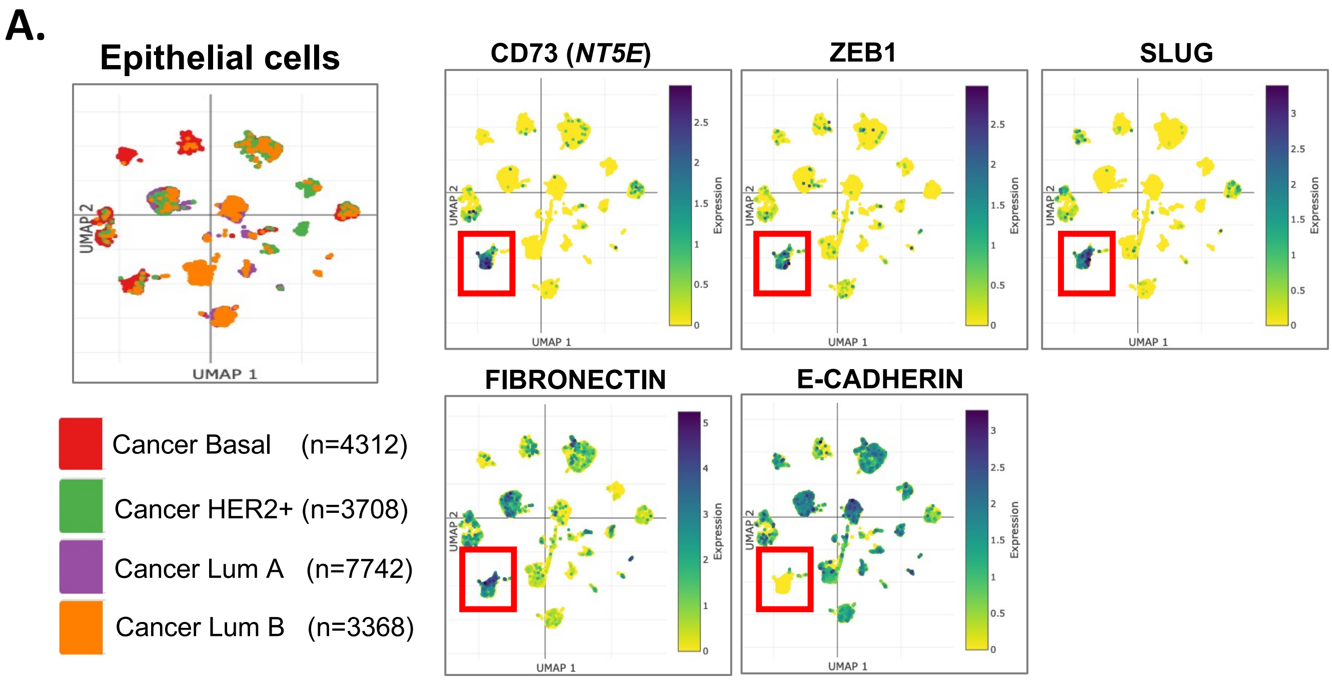

**Supplementary Figure Legends**

***Supplementary Figure 1*: Targeting CD73 sensitizes quasi-mesenchymal tumors to anti-CTLA4 immune checkpoint blockade therapy (A)** Schema and tumor kinetics, and (**B**) tumor weights for Snail$^{HI}$ qM and sgCD73 tumor-bearing mice receiving Control or anti-CTLA4 antibodies. Data represent three independent experiments where n=3-5 for each group. (**C**) Schema and tumor kinetics, and (**D**) tumor weights obtained from Snail$^{HI}$ qM tumor-bearing mice receiving Control, anti-CD73, anti-CTLA4 or combinations of anti-CD73 and anti-CTLA4 antibodies. Data represent three independent experiments with n=2-3 mice in each group. (**E**) Tumor weights for sgCD73 tumor-bearing mice treated with the indicated antibodies. Data represent three independent experiments where n=2-4 for each group. (**F**) Tumor weights for sgCD73 tumors propagated in Wild Type (WT) or CD8 knock-out (CD8-KO) mice treated with the indicated antibodies. Data represent four independent experiments where n=2-4 for each group (**G**) Tumor weights for sgCD73 tumors propagated in Wild Type (WT) or B2M knock-out (B2M-KO) mice treated with the indicated antibodies. Data represent three independent experiments where n=3-5 for each group. (**H**) Tumor weights for Snail$^{HI}$ qM tumor-bearing mice treated with the indicated antibodies. Data represent three independent experiments where n=3-5 for each group. (**A, C**) Data represent SEM, two-way ANOVA, *, p<0.05, ****, p<0.0001. (**B, D, E, F, G, H**) Bar graph data represent SEM, two-tailed unpaired t-test, ***, p<0.001, ns= *non-significant.*

***Supplementary Figure 2*: Targeting CD73 sensitizes quasi-mesenchymal tumors to anti-CTLA4 immune checkpoint blockade therapy in a CD4$^{+}$ T-cell dependent manner** (**A**) Tumor weights for sgCD73 tumor-bearing mice treated with the indicated antibodies. Data represent three independent experiments where n=3-4 for each group. (**B**) Tumor weights for sgCD73 tumors propagated in Wild Type (WT) or CD4 knock-out (CD4-KO) mice treated with the indicated antibodies. Data represent four independent experiments where n=2-4 for each group (**C**) Tumor weights for sgCD73 tumors propagated in Wild Type (WT) or MHC-II knock-out (MHC-II-KO) mice treated with the indicated antibodies. Data represent three independent experiments where n=3-7 for each group. (**D**) Tumor weights for Snail$^{HI}$ qM tumor-bearing mice treated with the indicated antibodies. Data represent three independent experiments where n=3-4 for each group (**E**) Tumor weights for Snail$^{HI}$ qM tumors propagated in Wild Type (WT) or CD4 knock-out (CD4-

KO) mice treated with the indicated antibodies. Data represent three independent experiments where n=2-4 for each group. (**A**, **B, C, D, E**) Bar graph data represent SEM, two-tailed unpaired t-test, **, $p<0.01$, ***, $p<0.001$, ns= *non-significant.*

***Supplementary Figure 3*****: Presence of $CD4^+$ T-cell helper subsets in responders and non-responders.** Snail$^{HI}$ qM and sgCD73 tumors that were untreated or treated with anti-CTLA4 were analyzed by flow cytometry for the indicated immune cell populations after gating on $CD45^+$ cells followed by $CD3^+$ T-cells. (**A**) TH2, % of $CD4^+$ T-cell expressing GATA3. (**B**) TH2, % of $CD4^+$ T-cell expressing IL-4. (**C**) TH17, % of $CD4^+$ T-cell expressing RORgt. (**D**) TH17, % of $CD4^+$ T-cell expressing IL-17A. (**E**) Tregs, % of $CD4^+$ T-cell expressing FOXP3 and CD25. (**F**) Tfh, % of $CD4^+$ T-cell expressing CXCR5 and BCL6. (**G**) T-cytotox, % of $CD4^+$ T-cell expressing Granzyme A. (**H**) T-cytotox, % of $CD4^+$ T-cell expressing Granzyme B. (**I**) TNF-alpha, % of $CD4^+$ T-cell expressing TNF-alpha. Bar graph data represent SEM, One-way ANOVA with Sidak's multiple comparison test. All comparisons are not significant.

***Supplementary Figure 4:*** **Activation and exhaustion markers on $CD4^+$ T-cell subsets present in responders and non-responders.** Snail$^{HI}$ qM and sgCD73 tumors that were untreated or treated with anti-CTLA4 were analyzed by flow cytometry for the indicated immune cell populations after gating on $CD45^+$ cells followed by $CD3^+$ T-cells. (**A**) CD103 and KLRG1, % of $CD4^+$ T-cell expressing CD103 and KLRG1. (**B**) KLRG1, % of $CD4^+$ T-cell expressing KLRG1. (**C**) Tim3, % of $CD4^+$ T-cell expressing Tim3. (**D**) TIGIT, % of $CD4^+$ T-cell expressing TIGIT. (**E**) Ki67, % of $CD4^+$ T-cell expressing Ki67. (**F**) CTLA4, % of $CD4^+$ T-cell expressing CTLA4. (**G**) PD1, % of $CD4^+$ T-cell expressing PD1. (**H**) 41BB, % of $CD4^+$ T-cell expressing 41BB. Bar graph data represent SEM, One-way ANOVA with Sidak's multiple comparison test. All comparisons are not significant.

***Supplementary Figure 5*****: sgCD73 cells lacking MHC-I respond to anti-CTLA4 immune checkpoint blockade therapy in a $CD4^+$ T-cell dependent manner** (**A**) Western blot analysis for control CD73 and CD73 and B2M double knock-out (DKO) cell lines for the indicated markers. Data represent 3 three independent experiments **(B)** Mean fluorescent intensity (MFI) for MHCI (H2-KB) as determined by flow cytometry for the indicated cell lines that were either untreated or treated with IFN-gamma for 48 hours. Data represent from pooled values from three independent experiments. (**C**) Western Blot analysis for the indicated cell lines that were either untreated or treated with IFN-gamma for 48 hours. Blots were probed for the indicated markers. Data represents three independent experiments. (**D**)Tumor weights for CD73 and B2M double knock-out (DKO) tumor-bearing mice receiving Control or anti-CTLA4 antibodies. Data represent four independent experiments where n=3-5 for each group. (**E**) Tumor weights for CD73 and B2M double knock-out (DKO) tumor-bearing mice treated with the indicated antibodies. Data represent four independent experiments where n=3-5 for each group. (**F**) Tumor weights for CD73 and B2M double knock-out (DKO) tumors propagated in Wild Type (WT) or CD4 knock-out (CD4-KO) mice treated with the indicated antibodies. Data represent 3 independent experiments where n=2-5 for each group. (**B**) Data represent SEM, two-way ANOVA, ****, p<0.0001. (**D, E, F**) Data represent SEM, two-tailed unpaired t-test, ***, p<0.001, ****, p<0.0001.

***Supplementary Figure 6*: EMP regulates CD73 expression on human breast cancer cell lines**
(**A**) Expression of CD73 (*NT5E*) from transcriptomic data obtained from the CCLE for the indicated breast cancer cell lines. (**B-D**) Data from panel A represented as correlation plots between CD73 (*NT5E*) against (**B**) ssGSEA based Hallmark EMT score; (**C**) ssGSEA based Mesenchymal score – Epithelial score (higher values indicate a higher mesenchymal nature); (**D**) ssGSEA based pEMT score. Numbers on top of each panel represent the correlation co-efficient and p-value (**E-G**) MCF7RAS cells expressing doxycycline-controlled control Luciferase or the indicated EMT-TFs were treated with doxycycline for 4 days to induce EMP. (**E**) Flow cytometry plots for the indicated cell lines representing the % of cells expressing CD73. Data represent four independent experiments. (**F**)Western blot for the indicated markers. Data represent four independent experiments. (**G**) Bar graph representing percentage of cells expressing CD73. Data represent four independent experiments. Data represent SEM, two-tailed unpaired t-test, ***, p<0.01.*

***Supplementary Figure 7*: CD73 expression on human breast cancer patient samples**
(**A**) scRNA-Seq transcriptomic data of human breast tumors obtained from ref 28 were analyzed in the single cell portal after specifically selecting for epithelial cancer cells belonging to the basal,

HER2+, Luminal A or Luminal B subsets where n represents the number of cells in each subset. UMAP plots for the indicated markers are represented for each of these subsets.

**Supplementary Table 1: Genes present in each cluster depicted in Figure 1A**

**Cluster 0 B cells**

```
cluster.markers <- FindMarkers(danx, ident.1 =0, min.pct = 0.25)
  |++++++++++++++++++++++++++++++++++++++++++++++++++| 100% elapsed=10s
> head(cluster.markers, n = 100)
                 p_val avg_log2FC pct.1 pct.2     p_val_adj
Fcmr      0.000000e+00  2.6580313 0.693 0.017  0.000000e+00
Bank1     0.000000e+00  2.5322032 0.711 0.039  0.000000e+00
Pax5      0.000000e+00  2.4695290 0.657 0.017  0.000000e+00
Igkc      0.000000e+00  5.0722286 0.958 0.046  0.000000e+00
Cd79a     0.000000e+00  3.9834118 0.970 0.027  0.000000e+00
Ebf1      0.000000e+00  3.7401276 0.940 0.062  0.000000e+00
Cd79b     0.000000e+00  3.1219841 0.853 0.039  0.000000e+00
Ighd      0.000000e+00  3.4469762 0.886 0.029  0.000000e+00
Ighm      0.000000e+00  3.5589081 0.978 0.246  0.000000e+00
Mef2c     0.000000e+00  3.0843168 0.882 0.140  0.000000e+00
Ly6d      0.000000e+00  3.4892170 0.879 0.031  0.000000e+00
Iglc3     0.000000e+00  2.7172031 0.697 0.021  0.000000e+00
Iglc2     0.000000e+00  3.3441669 0.820 0.023  0.000000e+00
H2-Aa     0.000000e+00  2.3943922 0.996 0.257  0.000000e+00
Ms4a1     0.000000e+00  3.3863465 0.866 0.021  0.000000e+00
Cd74     6.130601e-298  2.3882968 1.000 0.325 1.032148e-293
H2-Eb1   4.302153e-291  2.0269734 0.982 0.230 7.243104e-287
H2-Ab1   1.315096e-290  2.0149622 0.993 0.244 2.214096e-286
Scd1     7.889363e-266  2.4952785 0.756 0.144 1.328253e-261
Cr2      9.414711e-262  2.2997673 0.558 0.015 1.585061e-257
H2-Ob    1.140625e-254  2.2469861 0.618 0.050 1.920356e-250
Gm31243  2.788041e-248  1.9843575 0.531 0.014 4.693945e-244
Bcl11a   2.267203e-246  2.1931573 0.585 0.034 3.817062e-242
Fcer2a   4.309203e-245  2.0960854 0.531 0.016 7.254974e-241
Ralgps2  5.365689e-238  2.3393789 0.684 0.116 9.033674e-234
H2-DMb2  1.856127e-227  1.9522744 0.584 0.050 3.124975e-223
Cd37     5.544814e-214  2.0348142 0.699 0.150 9.335250e-210
Pou2af1  1.033252e-201  1.7633509 0.476 0.023 1.739584e-197
Cd83     1.071778e-198  2.0675395 0.566 0.067 1.804445e-194
Mzb1     1.578859e-189  1.6812998 0.430 0.014 2.658168e-185
Cd22     8.979813e-189  1.7122754 0.455 0.024 1.511841e-184
Tnfrsf13c 9.003371e-189  1.5975205 0.427 0.013 1.515808e-184
Cd19     1.436013e-187  1.6457795 0.419 0.011 2.417672e-183
Pou2f2   3.560917e-180  1.9091689 0.601 0.112 5.995159e-176
H2-Oa    1.622251e-175  1.6599540 0.484 0.047 2.731221e-171
Stap1    6.085959e-174  1.9480454 0.591 0.115 1.024632e-169
Vpreb3   1.355048e-173  1.6746976 0.391 0.010 2.281358e-169
Jund     9.332222e-170  1.5934180 0.955 0.689 1.571173e-165
Fcrla    1.690489e-168  1.5611184 0.396 0.016 2.846107e-164
Hvcn1    1.546337e-164  1.8329939 0.528 0.085 2.603413e-160
```

| | | | | | |
|---|---|---|---|---|---|
| Cxcr4 | 2.963927e-163 | 1.8861429 | 0.759 | 0.274 | 4.990067e-159 |
| Ly6e | 8.019408e-162 | 1.3900666 | 0.938 | 0.614 | 1.350147e-157 |
| Stk17b | 3.492480e-158 | 1.4702308 | 0.925 | 0.546 | 5.879940e-154 |
| Spib | 3.980694e-158 | 1.4809034 | 0.373 | 0.014 | 6.701897e-154 |
| Cd55 | 2.577189e-153 | 1.8810896 | 0.547 | 0.110 | 4.338956e-149 |
| Siglecg | 3.620703e-152 | 1.4736013 | 0.374 | 0.019 | 6.095815e-148 |
| Npc2 | 4.662864e-150 | -1.9361674 | 0.223 | 0.761 | 7.850398e-146 |
| Napsa | 1.126790e-146 | 1.4915262 | 0.521 | 0.087 | 1.897063e-142 |
| Fxyd5 | 6.010977e-145 | -1.8131248 | 0.188 | 0.753 | 1.012008e-140 |
| Blnk | 1.035147e-144 | 1.4885018 | 0.408 | 0.038 | 1.742774e-140 |
| Cd180 | 7.173099e-137 | 1.5735909 | 0.391 | 0.038 | 1.207663e-132 |
| Rasgrp3 | 1.832035e-136 | 1.4037156 | 0.340 | 0.017 | 3.084414e-132 |
| Ciita | 5.631452e-133 | 1.5629577 | 0.438 | 0.065 | 9.481113e-129 |
| Gm30211 | 2.844641e-131 | 1.6423697 | 0.306 | 0.009 | 4.789237e-127 |
| Cxcr5 | 1.675567e-128 | 1.3363281 | 0.320 | 0.016 | 2.820984e-124 |
| H3f3a | 5.568426e-126 | 0.9650033 | 0.978 | 0.906 | 9.375002e-122 |
| Cd69 | 8.601540e-126 | 1.6966716 | 0.668 | 0.226 | 1.448155e-121 |
| Man1a | 1.334656e-119 | 1.6749552 | 0.627 | 0.255 | 2.247027e-115 |
| Fchsd2 | 7.969338e-119 | 1.8157353 | 0.537 | 0.163 | 1.341718e-114 |
| B3gnt5 | 8.496375e-118 | 1.3038391 | 0.325 | 0.025 | 1.430450e-113 |
| BE692007 | 1.165388e-117 | 1.6005070 | 0.554 | 0.156 | 1.962047e-113 |
| H2-Eb2 | 2.136695e-114 | 1.1488385 | 0.280 | 0.011 | 3.597340e-110 |
| Fcrl1 | 2.766658e-114 | 1.1756483 | 0.291 | 0.015 | 4.657946e-110 |
| Arl4c | 6.665471e-113 | -1.4559402 | 0.021 | 0.502 | 1.122199e-108 |
| Iglc1 | 8.274487e-109 | 1.9185369 | 0.268 | 0.011 | 1.393093e-104 |
| Txn1 | 2.229944e-106 | -1.8487135 | 0.270 | 0.705 | 3.754334e-102 |
| Ifitm2 | 3.452274e-106 | -2.3695058 | 0.041 | 0.501 | 5.812249e-102 |
| Lgals1 | 5.974376e-106 | -3.2276843 | 0.177 | 0.616 | 1.005846e-101 |
| Serinc3 | 7.153651e-106 | 1.3038333 | 0.769 | 0.461 | 1.204389e-101 |
| Cd63 | 1.372212e-105 | -2.5107683 | 0.037 | 0.488 | 2.310257e-101 |
| Ctsd | 6.105241e-104 | -1.7989219 | 0.089 | 0.562 | 1.027878e-99 |
| Swap70 | 1.216466e-103 | 1.5437787 | 0.520 | 0.174 | 2.048042e-99 |
| Klf2 | 7.504715e-102 | 1.2241695 | 0.891 | 0.572 | 1.263494e-97 |
| Chst3 | 6.417026e-101 | 1.1645504 | 0.257 | 0.013 | 1.080370e-96 |
| Ptprcap | 3.378129e-98 | 1.3043564 | 0.601 | 0.227 | 5.687418e-94 |
| Ikzf3 | 5.240927e-98 | 1.3473379 | 0.375 | 0.068 | 8.823624e-94 |
| Tsc22d3 | 1.253568e-97 | 1.3772735 | 0.764 | 0.473 | 2.110508e-93 |
| Anxa2 | 3.036627e-97 | -2.1948150 | 0.076 | 0.507 | 5.112465e-93 |
| Zfp36l1 | 5.288706e-97 | 1.3725961 | 0.837 | 0.611 | 8.904066e-93 |
| Cd9 | 5.781223e-97 | -2.4940169 | 0.045 | 0.475 | 9.733267e-93 |
| Serpinb1a | 2.648165e-96 | 1.3221427 | 0.325 | 0.044 | 4.458451e-92 |
| Snn | 3.344955e-95 | 1.3950567 | 0.383 | 0.081 | 5.631566e-91 |
| Chchd10 | 1.055111e-94 | 1.3614879 | 0.399 | 0.085 | 1.776385e-90 |
| Gm8369 | 1.811398e-94 | 1.4846161 | 0.605 | 0.227 | 3.049670e-90 |
| Xist | 2.107184e-94 | 1.2605884 | 0.820 | 0.467 | 3.547654e-90 |
| Pkm | 1.287406e-93 | -1.5086559 | 0.293 | 0.714 | 2.167477e-89 |
| Lgals3 | 1.205158e-92 | -1.9768371 | 0.070 | 0.497 | 2.029003e-88 |
| Syk | 6.916932e-92 | 1.4220055 | 0.472 | 0.142 | 1.164535e-87 |
| Irf8 | 1.463969e-91 | 1.2361998 | 0.537 | 0.188 | 2.464738e-87 |
| Emb | 3.130513e-91 | -1.9380176 | 0.030 | 0.444 | 5.270531e-87 |
| Zfp318 | 6.540463e-91 | 1.4819586 | 0.408 | 0.106 | 1.101152e-86 |
| Serp1 | 5.940389e-90 | 1.2383622 | 0.755 | 0.502 | 1.000124e-85 |

```
Actn1      1.897084e-88 -1.3279241 0.024 0.430  3.193930e-84
Mt1        5.212335e-88 -2.3017351 0.054 0.466  8.775486e-84
Ldha       5.474895e-88 -1.4253451 0.243 0.668  9.217534e-84
Myl6       1.031787e-87 -1.1378593 0.643 0.895  1.737116e-83
Prdx5      1.770777e-87 -1.7539513 0.235 0.625  2.981279e-83
Ddx5       1.785117e-87  0.8057150 0.974 0.839  3.005423e-83
Tagln2     1.934577e-85 -1.3114655 0.126 0.559  3.257053e-81
Ltb        4.141011e-85  1.1123599 0.713 0.329  6.971805e-81
```

**Cluster 1 CD4 T-cells**

```
cluster.markers <- FindMarkers(danx, ident.1 =1, min.pct = 0.25)
  |++++++++++++++++++++++++++++++++++++++++++++++++++| 100% elapsed=09s
> head(cluster.markers, n = 100)
                 p_val avg_log2FC pct.1 pct.2     p_val_adj
Trbc2     0.000000e+00  2.6444429 0.924 0.104  0.000000e+00
Lef1     1.294532e-299  2.5474070 0.850 0.094 2.179474e-295
Cd3d     4.480145e-283  2.2445770 0.894 0.105 7.542772e-279
Tcf7     1.582363e-270  2.4245808 0.873 0.125 2.664066e-266
Ms4a4b   6.322609e-266  2.2625798 0.901 0.122 1.064474e-261
Cd28     2.413297e-255  2.3656143 0.779 0.087 4.063026e-251
Bcl11b   6.188099e-246  2.1691403 0.756 0.084 1.041828e-241
Cd3g     1.303628e-208  1.7975086 0.703 0.086 2.194788e-204
Trac     1.011722e-186  1.7286718 0.645 0.085 1.703336e-182
Itk      3.063671e-178  1.6429087 0.608 0.075 5.157996e-174
Il7r     3.733255e-175  1.9592325 0.668 0.099 6.285309e-171
Txk      1.418447e-173  1.6613236 0.631 0.086 2.388097e-169
Cd3e     3.252179e-165  1.5132715 0.583 0.073 5.475369e-161
Lat      1.593794e-162  1.5386921 0.569 0.072 2.683311e-158
Prkcq    1.199339e-160  1.4915141 0.509 0.054 2.019206e-156
Satb1    5.395607e-154  1.8325920 0.912 0.305 9.084044e-150
Skap1    6.935818e-145  1.4681923 0.548 0.078 1.167714e-140
Emb      1.648035e-144  1.9415222 0.811 0.241 2.774631e-140
Ms4a6b   1.117958e-142  1.6116970 0.786 0.211 1.882195e-138
Trbc1    2.893739e-142  2.1223333 0.541 0.075 4.871899e-138
Rps16    9.468491e-140  0.9753005 0.993 0.982 1.594115e-135
S1pr1    3.354734e-139  1.5591401 0.763 0.188 5.648030e-135
Themis   4.435013e-132  1.2661273 0.454 0.053 7.466787e-128
Cd4      1.219231e-129  1.2462036 0.376 0.029 2.052697e-125
Rpl12    1.074183e-124  1.0285085 0.988 0.949 1.808495e-120
Cd247    6.136966e-119  1.1976564 0.422 0.051 1.033220e-114
Rps24    1.932219e-118  0.9249482 0.998 0.985 3.253084e-114
Rpl5     3.383351e-116  0.9644925 0.993 0.923 5.696210e-112
Nsg2     1.092044e-110  1.1038247 0.380 0.042 1.838565e-106
Fth1     2.018245e-110 -2.1760754 0.963 0.987 3.397917e-106
Thy1     9.447011e-109  1.3331189 0.512 0.096 1.590499e-104
Rps15a   1.973059e-108  0.8503165 0.993 0.962 3.321841e-104
Dapl1    8.053439e-108  1.5660589 0.392 0.046 1.355877e-103
Rpl18    6.978890e-107  0.8486208 0.995 0.946 1.174966e-102
Fam78a   9.811026e-106  1.1374142 0.426 0.063 1.651784e-101
Rplp1    7.807529e-105  0.7912974 1.000 0.987 1.314476e-100
Rps23    3.740046e-104  0.7982672 0.991 0.978 6.296741e-100
```

| | | | | | |
|---|---|---|---|---|---|
| Tpt1 | 1.300509e-103 | 0.6820538 | 0.995 | 0.998 | 2.189538e-99 |
| Trat1 | 1.053667e-102 | 0.9785748 | 0.276 | 0.017 | 1.773955e-98 |
| Cd5 | 9.982318e-102 | 1.1090554 | 0.355 | 0.040 | 1.680623e-97 |
| Rps14 | 3.581568e-100 | 0.7374091 | 0.998 | 0.980 | 6.029928e-96 |
| Vps37b | 5.904253e-100 | 1.5479053 | 0.696 | 0.247 | 9.940401e-96 |
| Fyb | 9.430668e-99 | 1.3434626 | 0.675 | 0.213 | 1.587747e-94 |
| Rps13 | 4.226763e-98 | 0.7767361 | 0.995 | 0.972 | 7.116178e-94 |
| Rpl13a | 5.112227e-98 | 0.7715551 | 0.995 | 0.966 | 8.606945e-94 |
| Lck | 6.522176e-97 | 1.1934959 | 0.454 | 0.082 | 1.098074e-92 |
| Rpl30 | 1.249740e-96 | 0.7294297 | 0.998 | 0.980 | 2.104062e-92 |
| Saraf | 5.730406e-94 | 1.5087012 | 0.661 | 0.260 | 9.647711e-90 |
| Inpp4b | 6.017432e-94 | 1.2992602 | 0.541 | 0.138 | 1.013095e-89 |
| Ets1 | 2.040019e-92 | 1.3199719 | 0.908 | 0.493 | 3.434576e-88 |
| Rps29 | 6.553019e-92 | 0.7316403 | 0.998 | 0.981 | 1.103266e-87 |
| Rps6 | 1.461948e-91 | 0.8237027 | 0.988 | 0.942 | 2.461336e-87 |
| Limd2 | 3.405256e-91 | 1.2930584 | 0.816 | 0.373 | 5.733089e-87 |
| H2-Q7 | 1.795699e-90 | 1.1953381 | 0.795 | 0.312 | 3.023239e-86 |
| Gimap3 | 2.330726e-90 | 1.2902570 | 0.599 | 0.164 | 3.924010e-86 |
| Gm2682 | 3.137373e-89 | 0.9711247 | 0.339 | 0.043 | 5.282082e-85 |
| Grap2 | 5.530135e-89 | 1.1552659 | 0.493 | 0.109 | 9.310535e-85 |
| Rps15 | 9.574242e-87 | 0.7225800 | 0.995 | 0.957 | 1.611919e-82 |
| Hcst | 1.514282e-86 | 1.2308977 | 0.661 | 0.218 | 2.549446e-82 |
| Rps7 | 1.205205e-85 | 0.7546204 | 0.995 | 0.959 | 2.029084e-81 |
| Rps3 | 6.643433e-84 | 0.7436663 | 0.991 | 0.952 | 1.118488e-79 |
| Dgka | 2.899102e-83 | 1.2407311 | 0.514 | 0.138 | 4.880928e-79 |
| Rps3a1 | 8.493875e-83 | 0.6361312 | 0.995 | 0.979 | 1.430029e-78 |
| Gimap6 | 9.805481e-83 | 1.1838152 | 0.767 | 0.286 | 1.650851e-78 |
| Shisa5 | 1.435637e-81 | 1.1044011 | 0.935 | 0.601 | 2.417038e-77 |
| Atp1b3 | 3.139142e-81 | 1.3419885 | 0.705 | 0.335 | 5.285059e-77 |
| H2-K1 | 3.834476e-81 | 0.9812863 | 0.977 | 0.725 | 6.455724e-77 |
| Ablim1 | 3.343371e-80 | 1.1121990 | 0.795 | 0.320 | 5.628899e-76 |
| Rplp2 | 1.945784e-79 | 0.7158542 | 0.993 | 0.951 | 3.275923e-75 |
| Tmsb10 | 5.006154e-79 | 0.8764945 | 0.982 | 0.785 | 8.428360e-75 |
| Rpl19 | 3.238566e-78 | 0.6228639 | 0.995 | 0.975 | 5.452449e-74 |
| Cd27 | 2.061002e-75 | 1.0170413 | 0.364 | 0.066 | 3.469903e-71 |
| Rps19 | 6.936412e-75 | 0.7081722 | 0.995 | 0.955 | 1.167814e-70 |
| Peli1 | 4.251360e-74 | 1.2672217 | 0.680 | 0.291 | 7.157590e-70 |
| Rpl8 | 5.758895e-74 | 0.6660058 | 0.995 | 0.963 | 9.695676e-70 |
| Rpl35a | 2.747114e-73 | 0.6200609 | 1.000 | 0.978 | 4.625040e-69 |
| Spn | 4.427479e-72 | 1.0142474 | 0.366 | 0.073 | 7.454103e-68 |
| Smc4 | 6.145170e-71 | 1.4594632 | 0.650 | 0.281 | 1.034601e-66 |
| Cd6 | 9.876243e-71 | 0.9736029 | 0.302 | 0.045 | 1.662764e-66 |
| Rps28 | 1.014432e-70 | 0.6865755 | 0.995 | 0.948 | 1.707898e-66 |
| Rps10 | 3.408906e-70 | 0.5906980 | 0.993 | 0.985 | 5.739235e-66 |
| Rps21 | 4.723963e-69 | 0.6646083 | 0.993 | 0.963 | 7.953264e-65 |
| Cd74 | 3.012076e-68 | -4.0970834 | 0.152 | 0.577 | 5.071131e-64 |
| Rps5 | 5.312140e-67 | 0.6635770 | 0.998 | 0.946 | 8.943518e-63 |
| H2-Aa | 4.739429e-66 | -3.4912686 | 0.090 | 0.529 | 7.979303e-62 |
| Rps8 | 3.469599e-65 | 0.6010080 | 0.998 | 0.970 | 5.841417e-61 |
| Ltb | 4.308477e-65 | 0.9302147 | 0.820 | 0.363 | 7.253752e-61 |
| H2-Ab1 | 1.085355e-64 | -3.2972965 | 0.083 | 0.519 | 1.827303e-60 |
| Rps20 | 5.882025e-64 | 0.6172692 | 1.000 | 0.968 | 9.902978e-60 |

```
Rapgef6    1.139033e-63  1.1229813 0.700 0.335  1.917677e-59
Arhgap15   1.497680e-63  1.0437633 0.650 0.260  2.521495e-59
Rpl17      3.420248e-63  0.6291010 0.991 0.963  5.758330e-59
Ccr7       4.125385e-63  1.0333539 0.671 0.251  6.945498e-59
Ftl1       9.966953e-63 -1.8713496 0.719 0.878  1.678036e-58
Rps4x      2.324780e-62  0.5643944 0.998 0.947  3.914000e-58
Gimap4     3.997052e-62  0.9942609 0.576 0.192  6.729437e-58
Rps9       8.281657e-62  0.5006397 0.998 0.991  1.394300e-57
H2-Eb1     9.178572e-62 -3.0830867 0.081 0.503  1.545304e-57
Ptpn18     1.865629e-61  0.9402664 0.850 0.460  3.140973e-57
Rpsa       4.775146e-61  0.6174689 1.000 0.955  8.039436e-57
```

**Cluster2 cancer cells**

```
cluster.markers <- FindMarkers(danx, ident.1 =2, min.pct = 0.25)
  |++++++++++++++++++++++++++++++++++++++++++++++++++| 100% elapsed=12s
> head(cluster.markers, n = 100)
                  p_val avg_log2FC pct.1 pct.2     p_val_adj
Gng11      6.592944e-203  2.6002052 0.835 0.148 1.109988e-198
Tnnt2      5.918788e-193  2.2475612 0.744 0.111 9.964872e-189
Krt8       2.802489e-171  2.1151632 0.733 0.120 4.718271e-167
Lgals1     9.280875e-171  2.7854764 0.972 0.423 1.562528e-166
S100a6     6.825891e-167  2.8033668 0.997 0.536 1.149207e-162
Krt18      6.132317e-164  1.8497726 0.713 0.114 1.032437e-159
S100a4     1.979708e-155  2.8324898 0.915 0.286 3.333037e-151
Hmga2      4.074082e-150  1.9841745 0.727 0.134 6.859124e-146
Vim        2.390897e-138  2.0296142 0.967 0.499 4.025315e-134
Cd52       3.938808e-137 -3.6979456 0.055 0.824 6.631377e-133
Tpm1       5.746629e-136  1.9743326 0.807 0.222 9.675025e-132
Ccnd1      9.707790e-134  2.1473020 0.711 0.163 1.634403e-129
Cald1      5.232297e-129  1.7326209 0.741 0.156 8.809096e-125
Prkg2      6.584470e-117  1.5801110 0.579 0.106 1.108561e-112
S100a10    5.284820e-115  2.0296250 0.917 0.552 8.897522e-111
Coro1a     3.355929e-114 -2.7189024 0.036 0.745 5.650042e-110
Lmna       1.453663e-112  1.7051134 0.736 0.195 2.447386e-108
Malat1     5.178687e-111 -2.1299677 0.311 0.897 8.718837e-107
Anxa2      1.904200e-110  1.7521788 0.865 0.317 3.205911e-106
Ptprc      1.675159e-109 -3.0532887 0.017 0.712 2.820298e-105
Tm4sf1     2.427089e-108  1.3830293 0.661 0.137 4.086247e-104
Stk17b     1.042632e-105 -2.7160725 0.080 0.736 1.755376e-101
Serpinb6a  1.215978e-104  1.5285634 0.727 0.209 2.047220e-100
Arhgdib    3.999086e-104 -2.4914795 0.091 0.738 6.732861e-100
S100a11    3.985548e-102  1.7218962 0.923 0.530  6.710069e-98
Srgn        1.365039e-98 -3.1564828 0.036 0.688  2.298179e-94
Fos         1.958852e-95 -3.4453654 0.077 0.691  3.297923e-91
Rhoc        7.318176e-95  1.5359665 0.614 0.164  1.232088e-90
H2-K1       3.166499e-94 -1.9665008 0.287 0.837  5.331117e-90
Laptm5      1.896983e-92 -2.4273331 0.028 0.658  3.193760e-88
Cd53        2.411535e-88 -2.2857270 0.028 0.635  4.060061e-84
Nedd4       6.521114e-88  1.4424583 0.653 0.175  1.097895e-83
Mif         1.547816e-86  1.5419668 0.906 0.594  2.605903e-82
Gpx4        3.368322e-86  1.4885760 0.851 0.493  5.670906e-82
Lcp1        3.299991e-85 -2.4454106 0.039 0.628  5.555865e-81
```

| | | | | | |
|---|---|---|---|---|---|
| Cyba | 4.570164e-84 | -2.3896578 | 0.033 | 0.625 | 7.694329e-80 |
| Hspb1 | 6.244446e-84 | 1.4589468 | 0.512 | 0.114 | 1.051315e-79 |
| Pgk1 | 6.698304e-84 | 1.6772995 | 0.848 | 0.456 | 1.127727e-79 |
| Timp1 | 1.114383e-82 | 1.4900905 | 0.551 | 0.132 | 1.876175e-78 |
| Shisa5 | 3.904188e-81 | -2.0348769 | 0.179 | 0.725 | 6.573091e-77 |
| Rac2 | 4.759131e-80 | -2.1494929 | 0.022 | 0.601 | 8.012473e-76 |
| H2-D1 | 5.425608e-78 | -1.4345501 | 0.364 | 0.844 | 9.134554e-74 |
| Zfp36 | 1.233977e-77 | -2.7768068 | 0.074 | 0.623 | 2.077524e-73 |
| Junb | 2.914876e-77 | -2.1815490 | 0.499 | 0.837 | 4.907485e-73 |
| Xist | 9.879515e-77 | -2.5099227 | 0.069 | 0.639 | 1.663315e-72 |
| Ppp1r14b | 4.301885e-76 | 1.4718171 | 0.625 | 0.234 | 7.242653e-72 |
| Ptpn18 | 1.622254e-75 | -2.0772402 | 0.039 | 0.594 | 2.731227e-71 |
| Btg1 | 1.756783e-75 | -1.6245702 | 0.598 | 0.899 | 2.957719e-71 |
| Anxa1 | 2.483996e-75 | 1.3112842 | 0.719 | 0.247 | 4.182056e-71 |
| Ly6e | 5.389892e-74 | -1.8070051 | 0.275 | 0.768 | 9.074423e-70 |
| Igfbp4 | 6.431222e-74 | 1.7749083 | 0.656 | 0.241 | 1.082761e-69 |
| Nupr1 | 1.134515e-73 | 1.4267376 | 0.532 | 0.149 | 1.910070e-69 |
| Ddx5 | 1.704739e-73 | -1.2930995 | 0.628 | 0.914 | 2.870099e-69 |
| Jund | 1.940590e-73 | -1.9640263 | 0.416 | 0.814 | 3.267177e-69 |
| Txn1 | 1.974202e-73 | 1.2536191 | 0.873 | 0.542 | 3.323766e-69 |
| Id3 | 3.490273e-73 | 1.7174599 | 0.576 | 0.182 | 5.876224e-69 |
| Klf2 | 9.465532e-73 | -2.5020085 | 0.264 | 0.720 | 1.593617e-68 |
| Phlda3 | 1.728772e-72 | 1.2920812 | 0.435 | 0.099 | 2.910560e-68 |
| Sox9 | 4.633834e-72 | 1.2026281 | 0.493 | 0.113 | 7.801523e-68 |
| Tnfrsf12a | 1.884684e-71 | 1.2920767 | 0.501 | 0.126 | 3.173054e-67 |
| Cd74 | 3.974678e-71 | -4.4424280 | 0.058 | 0.579 | 6.691767e-67 |
| 2200002D01Rik | 4.474225e-71 | 1.1967305 | 0.419 | 0.095 | 7.532805e-67 |
| Nme2 | 9.872404e-71 | 1.2614808 | 0.906 | 0.742 | 1.662118e-66 |
| Ctla2a | 2.404649e-70 | 1.2212208 | 0.543 | 0.151 | 4.048466e-66 |
| Tpi1 | 5.422973e-70 | 1.2523945 | 0.771 | 0.340 | 9.130118e-66 |
| Btg2 | 1.625863e-69 | -2.3571327 | 0.052 | 0.575 | 2.737302e-65 |
| Aprt | 3.478944e-69 | 1.3349799 | 0.777 | 0.420 | 5.857150e-65 |
| Pls3 | 1.090891e-66 | 1.2554964 | 0.496 | 0.129 | 1.836624e-62 |
| Rps27l | 1.680665e-66 | 1.4538048 | 0.793 | 0.467 | 2.829567e-62 |
| Ank | 2.602174e-66 | 1.1055894 | 0.534 | 0.146 | 4.381020e-62 |
| Nenf | 2.502795e-65 | 1.3016035 | 0.570 | 0.197 | 4.213705e-61 |
| Wwtr1 | 4.970746e-65 | 1.1488142 | 0.444 | 0.112 | 8.368749e-61 |
| Anxa5 | 6.402793e-65 | 1.4020763 | 0.744 | 0.342 | 1.077974e-60 |
| Cdkn2a | 1.424465e-64 | 1.1872539 | 0.388 | 0.084 | 2.398230e-60 |
| Lmo1 | 2.977062e-63 | 1.1500512 | 0.375 | 0.084 | 5.012181e-59 |
| Akap13 | 4.296955e-63 | -1.5959800 | 0.085 | 0.585 | 7.234354e-59 |
| H2-Ab1 | 5.087440e-63 | -3.5962001 | 0.025 | 0.515 | 8.565214e-59 |
| Gmfg | 1.447633e-62 | -1.8632973 | 0.017 | 0.503 | 2.437235e-58 |
| H2-Aa | 4.874561e-62 | -3.8039243 | 0.044 | 0.523 | 8.206811e-58 |
| Crip1 | 8.403135e-62 | -2.4279648 | 0.041 | 0.528 | 1.414752e-57 |
| Prdx2 | 8.834898e-61 | 1.2654180 | 0.733 | 0.383 | 1.487443e-56 |
| Dap | 1.869529e-60 | 1.3458625 | 0.603 | 0.248 | 3.147539e-56 |
| Ltb | 2.565035e-60 | -2.0394915 | 0.019 | 0.498 | 4.318493e-56 |
| Ighm | 2.617745e-60 | -3.1437180 | 0.033 | 0.510 | 4.407236e-56 |
| Lsp1 | 1.028238e-59 | -1.8161260 | 0.030 | 0.504 | 1.731142e-55 |
| Dstn | 2.198849e-59 | 1.2955754 | 0.642 | 0.277 | 3.701981e-55 |
| Zfp36l1 | 4.126283e-59 | -1.7428711 | 0.314 | 0.728 | 6.947010e-55 |

```
Ywhae           6.741316e-59  1.1125687 0.843 0.518  1.134968e-54
Tceal9          1.691500e-58  1.2841388 0.598 0.237  2.847810e-54
Uqcc2           5.415502e-58  1.3379481 0.661 0.323  9.117538e-54
B2m             6.848314e-58 -0.9944819 0.755 0.947  1.152982e-53
Clk1            3.549692e-57 -1.5106732 0.107 0.570  5.976262e-53
Cytip           1.269238e-56 -1.7966850 0.011 0.467  2.136889e-52
Eef1b2          2.143374e-56  0.9649961 0.948 0.896  3.608584e-52
H2-Eb1          2.563651e-56 -3.1471363 0.044 0.496  4.316162e-52
Serinc3         2.596442e-56 -1.5599612 0.149 0.606  4.371371e-52
Ptma            6.560175e-56  0.9556268 0.964 0.943  1.104471e-51
Satb1           9.753135e-56 -2.3243822 0.008 0.459  1.642038e-51
Mbnl1           1.145433e-55 -1.4410230 0.421 0.773  1.928452e-51
Cxcr4           1.175180e-55 -2.0780470 0.014 0.466  1.978533e-51
```

**Cluster 3 macrophage - 1**

```
cluster.markers <- FindMarkers(danx, ident.1 =3, min.pct = 0.25)
  |++++++++++++++++++++++++++++++++++++++++++++++++++| 100% elapsed=09s
> head(cluster.markers, n = 100)
                        p_val  avg_log2FC pct.1 pct.2      p_val_adj
C1qc           3.305416e-202  2.5182313 0.843 0.105 5.564998e-198
C1qa           5.585068e-202  2.7337662 0.881 0.124 9.403021e-198
C1qb           5.781051e-198  2.6299947 0.854 0.116 9.732977e-194
Aif1           7.346758e-179  2.7875494 0.805 0.128 1.236900e-174
Apoe           1.318585e-134  3.0934119 0.935 0.325 2.219970e-130
Lyz2           2.900098e-133  2.1960125 0.874 0.209 4.882605e-129
Fcer1g         2.669431e-132  2.0434442 0.920 0.215 4.494253e-128
Arg1           3.660182e-129  2.6277490 0.605 0.088 6.162282e-125
Pf4            9.452318e-115  2.2501201 0.540 0.074 1.591392e-110
Fabp5          4.042700e-108  2.9440291 0.759 0.224 6.806289e-104
Mmp13          1.780977e-107  2.3361352 0.421 0.043 2.998453e-103
Ftl1           2.043763e-103  2.0801974 0.992 0.838  3.440880e-99
Bcl2a1b        4.727785e-102  2.0918115 0.755 0.215  7.959698e-98
Tyrobp         1.515708e-101  1.5775054 0.877 0.231  2.551846e-97
Fth1            1.859270e-96  1.8411279 1.000 0.981  3.130266e-92
Mmp12           1.193978e-94  2.7199064 0.402 0.047  2.010181e-90
Ccl12           2.250848e-94  2.0861031 0.398 0.045  3.789527e-90
Fcgr3           3.407408e-89  1.7736210 0.621 0.134  5.736712e-85
Tmsb4x          1.944275e-88  1.2225575 1.000 0.998  3.273381e-84
Mt1             2.025259e-88  2.2906447 0.812 0.305  3.409725e-84
Cxcl16          8.140776e-87  1.8286465 0.506 0.090  1.370581e-82
Clec4n          1.767718e-80  1.6244833 0.487 0.088  2.976130e-76
Malat1          2.191423e-80 -2.6308365 0.314 0.872  3.689480e-76
Ms4a7           3.921904e-76  1.7195185 0.391 0.058  6.602918e-72
Ccl24           2.961490e-73  2.5461185 0.326 0.039  4.985965e-69
Msr1            2.409538e-70  1.6622123 0.425 0.076  4.056698e-66
Adamdec1        4.447690e-70  1.6843357 0.287 0.030  7.488131e-66
Mafb            1.693178e-68  1.6344840 0.525 0.124  2.850635e-64
Ctsc            1.231472e-65  1.8202099 0.762 0.337  2.073305e-61
Ctss            5.103896e-60  1.8202502 0.774 0.378  8.592919e-56
Lgmn            1.245866e-59  1.9658553 0.644 0.247  2.097540e-55
Tmsb10          1.027932e-57 -1.6641686 0.368 0.863  1.730626e-53
Ms4a6d          1.429820e-57  1.3659219 0.410 0.085  2.407245e-53
```

| | | | | | |
|---|---|---|---|---|---|
| Ccl6 | 2.441966e-56 | 1.3748560 | 0.544 | 0.147 | 4.111294e-52 |
| Mrc1 | 3.504507e-56 | 1.3110584 | 0.364 | 0.066 | 5.900188e-52 |
| Trf | 1.435537e-55 | 1.6560044 | 0.467 | 0.119 | 2.416871e-51 |
| Ctsz | 4.968463e-55 | 1.7603140 | 0.670 | 0.295 | 8.364904e-51 |
| Klf2 | 9.219507e-55 | -1.9477089 | 0.142 | 0.714 | 1.552196e-50 |
| AW112010 | 4.946651e-54 | 1.7042760 | 0.693 | 0.306 | 8.328181e-50 |
| Wfdc17 | 7.003156e-54 | 0.6701023 | 0.571 | 0.169 | 1.179051e-49 |
| Dab2 | 2.823920e-52 | 1.7315223 | 0.567 | 0.205 | 4.754352e-48 |
| Fcgr4 | 3.926342e-52 | 1.4779190 | 0.372 | 0.078 | 6.610389e-48 |
| Ntpcr | 5.563109e-52 | 1.5408374 | 0.421 | 0.111 | 9.366050e-48 |
| Atp6v0c | 3.420577e-50 | 1.3984978 | 0.793 | 0.495 | 5.758884e-46 |
| Cd68 | 1.340962e-49 | 1.3577768 | 0.395 | 0.096 | 2.257644e-45 |
| Junb | 9.912611e-49 | -1.6871838 | 0.341 | 0.840 | 1.668887e-44 |
| Xist | 4.283867e-48 | -2.3423909 | 0.107 | 0.612 | 7.212319e-44 |
| Npl | 3.008948e-47 | 1.4273069 | 0.268 | 0.044 | 5.065865e-43 |
| Foxp1 | 8.628217e-47 | -1.6051581 | 0.126 | 0.640 | 1.452647e-42 |
| Bcl2a1a | 1.003585e-46 | 1.3521900 | 0.356 | 0.081 | 1.689636e-42 |
| Acp5 | 2.115966e-46 | 1.5959210 | 0.517 | 0.181 | 3.562440e-42 |
| Cotl1 | 2.066492e-44 | 1.3359229 | 0.789 | 0.525 | 3.479145e-40 |
| Ctsb | 2.162865e-42 | 1.6114683 | 0.697 | 0.386 | 3.641400e-38 |
| Ddx5 | 2.345796e-42 | -1.1318501 | 0.517 | 0.913 | 3.949382e-38 |
| Ms4a6c | 2.367701e-42 | 1.1101981 | 0.425 | 0.118 | 3.986261e-38 |
| Jund | 3.380157e-42 | -1.5591590 | 0.356 | 0.804 | 5.690832e-38 |
| Fos | 3.062850e-39 | -2.0900247 | 0.203 | 0.653 | 5.156614e-35 |
| Ets1 | 3.663482e-39 | -1.4894699 | 0.130 | 0.603 | 6.167838e-35 |
| Csf1r | 1.339308e-38 | 1.3094839 | 0.352 | 0.091 | 2.254859e-34 |
| Mpeg1 | 4.280153e-38 | 1.2203320 | 0.395 | 0.112 | 7.206065e-34 |
| Sdcbp | 1.376361e-37 | 1.2258867 | 0.736 | 0.490 | 2.317242e-33 |
| Luc7l2 | 1.439196e-37 | -1.2150518 | 0.161 | 0.630 | 2.423031e-33 |
| Prdx5 | 2.760177e-37 | 1.3029413 | 0.770 | 0.491 | 4.647035e-33 |
| Mif | 1.169356e-36 | 1.1506997 | 0.854 | 0.612 | 1.968728e-32 |
| Cd300c2 | 1.421859e-36 | 1.1450143 | 0.372 | 0.105 | 2.393841e-32 |
| Cst3 | 4.763771e-36 | 0.9277905 | 0.751 | 0.473 | 8.020284e-32 |
| Msrb1 | 4.948409e-36 | 1.1619586 | 0.586 | 0.278 | 8.331142e-32 |
| Ifitm1 | 1.595133e-35 | 1.0054829 | 0.391 | 0.111 | 2.685565e-31 |
| Ccnl1 | 1.668688e-35 | -1.3520146 | 0.080 | 0.503 | 2.809404e-31 |
| Tmem176b | 2.025408e-35 | 1.0613694 | 0.352 | 0.096 | 3.409976e-31 |
| Clk1 | 2.197371e-34 | -1.2752187 | 0.123 | 0.549 | 3.699494e-30 |
| Ucp2 | 2.199697e-34 | 1.3064879 | 0.663 | 0.374 | 3.703410e-30 |
| Shisa5 | 3.527945e-34 | -1.2352951 | 0.249 | 0.695 | 5.939648e-30 |
| Gm2a | 4.806841e-34 | 1.4666279 | 0.594 | 0.300 | 8.092797e-30 |
| Eif4a2 | 1.743719e-33 | -1.1114077 | 0.161 | 0.599 | 2.935725e-29 |
| Apoc2 | 4.848185e-33 | 1.1558981 | 0.280 | 0.067 | 8.162405e-29 |
| Fcgr1 | 9.525911e-33 | 1.1053840 | 0.261 | 0.059 | 1.603782e-28 |
| Psap | 1.556838e-32 | 1.4186400 | 0.736 | 0.527 | 2.621092e-28 |
| Spi1 | 6.822350e-32 | 1.0964967 | 0.471 | 0.184 | 1.148611e-27 |
| Btg1 | 7.519564e-32 | -1.1002107 | 0.582 | 0.888 | 1.265994e-27 |
| Rbm39 | 8.898972e-32 | -1.0072729 | 0.299 | 0.739 | 1.498231e-27 |
| Spp1 | 1.088760e-31 | 1.1888557 | 0.628 | 0.296 | 1.833036e-27 |
| Selenop | 1.365170e-31 | 1.1383535 | 0.525 | 0.228 | 2.298401e-27 |
| Atox1 | 1.571587e-31 | 1.3453497 | 0.690 | 0.469 | 2.645923e-27 |
| Neat1 | 7.072311e-31 | -1.8977412 | 0.073 | 0.453 | 1.190694e-26 |

| | | | | | |
|---|---|---|---|---|---|
| Ighm | 9.876673e-31 | -2.4370925 | 0.107 | 0.482 | 1.662837e-26 |
| Stk17b | 1.161986e-30 | -1.3242360 | 0.299 | 0.687 | 1.956320e-26 |
| Ltb | 2.202117e-30 | -1.6297763 | 0.103 | 0.469 | 3.707485e-26 |
| Btg2 | 4.967242e-30 | -1.5831770 | 0.149 | 0.544 | 8.362848e-26 |
| 4930523C07Rik | 1.186609e-29 | -1.2392765 | 0.092 | 0.469 | 1.997775e-25 |
| Gapdh | 1.419426e-29 | 1.0665594 | 0.751 | 0.507 | 2.389745e-25 |
| Gatm | 1.739270e-29 | 1.4279177 | 0.433 | 0.184 | 2.928236e-25 |
| Tsc22d3 | 1.765257e-29 | -1.3082181 | 0.192 | 0.591 | 2.971987e-25 |
| Klf6 | 4.462732e-29 | -1.2921913 | 0.207 | 0.603 | 7.513455e-25 |
| Rps20 | 5.235039e-29 | -0.6814047 | 0.985 | 0.972 | 8.813712e-25 |
| Rps27 | 2.983408e-28 | -0.9728570 | 0.533 | 0.873 | 5.022865e-24 |
| Fosb | 8.573922e-28 | -1.5693825 | 0.092 | 0.445 | 1.443506e-23 |
| Macf1 | 1.927646e-27 | -1.0055821 | 0.161 | 0.550 | 3.245385e-23 |
| Zfp36l1 | 2.128664e-27 | -1.2055458 | 0.314 | 0.711 | 3.583818e-23 |
| Rpsa | 4.420451e-27 | -0.6727000 | 0.916 | 0.967 | 7.442271e-23 |

**Cluster 4 macrophage**

```
cluster.markers <- FindMarkers(danx, ident.1 = 4, min.pct = 0.25)
  |++++++++++++++++++++++++++++++++++++++++++++++++++| 100% elapsed=11s
> head(cluster.markers, n = 100)
```

| | p_val | avg_log2FC | pct.1 | pct.2 | p_val_adj |
|---|---|---|---|---|---|
| Ccr2 | 7.486681e-215 | 2.9145359 | 0.575 | 0.025 | 1.260458e-210 |
| Clec4a3 | 3.995151e-199 | 1.3411051 | 0.509 | 0.018 | 6.726236e-195 |
| Csf1r | 5.822163e-196 | 1.9521025 | 0.721 | 0.061 | 9.802194e-192 |
| Mpeg1 | 4.910990e-190 | 2.1703026 | 0.783 | 0.081 | 8.268143e-186 |
| Ccl9 | 9.225239e-186 | 2.4519573 | 0.619 | 0.044 | 1.553161e-181 |
| Ccr5 | 1.183972e-182 | 1.6064800 | 0.602 | 0.040 | 1.993336e-178 |
| Cfp | 4.432870e-180 | 1.5308875 | 0.655 | 0.052 | 7.463180e-176 |
| Plbd1 | 1.711718e-174 | 1.4797275 | 0.659 | 0.053 | 2.881848e-170 |
| Ms4a6c | 1.191477e-172 | 2.2370145 | 0.765 | 0.092 | 2.005971e-168 |
| C3ar1 | 1.335630e-170 | 1.4401350 | 0.553 | 0.035 | 2.248667e-166 |
| Itgam | 1.741263e-168 | 1.5045052 | 0.664 | 0.057 | 2.931591e-164 |
| Sirpa | 2.050092e-167 | 1.6663228 | 0.770 | 0.090 | 3.451534e-163 |
| Pirb | 4.037123e-167 | 1.2538800 | 0.708 | 0.066 | 6.796901e-163 |
| Csf2rb | 1.590983e-163 | 1.3550568 | 0.664 | 0.061 | 2.678580e-159 |
| Cysltr1 | 3.027248e-160 | 0.8698410 | 0.442 | 0.018 | 5.096675e-156 |
| Pld4 | 7.392933e-158 | 1.4095212 | 0.681 | 0.069 | 1.244674e-153 |
| Csf2ra | 3.005411e-154 | 0.9972525 | 0.588 | 0.047 | 5.059911e-150 |
| Adgre1 | 3.409373e-152 | 1.2237033 | 0.527 | 0.036 | 5.740021e-148 |
| Fcgr1 | 1.331156e-151 | 1.1977137 | 0.535 | 0.038 | 2.241135e-147 |
| Alox5ap | 3.775033e-151 | 1.6002660 | 0.699 | 0.076 | 6.355645e-147 |
| Mrc1 | 1.628502e-148 | 2.1025250 | 0.575 | 0.051 | 2.741746e-144 |
| Cd68 | 2.062121e-146 | 1.3431984 | 0.677 | 0.075 | 3.471788e-142 |
| Csf2rb2 | 6.699180e-146 | 0.7641120 | 0.460 | 0.026 | 1.127874e-141 |
| Ms4a6d | 1.212897e-144 | 1.5957532 | 0.646 | 0.069 | 2.042034e-140 |
| Fcgr2b | 2.196198e-144 | 2.0141627 | 0.726 | 0.102 | 3.697519e-140 |
| Olfm1 | 3.316754e-144 | 1.0799251 | 0.504 | 0.036 | 5.584086e-140 |
| Clec4a1 | 6.277212e-143 | 0.9458184 | 0.314 | 0.006 | 1.056831e-138 |
| Rassf4 | 3.652358e-142 | 0.9525945 | 0.478 | 0.031 | 6.149110e-138 |
| Clec10a | 1.926517e-140 | 1.2106961 | 0.292 | 0.004 | 3.243485e-136 |
| Cd300c2 | 1.746008e-139 | 1.4690636 | 0.681 | 0.081 | 2.939579e-135 |
| Pid1 | 5.268490e-139 | 1.2912431 | 0.451 | 0.029 | 8.870030e-135 |

| | | | | | |
|---|---|---|---|---|---|
| Lair1 | 1.697111e-134 | 1.4961844 | 0.540 | 0.050 | 2.857255e-130 |
| Spint1 | 1.980834e-134 | 0.9942573 | 0.434 | 0.026 | 3.334933e-130 |
| Tbxas1 | 6.277205e-133 | 0.7521968 | 0.385 | 0.018 | 1.056830e-128 |
| Trem2 | 7.846557e-133 | 1.1022519 | 0.513 | 0.042 | 1.321046e-128 |
| Mertk | 1.667623e-132 | 0.6128315 | 0.332 | 0.011 | 2.807610e-128 |
| Cx3cr1 | 5.778486e-132 | 1.5875245 | 0.447 | 0.030 | 9.728660e-128 |
| Clec4a2 | 9.885102e-131 | 1.1311011 | 0.553 | 0.052 | 1.664256e-126 |
| Cd300a | 6.398784e-128 | 0.9204561 | 0.460 | 0.033 | 1.077299e-123 |
| Lrrc25 | 7.792424e-128 | 0.7269736 | 0.465 | 0.033 | 1.311933e-123 |
| Slamf9 | 1.599394e-125 | 0.8186367 | 0.412 | 0.025 | 2.692740e-121 |
| P2ry6 | 1.173634e-124 | 0.7201477 | 0.412 | 0.026 | 1.975930e-120 |
| F13a1 | 5.066055e-123 | 2.0687301 | 0.394 | 0.024 | 8.529211e-119 |
| Lilrb4a | 5.236811e-123 | 1.0314705 | 0.642 | 0.077 | 8.816694e-119 |
| Fcer1g | 2.433441e-122 | 2.1171333 | 0.920 | 0.225 | 4.096941e-118 |
| Tyrobp | 1.689323e-120 | 2.0248637 | 0.934 | 0.235 | 2.844145e-116 |
| Ccl6 | 1.378329e-119 | 2.2667333 | 0.761 | 0.133 | 2.320556e-115 |
| Tmem176a | 1.055863e-118 | 1.5931604 | 0.584 | 0.072 | 1.777651e-114 |
| Tnfsf13 | 1.363595e-118 | 0.7179979 | 0.363 | 0.019 | 2.295749e-114 |
| Apoc2 | 2.062266e-118 | 1.3179024 | 0.509 | 0.049 | 3.472031e-114 |
| Igsf6 | 1.644229e-116 | 0.7239505 | 0.465 | 0.039 | 2.768223e-112 |
| Hfe | 3.803651e-116 | 0.7009115 | 0.358 | 0.020 | 6.403826e-112 |
| Msr1 | 4.678772e-115 | 1.4180168 | 0.571 | 0.068 | 7.877181e-111 |
| Emilin2 | 6.118121e-115 | 1.0818534 | 0.558 | 0.063 | 1.030047e-110 |
| Grn | 8.075205e-114 | 1.8987826 | 0.845 | 0.212 | 1.359542e-109 |
| F10 | 1.130672e-113 | 0.7398188 | 0.270 | 0.007 | 1.903599e-109 |
| Dhrs3 | 1.616170e-113 | 0.8056992 | 0.447 | 0.039 | 2.720984e-109 |
| Trf | 1.929105e-113 | 1.9421374 | 0.664 | 0.106 | 3.247840e-109 |
| Naip2 | 1.935364e-113 | 0.6189266 | 0.345 | 0.018 | 3.258378e-109 |
| Lyz2 | 1.883057e-112 | 3.3088760 | 0.810 | 0.223 | 3.170315e-108 |
| C5ar1 | 7.492212e-112 | 1.0384898 | 0.527 | 0.056 | 1.261389e-107 |
| Mafb | 1.120684e-111 | 2.2439369 | 0.664 | 0.117 | 1.886783e-107 |
| Slc8a1 | 2.520874e-111 | 0.6348464 | 0.341 | 0.018 | 4.244143e-107 |
| Naaa | 3.805889e-111 | 1.4157431 | 0.535 | 0.066 | 6.407595e-107 |
| Tifab | 8.546939e-111 | 0.5648720 | 0.385 | 0.025 | 1.438963e-106 |
| Zeb2 | 2.694251e-110 | 1.8979495 | 0.805 | 0.204 | 4.536042e-106 |
| Atf3 | 6.300058e-110 | 1.5597199 | 0.558 | 0.074 | 1.060678e-105 |
| Gda | 2.125809e-109 | 1.2230504 | 0.704 | 0.112 | 3.579012e-105 |
| Cst3 | 8.317356e-108 | 2.9741301 | 0.956 | 0.459 | 1.400310e-103 |
| Psap | 2.649459e-106 | 2.4814925 | 0.973 | 0.509 | 4.460629e-102 |
| Slc11a1 | 3.830757e-106 | 0.6752896 | 0.385 | 0.028 | 6.449462e-102 |
| Clec4n | 1.287318e-105 | 1.4142280 | 0.597 | 0.084 | 2.167329e-101 |
| Tmem176b | 2.150540e-105 | 1.5504018 | 0.571 | 0.080 | 3.620649e-101 |
| Cd302 | 9.348497e-105 | 0.8988874 | 0.491 | 0.053 | 1.573913e-100 |
| Fes | 2.590244e-104 | 0.6619256 | 0.469 | 0.047 | 4.360935e-100 |
| Evi2a | 5.313906e-104 | 0.9955980 | 0.602 | 0.090 | 8.946492e-100 |
| Hacd4 | 6.284761e-104 | 0.9022710 | 0.473 | 0.051 | 1.058102e-99 |
| Ctss | 8.638267e-104 | 2.2527596 | 0.934 | 0.370 | 1.454339e-99 |
| Cebpa | 9.764370e-104 | 0.6890983 | 0.398 | 0.033 | 1.643929e-99 |
| Shtn1 | 1.164979e-103 | 0.5390791 | 0.363 | 0.024 | 1.961358e-99 |
| Tmem106a | 5.841985e-103 | 0.8174694 | 0.438 | 0.044 | 9.835566e-99 |
| Slc15a3 | 7.214760e-103 | 0.7079078 | 0.465 | 0.046 | 1.214677e-98 |
| Sirpb1c | 1.226463e-102 | 0.7773722 | 0.456 | 0.044 | 2.064874e-98 |

```
Cybb      6.256996e-102  1.7429814 0.832 0.205  1.053428e-97
Ifi30     7.242782e-102  1.9466142 0.867 0.252  1.219395e-97
Ifi207    8.320820e-102  1.0003418 0.500 0.062  1.400893e-97
Tgfbi     1.869721e-101  2.1592460 0.765 0.192  3.147863e-97
Rab3il1    1.756092e-99  0.7132039 0.434 0.042  2.956557e-95
Nfam1      6.723534e-99  0.7170652 0.504 0.057  1.131974e-94
Cd14       1.558101e-98  1.1718052 0.606 0.102  2.623220e-94
Ccr1       1.586948e-97  1.0931140 0.624 0.095  2.671785e-93
Ctsz       2.186947e-97  1.7999442 0.898 0.280  3.681945e-93
Ccrl2      3.221208e-97  0.6805951 0.425 0.041  5.423227e-93
Fcgr4      4.320661e-97  1.3060132 0.522 0.069  7.274265e-93
Wfdc17     8.924400e-97  1.1716743 0.765 0.157  1.502512e-92
P2ry14     2.033049e-96  0.5828204 0.274 0.012  3.422841e-92
Fcgr3      3.558025e-96  1.6357674 0.686 0.135  5.990290e-92
Gpr35      6.533225e-96  0.6093655 0.350 0.026  1.099934e-91
Nxpe5      1.512366e-95  0.4809150 0.257 0.010  2.546219e-91
Lilr4b     5.623050e-95  0.5997676 0.504 0.060  9.466967e-91
```

**Cluster 5 CD8 Tcells**

```
cluster.markers <- FindMarkers(danx, ident.1 = 5, min.pct = 0.25)
  |++++++++++++++++++++++++++++++++++++++++++++++++++| 100% elapsed=07s
> head(cluster.markers, n = 100)
                  p_val avg_log2FC pct.1 pct.2     p_val_adj
Nkg7      1.797409e-218  2.5869612 0.635 0.032 3.026117e-214
Il2rb     1.844106e-190  1.9345172 0.502 0.019 3.104737e-186
Cd8b1     3.758711e-156  1.9407314 0.576 0.044 6.328166e-152
Cd8a      6.133212e-150  1.6152904 0.507 0.033 1.032588e-145
Klrd1     1.440056e-135  1.6949052 0.547 0.048 2.424478e-131
Ctsw      2.715481e-128  0.9896830 0.389 0.019 4.571784e-124
Cd3d      2.085061e-123  1.8589572 0.892 0.177 3.510409e-119
Ccl5      3.143120e-122  4.3761170 0.389 0.022 5.291757e-118
Cst7      3.011525e-108  0.8666569 0.315 0.014 5.070204e-104
Ms4a4b    3.038403e-106  1.9414231 0.847 0.196 5.115455e-102
Trbc2     3.068431e-105  1.7636314 0.872 0.182 5.166011e-101
Cd3g      1.651908e-103  1.6231071 0.749 0.138  2.781152e-99
Cd3e       7.142187e-89  1.3160819 0.640 0.114  1.202459e-84
Cxcr6      3.238233e-88  0.9278595 0.251 0.011  5.451889e-84
Lck        1.335137e-87  1.2840771 0.606 0.103  2.247837e-83
Il7r       3.733415e-87  1.8032607 0.700 0.147  6.285577e-83
Itk        2.794563e-83  1.2086348 0.645 0.120  4.704927e-79
Skap1      2.328440e-82  1.2049130 0.626 0.115  3.920162e-78
Ly6c2      5.470845e-82  1.6385490 0.389 0.042  9.210715e-78
Thy1       3.030406e-80  1.3152608 0.631 0.124  5.101992e-76
Bcl11b     3.562937e-80  1.2128490 0.714 0.148  5.998561e-76
Cd28       2.177711e-78  1.2659155 0.719 0.154  3.666394e-74
Cd27       2.414658e-78  1.1377827 0.517 0.081  4.065319e-74
Trac       4.661573e-77  1.2989683 0.655 0.134  7.848225e-73
Tcf7       2.952054e-75  1.3970989 0.773 0.200  4.970079e-71
H2-Q7      1.836844e-71  1.4358195 0.911 0.346  3.092511e-67
Sidt1      4.139455e-71  0.8462950 0.369 0.043  6.969186e-67
Klk8       1.979847e-70  1.0412938 0.468 0.075  3.333271e-66
```

| | | | | | |
|---|---|---|---|---|---|
| Icos | 2.679938e-70 | 1.3861800 | 0.340 | 0.037 | 4.511944e-66 |
| Lat | 3.002572e-68 | 1.1498776 | 0.576 | 0.116 | 5.055130e-64 |
| Txk | 6.802237e-68 | 1.4882869 | 0.606 | 0.137 | 1.145225e-63 |
| Trbc1 | 5.570475e-67 | 1.8244809 | 0.557 | 0.116 | 9.378452e-63 |
| Ptpn22 | 2.289789e-66 | 1.3484259 | 0.635 | 0.159 | 3.855088e-62 |
| Sh2d1a | 2.529465e-66 | 1.1580468 | 0.365 | 0.046 | 4.258608e-62 |
| Hcst | 4.295772e-63 | 1.3323463 | 0.783 | 0.249 | 7.232361e-59 |
| Cd6 | 1.667133e-61 | 1.0507139 | 0.399 | 0.061 | 2.806785e-57 |
| Gm15472 | 3.515611e-60 | 1.1432128 | 0.453 | 0.083 | 5.918882e-56 |
| Ccnd2 | 4.390004e-60 | 1.2457917 | 0.759 | 0.237 | 7.391011e-56 |
| Gm2682 | 6.528843e-60 | 0.8546540 | 0.409 | 0.064 | 1.099196e-55 |
| Cd226 | 1.566299e-59 | 0.8873650 | 0.350 | 0.047 | 2.637020e-55 |
| Ctla4 | 1.840287e-59 | 1.6381069 | 0.251 | 0.023 | 3.098308e-55 |
| Epsti1 | 9.068090e-59 | 1.4759239 | 0.655 | 0.189 | 1.526704e-54 |
| Selplg | 3.697330e-58 | 1.2159510 | 0.709 | 0.221 | 6.224826e-54 |
| Dapl1 | 5.141642e-56 | 1.4539814 | 0.419 | 0.075 | 8.656469e-52 |
| Emb | 1.691361e-55 | 1.3611199 | 0.808 | 0.292 | 2.847576e-51 |
| AW112010 | 8.046753e-55 | 1.7230254 | 0.803 | 0.306 | 1.354751e-50 |
| Ikzf2 | 4.365437e-53 | 2.0146699 | 0.335 | 0.054 | 7.349650e-49 |
| Fth1 | 1.455623e-52 | -2.2487493 | 0.961 | 0.985 | 2.450686e-48 |
| Zap70 | 3.391000e-52 | 0.7092115 | 0.325 | 0.046 | 5.709087e-48 |
| Themis | 1.834259e-51 | 0.8511026 | 0.453 | 0.089 | 3.088159e-47 |
| Ms4a6b | 4.873447e-51 | 1.0781804 | 0.778 | 0.263 | 8.204935e-47 |
| Prkch | 1.121145e-50 | 0.8136565 | 0.424 | 0.081 | 1.887560e-46 |
| Il18r1 | 1.726591e-50 | 0.7751585 | 0.261 | 0.030 | 2.906889e-46 |
| Cd96 | 1.644282e-49 | 0.7138430 | 0.281 | 0.036 | 2.768314e-45 |
| Lef1 | 2.458165e-48 | 0.9202443 | 0.645 | 0.178 | 4.138567e-44 |
| Sh2d2a | 5.633531e-47 | 0.8654079 | 0.365 | 0.065 | 9.484612e-43 |
| Gimap4 | 5.141462e-46 | 1.1156133 | 0.680 | 0.218 | 8.656165e-42 |
| Tnfaip3 | 1.399066e-45 | 1.1286807 | 0.601 | 0.182 | 2.355467e-41 |
| Gimap3 | 4.901707e-45 | 1.0409444 | 0.645 | 0.200 | 8.252514e-41 |
| Tnfrsf18 | 1.557915e-44 | 0.9384438 | 0.291 | 0.044 | 2.622906e-40 |
| Ipcef1 | 2.105988e-44 | 0.8316567 | 0.429 | 0.093 | 3.545642e-40 |
| Cd2 | 3.650443e-44 | 0.9512766 | 0.547 | 0.146 | 6.145887e-40 |
| Gm43065 | 2.285188e-43 | 0.8365562 | 0.315 | 0.055 | 3.847343e-39 |
| Spn | 8.790210e-43 | 0.8765666 | 0.419 | 0.095 | 1.479920e-38 |
| Fam189b | 1.169464e-42 | 0.8609595 | 0.369 | 0.077 | 1.968910e-38 |
| Plcxd2 | 3.574763e-42 | 0.6287656 | 0.315 | 0.054 | 6.018471e-38 |
| Pdcd4 | 3.757007e-42 | 1.0565237 | 0.739 | 0.301 | 6.325297e-38 |
| Nsg2 | 7.413502e-39 | 0.7176283 | 0.360 | 0.074 | 1.248137e-34 |
| Prkcq | 1.181984e-38 | 0.7805959 | 0.424 | 0.101 | 1.989989e-34 |
| Ptprc | 2.577238e-37 | 0.9445189 | 0.961 | 0.594 | 4.339038e-33 |
| Cd247 | 5.133744e-36 | 0.7851279 | 0.379 | 0.088 | 8.643171e-32 |
| H2-K1 | 6.888119e-36 | 0.9124761 | 0.966 | 0.749 | 1.159684e-31 |
| Mbnl1 | 1.277516e-35 | 0.9663068 | 0.946 | 0.709 | 2.150826e-31 |
| Cd5 | 1.303212e-34 | 0.6981174 | 0.330 | 0.071 | 2.194088e-30 |
| Ablim1 | 1.405609e-34 | 0.8365646 | 0.833 | 0.360 | 2.366484e-30 |
| Atp8b4 | 1.899899e-34 | 0.6057795 | 0.261 | 0.046 | 3.198670e-30 |
| Grap2 | 2.316302e-34 | 0.7646930 | 0.493 | 0.144 | 3.899726e-30 |
| Inpp4b | 1.970151e-33 | 0.9998466 | 0.522 | 0.176 | 3.316946e-29 |
| Rinl | 2.089483e-33 | 0.6775331 | 0.365 | 0.089 | 3.517854e-29 |
| Tmsb10 | 7.795654e-33 | 0.8012428 | 0.970 | 0.804 | 1.312476e-28 |

```
Vps37b     1.151636e-32  0.9220883 0.670 0.290  1.938895e-28
H2-Q6      1.156005e-32  0.7823813 0.478 0.145  1.946251e-28
Stat4      4.157875e-32  0.7794018 0.330 0.076  7.000198e-28
Peli1      4.268284e-32  1.0451075 0.680 0.326  7.186083e-28
Saraf      4.613236e-32  0.9344158 0.650 0.297  7.766845e-28
Fyb        7.622956e-32  0.8714329 0.640 0.257  1.283401e-27
Il27ra     8.395746e-32  0.6200305 0.286 0.058  1.413508e-27
Ubash3a    3.020666e-31  0.6355767 0.251 0.046  5.085593e-27
Ptpn18     7.917389e-31  0.8100052 0.901 0.491  1.332972e-26
Rapgef6    8.427589e-31  0.9722422 0.729 0.366  1.418869e-26
H2-Ab1     8.951845e-31 -3.3533046 0.059 0.481  1.507133e-26
Rpl18      9.853667e-31  0.6353543 0.990 0.951  1.658963e-26
Hsd11b1    1.259433e-30  0.5581681 0.271 0.055  2.120381e-26
Gramd3     2.767235e-30  0.8343842 0.448 0.144  4.658917e-26
Shisa5     3.811612e-30  0.8360219 0.926 0.632  6.417230e-26
Itpkb      1.047163e-29  0.8269502 0.537 0.197  1.763004e-25
Ppm1h      1.996704e-29  0.6440226 0.350 0.094  3.361651e-25
Smc4       2.285514e-29  0.9668481 0.670 0.312  3.847891e-25
Fyn        3.808444e-29  0.8536446 0.468 0.164  6.411896e-25
Rps16      4.077547e-29  0.5904733 0.990 0.984  6.864959e-25
```

**Cluster 6 Cancer cells 2**

```
cluster.markers <- FindMarkers(danx, ident.1 = 6, min.pct = 0.25)
  |++++++++++++++++++++++++++++++++++++++++++++++++++| 100% elapsed=23s
> head(cluster.markers, n = 100)
                   p_val avg_log2FC pct.1 pct.2     p_val_adj
Lamc2      5.130054e-305  1.6788764 0.781 0.023 8.636959e-301
Fbln2      4.996883e-296  2.5153850 0.975 0.052 8.412752e-292
Ncam1      3.561863e-290  1.6088236 0.887 0.041 5.996752e-286
Lama5      7.392361e-288  1.8453435 0.944 0.053 1.244578e-283
Lgr6       5.198722e-287  1.4360754 0.869 0.039 8.752568e-283
Tfap2b     7.085840e-282  1.1190043 0.744 0.023 1.192972e-277
Igfbp3     4.191370e-278  2.3842360 0.869 0.041 7.056591e-274
Neo1       1.762826e-277  0.9993299 0.856 0.040 2.967893e-273
Ptprk      1.356073e-276  1.0699423 0.844 0.039 2.283085e-272
Unc5b      1.069312e-268  1.3260215 0.894 0.048 1.800293e-264
Pcdh7      4.470323e-265  1.7038695 0.887 0.051 7.526237e-261
Adgrl3     5.790160e-262  1.3122846 0.844 0.044 9.748313e-258
Efnb1      1.184621e-257  1.5135869 0.881 0.053 1.994427e-253
Itgb4      2.365199e-257  0.9425392 0.662 0.019 3.982048e-253
Slco2a1    1.415761e-256  2.1991599 0.938 0.065 2.383575e-252
Mir100hg   8.268627e-254  1.8081121 0.919 0.062 1.392106e-249
Ano1       1.329677e-253  1.4971401 0.925 0.057 2.238645e-249
Pxdn       8.524040e-251  1.1673408 0.863 0.048 1.435107e-246
Msln       1.509300e-250  1.0105375 0.656 0.019 2.541057e-246
Tnc        1.228283e-249  1.7032615 0.713 0.027 2.067938e-245
Bdnf       1.100669e-248  1.3706195 0.831 0.045 1.853086e-244
Clu        1.467981e-246  2.4081440 0.938 0.069 2.471492e-242
Gm14636    6.603362e-246  0.9847173 0.731 0.030 1.111742e-241
Ccbe1      1.094086e-244  1.2506062 0.856 0.050 1.842003e-240
Jag1       4.418768e-244  2.2903147 0.938 0.070 7.439438e-240
```

| | | | | | |
|---|---|---|---|---|---|
| Pgf | 3.208494e-243 | 1.2462210 | 0.825 | 0.046 | 5.401821e-239 |
| Itga3 | 2.253798e-242 | 1.3961888 | 0.806 | 0.043 | 3.794494e-238 |
| Tns4 | 2.719452e-240 | 1.5863827 | 0.900 | 0.060 | 4.578470e-236 |
| Vldlr | 2.821225e-240 | 0.7577602 | 0.606 | 0.016 | 4.749815e-236 |
| Cpe | 1.255397e-239 | 2.5035452 | 0.988 | 0.084 | 2.113586e-235 |
| Magi1 | 4.174426e-239 | 0.7569803 | 0.762 | 0.036 | 7.028064e-235 |
| Piezo2 | 1.014856e-238 | 1.9104444 | 0.881 | 0.059 | 1.708612e-234 |
| Smoc2 | 3.632847e-238 | 2.5423409 | 0.969 | 0.082 | 6.116261e-234 |
| Ltbp1 | 7.507251e-238 | 1.7166581 | 0.831 | 0.051 | 1.263921e-233 |
| Ackr3 | 5.064764e-237 | 1.1066641 | 0.856 | 0.052 | 8.527037e-233 |
| Lamb1 | 3.544285e-235 | 2.0049766 | 0.981 | 0.082 | 5.967158e-231 |
| Sdc1 | 1.737663e-233 | 1.9502032 | 0.956 | 0.079 | 2.925530e-229 |
| Adgrg6 | 1.803362e-233 | 1.1414101 | 0.744 | 0.036 | 3.036140e-229 |
| Slc47a1 | 2.205927e-233 | 0.8409313 | 0.706 | 0.030 | 3.713899e-229 |
| Bmp7 | 2.487010e-231 | 1.0522356 | 0.800 | 0.044 | 4.187130e-227 |
| Pkp1 | 5.808508e-230 | 1.2005494 | 0.812 | 0.048 | 9.779204e-226 |
| Bmp2 | 1.700371e-228 | 1.8714224 | 0.875 | 0.059 | 2.862744e-224 |
| Tfap2a | 6.551656e-228 | 0.8637825 | 0.800 | 0.045 | 1.103037e-223 |
| Nav2 | 1.896357e-226 | 1.0537049 | 0.850 | 0.054 | 3.192706e-222 |
| Flt1 | 4.087071e-226 | 0.9744887 | 0.731 | 0.035 | 6.880992e-222 |
| Tmprss11e | 5.761814e-225 | 0.6819822 | 0.444 | 0.004 | 9.700591e-221 |
| Wnt5a | 8.058202e-224 | 1.1667449 | 0.844 | 0.056 | 1.356679e-219 |
| Col18a1 | 1.113508e-223 | 2.1267655 | 0.938 | 0.075 | 1.874703e-219 |
| Gja1 | 6.371827e-223 | 1.0240103 | 0.856 | 0.057 | 1.072761e-218 |
| Epha2 | 3.292306e-222 | 0.8753976 | 0.762 | 0.042 | 5.542926e-218 |
| Has2 | 4.350734e-222 | 1.1123451 | 0.688 | 0.031 | 7.324895e-218 |
| Mab21l4 | 3.144671e-221 | 0.8122115 | 0.600 | 0.019 | 5.294368e-217 |
| Acvr1 | 6.546220e-221 | 0.8916899 | 0.787 | 0.046 | 1.102122e-216 |
| Angptl2 | 1.622611e-220 | 1.4883229 | 0.900 | 0.068 | 2.731828e-216 |
| Abcb1a | 2.426844e-218 | 1.3208608 | 0.838 | 0.057 | 4.085835e-214 |
| Ahnak2 | 6.989387e-217 | 1.1306013 | 0.838 | 0.058 | 1.176733e-212 |
| Tnfrsf23 | 5.197559e-215 | 0.9949921 | 0.887 | 0.066 | 8.750610e-211 |
| Selp | 1.327349e-214 | 0.9903695 | 0.475 | 0.008 | 2.234724e-210 |
| Inhba | 4.305094e-214 | 1.9828699 | 0.919 | 0.077 | 7.248055e-210 |
| Col7a1 | 8.959584e-214 | 1.1208285 | 0.750 | 0.043 | 1.508436e-209 |
| Areg | 2.882283e-213 | 2.7974220 | 0.912 | 0.082 | 4.852611e-209 |
| Rdh10 | 4.384293e-213 | 0.9737978 | 0.806 | 0.053 | 7.381396e-209 |
| Epn2 | 1.477432e-211 | 1.0480943 | 0.850 | 0.062 | 2.487405e-207 |
| Fgfr1 | 1.831356e-211 | 1.9462597 | 0.981 | 0.100 | 3.083272e-207 |
| Pcdh19 | 4.483609e-211 | 0.8117971 | 0.619 | 0.023 | 7.548604e-207 |
| Ptprs | 4.038874e-210 | 1.5150024 | 0.931 | 0.085 | 6.799848e-206 |
| Sema3a | 7.666798e-210 | 0.8203571 | 0.637 | 0.028 | 1.290782e-205 |
| Klra4 | 9.511685e-210 | 1.4794124 | 0.825 | 0.057 | 1.601387e-205 |
| Tbx3 | 1.104466e-209 | 0.7149889 | 0.731 | 0.041 | 1.859479e-205 |
| Cadm1 | 8.624313e-208 | 1.0225103 | 0.775 | 0.050 | 1.451989e-203 |
| Plod2 | 1.636787e-206 | 1.3763156 | 0.900 | 0.072 | 2.755694e-202 |
| Fnbp1l | 3.409132e-206 | 0.9685685 | 0.869 | 0.068 | 5.739614e-202 |
| Ghr | 2.262448e-205 | 1.4833689 | 0.925 | 0.084 | 3.809058e-201 |
| Etv1 | 4.474833e-205 | 1.1607311 | 0.875 | 0.072 | 7.533830e-201 |
| Bmp1 | 5.111999e-205 | 1.4792674 | 0.863 | 0.067 | 8.606562e-201 |
| Prrx1 | 6.013640e-205 | 0.7822096 | 0.781 | 0.047 | 1.012456e-200 |
| Htra1 | 6.800711e-205 | 1.2473065 | 0.856 | 0.067 | 1.144968e-200 |

```
Lamc1      2.679756e-204  1.4054391 0.887 0.074 4.511637e-200
Cdh2       2.844672e-204  0.8257599 0.744 0.044 4.789290e-200
Abca5      3.591369e-204  0.6341575 0.644 0.030 6.046428e-200
Zfp503     9.062269e-204  0.9161075 0.769 0.050 1.525724e-199
B3gnt3     1.363903e-203  0.5799887 0.644 0.030 2.296268e-199
Clstn1     2.040750e-203  0.8037746 0.744 0.046 3.435807e-199
Gprc5a     6.221997e-203  0.7722834 0.662 0.033 1.047535e-198
Cxadr      1.887710e-201  0.6509931 0.650 0.031 3.178149e-197
Map1b      7.274232e-201  1.3546224 0.925 0.084 1.224690e-196
Asap2      2.017407e-200  1.3014721 0.887 0.077 3.396506e-196
Arhgef40   1.390884e-198  0.9331330 0.850 0.067 2.341692e-194
Dcbld2     1.558435e-198  0.9227289 0.812 0.059 2.623781e-194
Ereg       2.474331e-198  0.7875364 0.600 0.025 4.165783e-194
Flnb       6.827685e-198  1.4673833 0.931 0.089 1.149509e-193
Hspg2      1.217089e-197  1.0573382 0.831 0.062 2.049090e-193
Ltbp3      2.911937e-197  1.6636338 0.931 0.092 4.902537e-193
Pkd2       2.743820e-196  0.8123280 0.744 0.049 4.619495e-192
Nppb       1.492874e-195  1.9230758 0.700 0.042 2.513402e-191
Gjb4       9.401103e-194  0.4910232 0.531 0.018 1.582770e-189
Frmd4a     1.872328e-193  1.4720788 0.944 0.098 3.152252e-189
Flrt3      1.809570e-192  0.5661047 0.494 0.013 3.046592e-188
Slc35e4    2.748369e-192  0.7989479 0.781 0.056 4.627154e-188
Col4a1     9.006539e-191  0.9790314 0.894 0.076 1.516341e-186
```

**Cluster 7 neutrophils**

```
cluster.markers <- FindMarkers(danx, ident.1 = 7, min.pct = 0.25)
  |++++++++++++++++++++++++++++++++++++++++++++++++++| 100% elapsed=15s
> head(cluster.markers, n = 100)
                       p_val avg_log2FC pct.1 pct.2     p_val_adj
Cxcr2           0.000000e+00   3.301580 0.788 0.001  0.000000e+00
Hdc             0.000000e+00   3.241031 0.689 0.006  0.000000e+00
S100a9          0.000000e+00   8.891881 0.887 0.033  0.000000e+00
Csf3r           0.000000e+00   3.568157 0.868 0.015  0.000000e+00
Wfdc21          0.000000e+00   4.278426 0.669 0.006  0.000000e+00
Stfa2l1         0.000000e+00   3.617743 0.543 0.000  0.000000e+00
Retnlg          0.000000e+00   6.815801 0.675 0.011  0.000000e+00
Cxcl2          8.310141e-304   6.007811 0.901 0.042 1.399095e-299
Mmp9           1.319218e-294   3.371336 0.722 0.018 2.221035e-290
Lcn2           3.671810e-275   4.208900 0.642 0.014 6.181860e-271
S100a8         9.917541e-264   8.500034 0.901 0.060 1.669717e-259
Clec4e         2.662691e-263   3.237590 0.675 0.020 4.482907e-259
Cstdc4         7.920616e-258   4.088434 0.470 0.002 1.333515e-253
Lrg1           8.293291e-252   2.785224 0.570 0.011 1.396259e-247
Asprv1         3.209148e-246   2.937711 0.457 0.002 5.402921e-242
Slpi           4.630684e-246   3.324865 0.728 0.031 7.796220e-242
G0s2           3.439697e-243   4.861974 0.550 0.010 5.791074e-239
Clec4d         1.860095e-229   2.999660 0.702 0.032 3.131656e-225
Hp             1.040243e-213   2.973029 0.636 0.027 1.751353e-209
Acod1          1.192564e-206   2.507072 0.417 0.004 2.007801e-202
Fpr2           5.733389e-204   1.942639 0.483 0.010 9.652734e-200
Il1f9          1.731623e-201   1.408045 0.358 0.001 2.915360e-197
```

| | | | | | |
|---|---|---|---|---|---|
| Il1r2 | 8.071814e-196 | 3.168504 | 0.722 | 0.048 | 1.358971e-191 |
| Trem1 | 1.208273e-194 | 2.545292 | 0.629 | 0.032 | 2.034248e-190 |
| Stfa2 | 6.578978e-194 | 2.670737 | 0.331 | 0.000 | 1.107637e-189 |
| Ifitm1 | 3.255564e-189 | 4.234558 | 0.894 | 0.094 | 5.481067e-185 |
| Slfn4 | 6.189358e-188 | 1.864329 | 0.391 | 0.005 | 1.042040e-183 |
| Chil1 | 2.610615e-187 | 1.463286 | 0.358 | 0.002 | 4.395232e-183 |
| Mxd1 | 2.443809e-183 | 3.123470 | 0.828 | 0.085 | 4.114397e-179 |
| Alox5ap | 7.789460e-180 | 3.531527 | 0.834 | 0.086 | 1.311434e-175 |
| Arg2 | 1.942139e-179 | 1.501150 | 0.371 | 0.004 | 3.269786e-175 |
| Cd300lf | 2.634680e-176 | 2.518634 | 0.702 | 0.054 | 4.435748e-172 |
| Mcemp1 | 8.420277e-165 | 2.009582 | 0.530 | 0.026 | 1.417638e-160 |
| Mmp8 | 1.320122e-164 | 3.165060 | 0.384 | 0.008 | 2.222558e-160 |
| Il1b | 1.153754e-161 | 5.279506 | 0.854 | 0.114 | 1.942461e-157 |
| Hcar2 | 3.424737e-160 | 2.312555 | 0.411 | 0.011 | 5.765887e-156 |
| Lilrb4a | 2.173690e-159 | 2.594476 | 0.781 | 0.085 | 3.659625e-155 |
| Cd33 | 5.761241e-154 | 2.731565 | 0.715 | 0.073 | 9.699626e-150 |
| Ccr1 | 1.950513e-153 | 2.795680 | 0.815 | 0.099 | 3.283884e-149 |
| Lilr4b | 4.801165e-153 | 2.859834 | 0.682 | 0.063 | 8.083241e-149 |
| Mirt2 | 4.135743e-147 | 1.179002 | 0.258 | 0.000 | 6.962936e-143 |
| Cd177 | 1.637688e-143 | 1.585569 | 0.298 | 0.003 | 2.757211e-139 |
| Pglyrp1 | 1.955334e-139 | 2.836148 | 0.636 | 0.060 | 3.292001e-135 |
| Osm | 7.198086e-127 | 3.276632 | 0.517 | 0.039 | 1.211870e-122 |
| Tyrobp | 2.882495e-125 | 2.964276 | 1.000 | 0.251 | 4.852969e-121 |
| Nfam1 | 8.989679e-124 | 2.229136 | 0.616 | 0.064 | 1.513502e-119 |
| Cd300ld | 1.532886e-122 | 1.903398 | 0.523 | 0.041 | 2.580767e-118 |
| Ankrd33b | 6.178968e-120 | 1.300985 | 0.311 | 0.008 | 1.040291e-115 |
| C5ar1 | 2.127145e-117 | 2.011470 | 0.609 | 0.065 | 3.581262e-113 |
| Bst1 | 1.165424e-115 | 1.728576 | 0.437 | 0.028 | 1.962107e-111 |
| Egr1 | 2.525292e-114 | 3.918077 | 0.861 | 0.190 | 4.251581e-110 |
| Slc40a1 | 7.623931e-112 | 1.655084 | 0.364 | 0.017 | 1.283565e-107 |
| Cd14 | 1.347845e-109 | 3.538852 | 0.702 | 0.111 | 2.269232e-105 |
| Gsr | 3.103168e-109 | 2.682593 | 0.841 | 0.188 | 5.224494e-105 |
| Slc7a11 | 2.214032e-108 | 2.193829 | 0.437 | 0.031 | 3.727543e-104 |
| Pygl | 2.270952e-108 | 2.083951 | 0.556 | 0.059 | 3.823375e-104 |
| Gda | 4.639159e-108 | 2.621174 | 0.742 | 0.126 | 7.810488e-104 |
| Chil3 | 6.305948e-108 | 2.954012 | 0.397 | 0.023 | 1.061669e-103 |
| Trim30b | 2.343961e-107 | 1.631358 | 0.437 | 0.031 | 3.946293e-103 |
| Msrb1 | 1.290469e-104 | 3.044801 | 0.921 | 0.272 | 2.172633e-100 |
| Ccrl2 | 1.128512e-103 | 2.987967 | 0.497 | 0.047 | 1.899963e-99 |
| Dusp1 | 7.155050e-101 | 4.028283 | 0.967 | 0.367 | 1.204624e-96 |
| Wfdc17 | 1.989066e-98 | 4.272861 | 0.788 | 0.173 | 3.348791e-94 |
| Gm9733 | 5.396365e-98 | 1.386217 | 0.325 | 0.016 | 9.085320e-94 |
| Ptafr | 1.073651e-96 | 2.048403 | 0.437 | 0.037 | 1.807598e-92 |
| Cebpb | 2.192310e-96 | 2.227762 | 0.682 | 0.116 | 3.690973e-92 |
| Itgam | 7.784143e-96 | 2.035519 | 0.589 | 0.078 | 1.310538e-91 |
| Fcer1g | 1.451826e-95 | 2.161304 | 0.980 | 0.241 | 2.444295e-91 |
| Grina | 2.379462e-93 | 2.863083 | 0.821 | 0.223 | 4.006063e-89 |
| Ifitm6 | 1.151018e-92 | 3.295757 | 0.384 | 0.028 | 1.937854e-88 |
| Mcl1 | 1.539717e-89 | 2.798343 | 0.993 | 0.551 | 2.592267e-85 |
| Srgn | 4.455917e-89 | 3.065175 | 0.987 | 0.581 | 7.501981e-85 |
| F630028O10Rik | 2.638218e-88 | 1.479152 | 0.285 | 0.013 | 4.441704e-84 |
| Siglece | 1.024501e-87 | 1.329498 | 0.344 | 0.023 | 1.724850e-83 |

```
Fgr           2.093674e-85   1.716405 0.517 0.065  3.524910e-81
Ppp1r3b       2.351442e-85   1.297582 0.351 0.025  3.958887e-81
Pim1          4.046354e-85   3.137019 0.901 0.335  6.812441e-81
Dhrs9         1.418936e-84   0.942291 0.272 0.012  2.388921e-80
Rps20         2.020386e-84  -3.006973 0.629 0.993  3.401522e-80
Ier3          6.383253e-84   3.476357 0.795 0.211  1.074685e-79
Il18rap       2.340067e-83   1.650678 0.457 0.052  3.939736e-79
Rpl13         1.459878e-82  -2.587860 0.828 0.994  2.457851e-78
Rps4x         3.450687e-81  -2.852966 0.517 0.981  5.809577e-77
Pirb          1.933209e-80   1.777327 0.576 0.092  3.254751e-76
Eef1a1        7.582466e-80  -2.210359 0.834 0.996  1.276584e-75
Rpl12         1.840795e-79  -3.072607 0.530 0.980  3.099163e-75
Rpsa          2.169931e-79  -2.645024 0.623 0.981  3.653296e-75
Ifitm2        2.183633e-79   2.788079 0.901 0.344  3.676364e-75
Rpl3          2.674553e-79  -2.888962 0.325 0.962  4.502877e-75
Rps8          3.421192e-79  -2.331277 0.735 0.988  5.759919e-75
Rpl23         4.142961e-79  -2.243854 0.894 0.992  6.975089e-75
Rps24         8.552865e-79  -2.697470 0.881 0.994  1.439960e-74
Rpl10a        8.728597e-79  -2.893325 0.305 0.963  1.469547e-74
Rpl32         1.499487e-78  -2.571325 0.702 0.984  2.524537e-74
Lst1          1.796817e-78   2.309246 0.656 0.125  3.025122e-74
Rpl13a        2.768446e-78  -2.419251 0.715 0.985  4.660956e-74
Csf2rb        3.085494e-78   1.676154 0.556 0.084  5.194738e-74
Rpl19         3.191713e-78  -2.306523 0.795 0.989  5.373569e-74
Fbxl5         3.198497e-78   2.854528 0.722 0.191  5.384989e-74
Rplp1         1.539616e-77  -2.260156 0.848 0.997  2.592098e-73
```

**Cluster 8 endothelial cells**

```
cluster.markers <- FindMarkers(danx, ident.1 = 8, min.pct = 0.25)
  |+++++++++++++++++++++++++++++++++++++++++++++++++++| 100% elapsed=18s
> head(cluster.markers, n = 100)
                p_val avg_log2FC pct.1 pct.2    p_val_adj
Sulf1      0.000000e+00  1.4335735 0.568 0.000  0.000000e+00
Col3a1     0.000000e+00  6.7762674 0.986 0.024  0.000000e+00
Col5a2     0.000000e+00  3.2652187 0.865 0.003  0.000000e+00
Prelp      0.000000e+00  1.9060909 0.622 0.002  0.000000e+00
Dpt        0.000000e+00  4.0216061 0.703 0.001  0.000000e+00
Cd34       0.000000e+00  2.6527245 0.743 0.003  0.000000e+00
Entpd2     0.000000e+00  1.3589611 0.527 0.000  0.000000e+00
Scn7a      0.000000e+00  1.7381422 0.554 0.000  0.000000e+00
Serping1   0.000000e+00  4.3035024 0.851 0.002  0.000000e+00
Fbn1       0.000000e+00  3.8687759 0.878 0.004  0.000000e+00
Cpxm1      0.000000e+00  3.6923896 0.851 0.005  0.000000e+00
Tshz2      0.000000e+00  1.0846617 0.622 0.003  0.000000e+00
Sfrp2      0.000000e+00  3.3119463 0.662 0.003  0.000000e+00
Ctsk       0.000000e+00  2.3830291 0.743 0.004  0.000000e+00
Gstm2      0.000000e+00  1.2595338 0.689 0.004  0.000000e+00
Svep1      0.000000e+00  2.5716228 0.595 0.000  0.000000e+00
Mfap2      0.000000e+00  1.4666867 0.649 0.001  0.000000e+00
Mmp23      0.000000e+00  1.2601177 0.608 0.003  0.000000e+00
Sod3       0.000000e+00  1.5327179 0.635 0.001  0.000000e+00
Adgrd1     0.000000e+00  1.5766964 0.716 0.000  0.000000e+00
```

| | | | | | |
|---|---|---|---|---|---|
| Col1a2 | 0.000000e+00 | 5.6385465 | 0.986 | 0.010 | 0.000000e+00 |
| Rarres2 | 0.000000e+00 | 3.8729920 | 0.838 | 0.004 | 0.000000e+00 |
| Mfap5 | 0.000000e+00 | 3.5775689 | 0.784 | 0.004 | 0.000000e+00 |
| C1s1 | 0.000000e+00 | 3.2238641 | 0.838 | 0.009 | 0.000000e+00 |
| Fxyd1 | 0.000000e+00 | 1.7071344 | 0.662 | 0.001 | 0.000000e+00 |
| Prss23 | 0.000000e+00 | 2.9173123 | 0.811 | 0.008 | 0.000000e+00 |
| Serpinh1 | 0.000000e+00 | 3.4627089 | 0.892 | 0.008 | 0.000000e+00 |
| Nnmt | 0.000000e+00 | 1.4640802 | 0.635 | 0.000 | 0.000000e+00 |
| Islr | 0.000000e+00 | 1.8762488 | 0.608 | 0.001 | 0.000000e+00 |
| Loxl1 | 0.000000e+00 | 2.8572041 | 0.838 | 0.002 | 0.000000e+00 |
| Rbms3 | 0.000000e+00 | 1.5274412 | 0.689 | 0.003 | 0.000000e+00 |
| Clec3b | 0.000000e+00 | 3.5134188 | 0.703 | 0.003 | 0.000000e+00 |
| Col6a2 | 0.000000e+00 | 2.2236102 | 0.770 | 0.008 | 0.000000e+00 |
| Lum | 0.000000e+00 | 3.5710096 | 0.797 | 0.001 | 0.000000e+00 |
| Aebp1 | 0.000000e+00 | 3.6240502 | 0.892 | 0.007 | 0.000000e+00 |
| Grb10 | 0.000000e+00 | 1.2243257 | 0.608 | 0.002 | 0.000000e+00 |
| Efemp1 | 0.000000e+00 | 3.2568860 | 0.757 | 0.000 | 0.000000e+00 |
| Slit3 | 0.000000e+00 | 1.6580718 | 0.662 | 0.000 | 0.000000e+00 |
| Gfpt2 | 0.000000e+00 | 1.3986686 | 0.581 | 0.001 | 0.000000e+00 |
| Col1a1 | 0.000000e+00 | 5.5226741 | 0.959 | 0.021 | 0.000000e+00 |
| Cavin1 | 0.000000e+00 | 2.5260908 | 0.851 | 0.008 | 0.000000e+00 |
| Mrc2 | 0.000000e+00 | 0.7856092 | 0.568 | 0.001 | 0.000000e+00 |
| Cygb | 0.000000e+00 | 1.9446561 | 0.608 | 0.002 | 0.000000e+00 |
| Serpina3n | 0.000000e+00 | 2.7849537 | 0.716 | 0.001 | 0.000000e+00 |
| Meg3 | 0.000000e+00 | 3.0319598 | 0.676 | 0.001 | 0.000000e+00 |
| Rian | 0.000000e+00 | 1.3601116 | 0.554 | 0.001 | 0.000000e+00 |
| Ogn | 0.000000e+00 | 2.6181922 | 0.649 | 0.001 | 0.000000e+00 |
| Gas1 | 0.000000e+00 | 2.4055894 | 0.662 | 0.004 | 0.000000e+00 |
| Scara3 | 0.000000e+00 | 1.9402883 | 0.595 | 0.001 | 0.000000e+00 |
| Loxl2 | 0.000000e+00 | 1.5386423 | 0.703 | 0.001 | 0.000000e+00 |
| Fbln1 | 0.000000e+00 | 1.8584934 | 0.716 | 0.001 | 0.000000e+00 |
| Ccdc80 | 0.000000e+00 | 2.7702875 | 0.824 | 0.008 | 0.000000e+00 |
| Abi3bp | 0.000000e+00 | 2.3490779 | 0.608 | 0.001 | 0.000000e+00 |
| Vgll3 | 0.000000e+00 | 1.8652341 | 0.730 | 0.006 | 0.000000e+00 |
| Adamts5 | 0.000000e+00 | 3.0552373 | 0.730 | 0.003 | 0.000000e+00 |
| Fndc1 | 0.000000e+00 | 1.9009050 | 0.568 | 0.001 | 0.000000e+00 |
| Thbs2 | 0.000000e+00 | 2.3914495 | 0.662 | 0.001 | 0.000000e+00 |
| Tnxb | 0.000000e+00 | 2.8053319 | 0.689 | 0.002 | 0.000000e+00 |
| Lox | 0.000000e+00 | 3.1870174 | 0.797 | 0.001 | 0.000000e+00 |
| Cd248 | 0.000000e+00 | 2.7168927 | 0.703 | 0.001 | 0.000000e+00 |
| Il33 | 0.000000e+00 | 1.6863847 | 0.581 | 0.001 | 0.000000e+00 |
| Srpx | 0.000000e+00 | 1.2771288 | 0.595 | 0.000 | 0.000000e+00 |
| Gpc3 | 0.000000e+00 | 1.5237458 | 0.608 | 0.002 | 0.000000e+00 |
| Tmem45a | 5.860106e-307 | 0.9368072 | 0.514 | 0.000 | 9.866074e-303 |
| Adamts2 | 1.943753e-306 | 2.2669786 | 0.757 | 0.010 | 3.272503e-302 |
| Lgi2 | 5.272895e-306 | 1.8387371 | 0.595 | 0.003 | 8.877446e-302 |
| Nid1 | 2.940387e-301 | 3.3292312 | 0.797 | 0.013 | 4.950435e-297 |
| Adamts15 | 4.828784e-299 | 1.3956653 | 0.514 | 0.000 | 8.129741e-295 |
| Bgn | 3.508549e-296 | 3.9082547 | 0.892 | 0.020 | 5.906993e-292 |
| Adamtsl1 | 3.354628e-292 | 0.9274473 | 0.527 | 0.001 | 5.647852e-288 |
| Fkbp10 | 1.259433e-291 | 0.9673931 | 0.527 | 0.001 | 2.120381e-287 |
| Hoxc8 | 1.044335e-290 | 0.7820579 | 0.486 | 0.000 | 1.758243e-286 |

```
C1ra      1.217190e-288  2.3306742 0.797 0.014 2.049260e-284
Lrrc17    1.366975e-282  1.3151188 0.473 0.000 2.301440e-278
Igsf10    2.189349e-282  1.0875626 0.486 0.000 3.685988e-278
Ccl11     3.441974e-275  1.9508656 0.486 0.001 5.794908e-271
Ndn       1.164491e-274  0.7041430 0.486 0.001 1.960537e-270
Spon2     2.505059e-274  1.0169226 0.473 0.000 4.217517e-270
Dcn       8.159521e-274  6.2438555 0.919 0.027 1.373737e-269
Timp3     4.273175e-273  2.8319635 0.595 0.005 7.194317e-269
Thbs3     4.955896e-273  1.4995581 0.595 0.005 8.343746e-269
Medag     8.579973e-273  1.5457798 0.689 0.010 1.444524e-268
Nsg1      3.911830e-270  1.2530440 0.514 0.002 6.585957e-266
Fxyd6     5.224634e-269  1.3773715 0.500 0.001 8.796194e-265
Sfrp4     6.046832e-268  3.0475060 0.486 0.001 1.018045e-263
Prg4      6.147157e-268  2.1018016 0.541 0.003 1.034935e-263
Ptgis     6.360328e-268  1.7189387 0.486 0.001 1.070825e-263
Bicc1     7.197644e-267  1.7363303 0.703 0.011 1.211795e-262
Dpep1     2.410728e-266  1.2953583 0.459 0.000 4.058702e-262
Lama4     4.550189e-264  1.3709379 0.635 0.007 7.660698e-260
Col6a3    5.339488e-264  2.6571697 0.784 0.017 8.989561e-260
Gpx7      2.150285e-261  0.6715112 0.500 0.002 3.620219e-257
Pamr1     2.414124e-258  1.2254912 0.446 0.000 4.064419e-254
Col6a1    2.206546e-256  2.9075347 0.838 0.022 3.714941e-252
Lbp       4.031535e-255  2.6558986 0.595 0.006 6.787493e-251
Serpinf1  1.127474e-254  2.7362840 0.770 0.017 1.898216e-250
Eln       2.117488e-252  1.5214323 0.473 0.001 3.565003e-248
Vcan      5.072300e-252  2.0332087 0.716 0.013 8.539724e-248
Chl1      2.322837e-250  1.3581847 0.432 0.000 3.910728e-246
Osmr      2.685397e-250  1.0863169 0.581 0.006 4.521134e-246
```

**Cluster 9 Fibroblasts**

```
cluster.markers <- FindMarkers(danx, ident.1 = 9, min.pct = 0.25)
  |++++++++++++++++++++++++++++++++++++++++++++++++++| 100% elapsed=22s
> head(cluster.markers, n = 100)
                 p_val avg_log2FC pct.1 pct.2     p_val_adj
Col4a3       0.000000e+00  0.7230677 0.725 0.007  0.000000e+00
Epha7        0.000000e+00  0.4976842 0.623 0.002  0.000000e+00
Tbx2        7.522537e-306  1.0695632 0.812 0.011 1.266494e-301
Acsl6       9.380088e-295  0.3820855 0.493 0.000 1.579232e-290
Eda2r       4.517953e-274  0.4787770 0.652 0.007 7.606426e-270
Gm43113     8.805879e-274  0.4499067 0.565 0.003 1.482558e-269
Gm43728     3.512266e-270  0.6319708 0.710 0.010 5.913251e-266
Gm43112     2.488658e-259  0.5467492 0.580 0.005 4.189904e-255
Cdh13       1.033197e-243  0.3255644 0.536 0.004 1.739490e-239
Phactr1     3.633069e-243  0.6933797 0.725 0.014 6.116635e-239
Tafa1       1.029129e-240  0.3537719 0.478 0.002 1.732642e-236
Gm43111     2.956446e-232  0.6693282 0.594 0.008 4.977472e-228
Nes         4.173032e-230  1.3165030 0.928 0.032 7.025716e-226
Tspan7      2.205343e-228  0.9992850 0.797 0.021 3.712916e-224
Fermt1      4.992820e-223  0.7063801 0.696 0.015 8.405911e-219
Col8a1      6.563702e-211  1.2205713 0.623 0.011 1.105065e-206
Ociad2      2.855545e-206  0.4292836 0.609 0.011 4.807595e-202
Gm43727     3.909556e-205  0.6455917 0.536 0.007 6.582128e-201
Gm10484     1.302231e-202  0.3658995 0.478 0.005 2.192435e-198
```

| | | | | | |
|---|---|---|---|---|---|
| Naalad2 | 6.520005e-200 | 0.3929533 | 0.536 | 0.008 | 1.097708e-195 |
| Sorbs2 | 1.237954e-196 | 0.7631532 | 0.667 | 0.016 | 2.084219e-192 |
| Edil3 | 1.412816e-191 | 0.4373316 | 0.551 | 0.010 | 2.378617e-187 |
| Gpc4 | 3.705230e-190 | 0.4190561 | 0.551 | 0.010 | 6.238125e-186 |
| Ngf | 9.236070e-189 | 0.6805670 | 0.725 | 0.023 | 1.554985e-184 |
| Prokr2 | 2.325181e-188 | 0.3491865 | 0.435 | 0.004 | 3.914675e-184 |
| Sema5a | 1.903647e-182 | 2.2327903 | 0.928 | 0.048 | 3.204980e-178 |
| Tenm3 | 3.603906e-180 | 1.3794103 | 0.884 | 0.043 | 6.067537e-176 |
| Col4a4 | 4.233993e-179 | 0.3981527 | 0.522 | 0.009 | 7.128351e-175 |
| Pdgfrl | 2.711203e-178 | 0.5131323 | 0.449 | 0.006 | 4.564581e-174 |
| Gm43623 | 7.186960e-175 | 0.3546871 | 0.493 | 0.008 | 1.209997e-170 |
| Nrn1 | 7.991284e-173 | 0.9134940 | 0.609 | 0.016 | 1.345413e-168 |
| Masp1 | 2.532708e-172 | 0.7564859 | 0.667 | 0.021 | 4.264066e-168 |
| Mpzl1 | 1.676859e-170 | 1.3130816 | 0.986 | 0.060 | 2.823160e-166 |
| Disp2 | 2.047403e-170 | 0.4188342 | 0.609 | 0.017 | 3.447008e-166 |
| Cldn12 | 1.404723e-168 | 1.4705870 | 0.870 | 0.044 | 2.364992e-164 |
| Pex1 | 6.932679e-168 | 1.0793341 | 0.855 | 0.041 | 1.167186e-163 |
| Serpine2 | 6.873451e-167 | 2.3901991 | 0.681 | 0.023 | 1.157214e-162 |
| Chst2 | 5.735701e-166 | 0.5031274 | 0.623 | 0.018 | 9.656625e-162 |
| Lingo1 | 3.793237e-162 | 0.2586262 | 0.362 | 0.003 | 6.386294e-158 |
| 2510017J16Rik | 2.917526e-161 | 0.6509473 | 0.565 | 0.015 | 4.911947e-157 |
| Srgap3 | 4.696925e-161 | 0.7421138 | 0.797 | 0.037 | 7.907744e-157 |
| Klhdc8a | 2.196034e-159 | 0.7425108 | 0.725 | 0.029 | 3.697243e-155 |
| Cd109 | 9.820392e-159 | 1.1114420 | 0.913 | 0.053 | 1.653361e-154 |
| Ibsp | 3.969527e-154 | 3.5842129 | 0.783 | 0.040 | 6.683096e-150 |
| Adgrg3 | 7.684864e-154 | 0.7208587 | 0.667 | 0.024 | 1.293824e-149 |
| Ank3 | 4.832928e-152 | 0.9390043 | 0.957 | 0.061 | 8.136718e-148 |
| Gdpd5 | 6.715319e-152 | 1.3851501 | 0.928 | 0.061 | 1.130591e-147 |
| Col7a1 | 1.589251e-151 | 1.1968641 | 0.957 | 0.061 | 2.675662e-147 |
| Gm17590 | 3.523869e-151 | 0.2834897 | 0.377 | 0.004 | 5.932786e-147 |
| Fzd1 | 7.727660e-151 | 1.5679986 | 0.826 | 0.046 | 1.301029e-146 |
| 1700109H08Rik | 7.417603e-149 | 0.4306067 | 0.507 | 0.012 | 1.248828e-144 |
| Akap12 | 7.940667e-149 | 1.1716861 | 0.739 | 0.034 | 1.336891e-144 |
| Psrc1 | 2.015702e-148 | 0.3851642 | 0.420 | 0.007 | 3.393637e-144 |
| Tspan9 | 1.247125e-146 | 0.8368776 | 0.884 | 0.054 | 2.099660e-142 |
| Tspan8 | 3.589393e-142 | 0.4842769 | 0.493 | 0.013 | 6.043101e-138 |
| Dennd2a | 5.366566e-142 | 0.3552877 | 0.551 | 0.017 | 9.035151e-138 |
| Grem1 | 3.327764e-141 | 1.6126170 | 0.841 | 0.052 | 5.602623e-137 |
| Pdgfrb | 3.882045e-138 | 0.5202530 | 0.565 | 0.019 | 6.535811e-134 |
| Cavin3 | 5.009175e-138 | 0.4604431 | 0.725 | 0.034 | 8.433447e-134 |
| Rbm48 | 2.581397e-136 | 1.1475196 | 0.855 | 0.053 | 4.346040e-132 |
| Mcam | 3.766162e-136 | 0.5586803 | 0.391 | 0.007 | 6.340711e-132 |
| Ndst3 | 5.682472e-135 | 0.6516902 | 0.681 | 0.033 | 9.567010e-131 |
| 4833413G10Rik | 1.541926e-134 | 0.4845746 | 0.493 | 0.014 | 2.595987e-130 |
| Pxdn | 1.672015e-133 | 1.4725472 | 0.942 | 0.073 | 2.815004e-129 |
| Itga3 | 1.067514e-132 | 1.1228672 | 0.928 | 0.066 | 1.797267e-128 |
| Ddr2 | 1.475235e-132 | 0.4378221 | 0.696 | 0.033 | 2.483706e-128 |
| Rbpms2 | 2.173718e-132 | 0.9852336 | 0.855 | 0.058 | 3.659672e-128 |
| D630045J12Rik | 7.204487e-132 | 0.2777600 | 0.449 | 0.011 | 1.212947e-127 |
| Cdh2 | 5.612466e-131 | 1.0730211 | 0.884 | 0.064 | 9.449147e-127 |
| Gpc1 | 2.035809e-130 | 1.4260649 | 0.899 | 0.068 | 3.427488e-126 |
| Nptx1 | 2.867170e-129 | 0.5298506 | 0.449 | 0.011 | 4.827167e-125 |
| Agrn | 1.694443e-128 | 0.6147046 | 0.696 | 0.036 | 2.852764e-124 |
| Cst6 | 4.138385e-128 | 0.6266173 | 0.681 | 0.035 | 6.967386e-124 |

| | | | | | |
|---|---|---|---|---|---|
| C1qtnf12 | 2.243517e-127 | 0.3289623 | 0.551 | 0.020 | 3.777185e-123 |
| Atg9b | 2.378931e-127 | 0.5252725 | 0.594 | 0.025 | 4.005168e-123 |
| Col27a1 | 4.153947e-127 | 0.5089348 | 0.580 | 0.023 | 6.993586e-123 |
| Adgrl2 | 7.582109e-127 | 0.5116929 | 0.435 | 0.011 | 1.276524e-122 |
| Pros1 | 1.102005e-126 | 0.7340433 | 0.797 | 0.048 | 1.855335e-122 |
| Cacna2d1 | 2.367485e-126 | 0.7617239 | 0.797 | 0.052 | 3.985898e-122 |
| Ptprn | 2.707732e-126 | 1.1451030 | 0.841 | 0.057 | 4.558738e-122 |
| Nectin3 | 3.376276e-126 | 0.8382153 | 0.826 | 0.056 | 5.684298e-122 |
| Col4a2 | 5.227043e-126 | 1.0980959 | 0.942 | 0.075 | 8.800250e-122 |
| Samd5 | 5.855151e-125 | 0.2534229 | 0.348 | 0.006 | 9.857732e-121 |
| Greb1 | 6.439267e-124 | 0.6418137 | 0.667 | 0.034 | 1.084115e-119 |
| Ano1 | 1.657502e-122 | 1.5463865 | 0.971 | 0.085 | 2.790570e-118 |
| Gpr149 | 1.313980e-121 | 0.2773102 | 0.348 | 0.006 | 2.212217e-117 |
| Unc5b | 3.103438e-121 | 1.1803327 | 0.928 | 0.076 | 5.224948e-117 |
| Piezo2 | 4.632115e-121 | 1.8777119 | 0.957 | 0.084 | 7.798629e-117 |
| Gprc5a | 5.595960e-121 | 0.8458766 | 0.783 | 0.051 | 9.421357e-117 |
| Zfp462 | 1.445562e-120 | 0.4733694 | 0.667 | 0.035 | 2.433749e-116 |
| Nav3 | 4.951815e-120 | 0.6318918 | 0.797 | 0.054 | 8.336876e-116 |
| Morc4 | 1.156045e-119 | 0.4349627 | 0.667 | 0.036 | 1.946318e-115 |
| Plxna1 | 1.880869e-119 | 0.7811181 | 0.826 | 0.060 | 3.166631e-115 |
| Cdr2l | 2.507754e-119 | 0.4750741 | 0.696 | 0.039 | 4.222055e-115 |
| Ltbp4 | 2.629938e-119 | 1.0146420 | 0.812 | 0.058 | 4.427763e-115 |
| Col18a1 | 3.261583e-119 | 2.5227649 | 1.000 | 0.103 | 5.491202e-115 |
| Loxl3 | 7.901678e-119 | 0.6913770 | 0.797 | 0.054 | 1.330327e-114 |
| Plod2 | 1.626028e-118 | 1.7308475 | 1.000 | 0.098 | 2.737581e-114 |
| Tinagl1 | 1.641555e-118 | 0.8436353 | 0.812 | 0.057 | 2.763722e-114 |
| P3h3 | 6.272223e-118 | 0.4939922 | 0.536 | 0.021 | 1.055991e-113 |

**Cluster 10 Mesenchymal progenitors**

```
cluster.markers <- FindMarkers(danx, ident.1 = 10, min.pct = 0.25)
  |++++++++++++++++++++++++++++++++++++++++++++++++++| 100% elapsed=11s
> head(cluster.markers, n = 100)
```

| | p_val | avg_log2FC | pct.1 | pct.2 | p_val_adj |
|---|---|---|---|---|---|
| Birc5 | 3.753883e-132 | 2.3077276 | 0.742 | 0.041 | 6.320037e-128 |
| Ccna2 | 3.603986e-105 | 1.2892541 | 0.470 | 0.018 | 6.067671e-101 |
| Pclaf | 1.498704e-97 | 2.6083450 | 0.682 | 0.052 | 2.523219e-93 |
| Ube2c | 2.790596e-93 | 2.1174373 | 0.576 | 0.036 | 4.698248e-89 |
| Hist1h2ap | 2.591653e-89 | 2.6884903 | 0.652 | 0.053 | 4.363307e-85 |
| Cdca8 | 1.897902e-84 | 1.6313352 | 0.470 | 0.025 | 3.195309e-80 |
| Hist1h1b | 5.608513e-77 | 2.0283746 | 0.455 | 0.027 | 9.442493e-73 |
| Hist1h2ae | 3.672976e-76 | 2.0976746 | 0.545 | 0.042 | 6.183822e-72 |
| Cdca3 | 9.502625e-74 | 1.5583323 | 0.424 | 0.024 | 1.599862e-69 |
| Aurkb | 1.723646e-72 | 1.1971640 | 0.333 | 0.013 | 2.901931e-68 |
| Mki67 | 1.445663e-71 | 1.9728857 | 0.455 | 0.029 | 2.433918e-67 |
| Top2a | 4.136779e-70 | 2.2238921 | 0.667 | 0.073 | 6.964682e-66 |
| Kif22 | 1.359009e-68 | 1.0382725 | 0.303 | 0.011 | 2.288028e-64 |
| Nusap1 | 4.909682e-66 | 1.1640561 | 0.318 | 0.014 | 8.265941e-62 |
| Cenpe | 4.094797e-58 | 0.8608544 | 0.303 | 0.014 | 6.894000e-54 |
| Knl1 | 6.534547e-57 | 0.9489863 | 0.318 | 0.017 | 1.100156e-52 |
| Bub1 | 6.464861e-56 | 0.7508618 | 0.258 | 0.010 | 1.088424e-51 |
| Tpx2 | 1.107190e-54 | 1.3155599 | 0.409 | 0.031 | 1.864066e-50 |
| Stmn1 | 6.498378e-54 | 2.5791525 | 0.818 | 0.171 | 1.094067e-49 |
| Spc24 | 1.299873e-48 | 0.9981040 | 0.333 | 0.023 | 2.188467e-44 |

| | | | | | |
|---|---|---|---|---|---|
| Racgap1 | 5.481379e-48 | 1.3001905 | 0.455 | 0.046 | 9.228450e-44 |
| Pbk | 6.415074e-48 | 1.0693481 | 0.258 | 0.013 | 1.080042e-43 |
| Kif11 | 4.265034e-46 | 0.9757336 | 0.258 | 0.014 | 7.180612e-42 |
| Ccnb1 | 5.459131e-46 | 0.9032933 | 0.273 | 0.016 | 9.190993e-42 |
| Kif15 | 1.478593e-44 | 0.8286229 | 0.273 | 0.016 | 2.489359e-40 |
| Cenpm | 3.283192e-43 | 0.6799864 | 0.258 | 0.015 | 5.527582e-39 |
| Tk1 | 1.019845e-42 | 1.4378491 | 0.379 | 0.036 | 1.717012e-38 |
| Ccnb2 | 5.159955e-42 | 0.8386548 | 0.273 | 0.017 | 8.687300e-38 |
| Bub1b | 1.134198e-41 | 0.8170572 | 0.258 | 0.016 | 1.909535e-37 |
| Tyms | 1.377777e-41 | 1.7654949 | 0.424 | 0.048 | 2.319626e-37 |
| Cenpf | 4.239479e-40 | 1.0059111 | 0.258 | 0.016 | 7.137587e-36 |
| Kif23 | 4.406599e-37 | 1.0289847 | 0.364 | 0.037 | 7.418950e-33 |
| Hmgb2 | 4.655295e-36 | 2.7193193 | 0.924 | 0.441 | 7.837655e-32 |
| H2afz | 3.778436e-34 | 2.1932427 | 0.970 | 0.629 | 6.361375e-30 |
| Rrm2 | 2.646718e-29 | 1.0010551 | 0.288 | 0.030 | 4.456014e-25 |
| Tuba1b | 2.156312e-28 | 2.1070331 | 0.833 | 0.363 | 3.630366e-24 |
| Cdk1 | 2.905147e-28 | 1.1111430 | 0.409 | 0.064 | 4.891106e-24 |
| Bcl2a1a | 4.674409e-27 | 1.5890567 | 0.500 | 0.097 | 7.869835e-23 |
| Ptma | 8.112307e-27 | 1.3195346 | 1.000 | 0.944 | 1.365788e-22 |
| H2afx | 1.509892e-26 | 1.3577150 | 0.485 | 0.098 | 2.542054e-22 |
| Bcl2a1b | 1.961235e-26 | 1.6118321 | 0.803 | 0.253 | 3.301934e-22 |
| Smc2 | 3.227376e-25 | 1.2253800 | 0.424 | 0.078 | 5.433611e-21 |
| Tubb5 | 5.887850e-25 | 1.7766304 | 0.909 | 0.575 | 9.912784e-21 |
| C1qb | 1.646363e-24 | 1.9018242 | 0.652 | 0.175 | 2.771817e-20 |
| Cks2 | 5.136486e-24 | 1.4679807 | 0.561 | 0.141 | 8.647788e-20 |
| Gmnn | 9.395210e-24 | 1.0371835 | 0.424 | 0.080 | 1.581778e-19 |
| Aif1 | 1.154331e-23 | 1.8347477 | 0.636 | 0.181 | 1.943431e-19 |
| C1qa | 1.209539e-23 | 1.7461773 | 0.667 | 0.183 | 2.036380e-19 |
| Tmsb4x | 1.847064e-23 | 1.1665860 | 1.000 | 0.999 | 3.109716e-19 |
| Gatm | 4.044414e-22 | 1.8730907 | 0.621 | 0.198 | 6.809176e-18 |
| Bcl2a1d | 5.998963e-22 | 1.2403591 | 0.455 | 0.096 | 1.009985e-17 |
| Rrm1 | 2.597002e-21 | 1.3246373 | 0.455 | 0.102 | 4.372313e-17 |
| Cxcl16 | 5.421460e-21 | 1.5169828 | 0.500 | 0.120 | 9.127570e-17 |
| Msr1 | 1.035051e-20 | 1.3425079 | 0.455 | 0.100 | 1.742612e-16 |
| Cenpw | 1.122721e-20 | 0.8388917 | 0.303 | 0.047 | 1.890214e-16 |
| Malat1 | 4.706924e-20 | -2.5014918 | 0.348 | 0.831 | 7.924578e-16 |
| Rfc5 | 6.352104e-20 | 0.8219691 | 0.288 | 0.044 | 1.069440e-15 |
| Hist1h1e | 6.866738e-20 | 1.3470422 | 0.485 | 0.123 | 1.156084e-15 |
| Incenp | 1.107681e-18 | 0.9051065 | 0.303 | 0.053 | 1.864891e-14 |
| Lyz2 | 1.391515e-18 | 1.6150202 | 0.727 | 0.260 | 2.342755e-14 |
| Junb | 3.700952e-18 | -2.4769801 | 0.379 | 0.803 | 6.230923e-14 |
| C1qc | 4.880997e-18 | 1.6059380 | 0.561 | 0.165 | 8.217646e-14 |
| Hmgn2 | 1.301497e-17 | 1.3530090 | 0.561 | 0.184 | 2.191200e-13 |
| Lig1 | 1.331136e-17 | 0.8019928 | 0.318 | 0.060 | 2.241101e-13 |
| Slamf9 | 7.288100e-17 | 0.7921366 | 0.288 | 0.051 | 1.227024e-12 |
| Fcer1g | 1.077675e-16 | 1.3930470 | 0.727 | 0.271 | 1.814373e-12 |
| Pf4 | 3.559068e-16 | 1.6432260 | 0.424 | 0.111 | 5.992048e-12 |
| Atpif1 | 4.401824e-16 | 1.2807235 | 0.833 | 0.457 | 7.410911e-12 |
| Ran | 1.846479e-15 | 1.2745943 | 0.833 | 0.517 | 3.108732e-11 |
| Arg1 | 2.649943e-15 | 0.9797746 | 0.470 | 0.129 | 4.461444e-11 |
| Apoe | 3.132128e-15 | 1.2094098 | 0.788 | 0.373 | 5.273251e-11 |
| Rps27 | 7.303832e-15 | -1.5579985 | 0.485 | 0.849 | 1.229673e-10 |

| | | | | | |
|---|---|---|---|---|---|
| Apoc2 | 9.917725e-15 | 1.1834588 | 0.348 | 0.080 | 1.669748e-10 |
| Jund | 1.004742e-14 | -1.8492103 | 0.318 | 0.773 | 1.691583e-10 |
| Cox8a | 1.567473e-14 | 0.9283069 | 0.955 | 0.831 | 2.638997e-10 |
| Ppia | 4.962612e-14 | 0.7894734 | 0.985 | 0.917 | 8.355053e-10 |
| Ccl24 | 5.181960e-14 | 1.9767242 | 0.288 | 0.060 | 8.724348e-10 |
| Klf2 | 6.068075e-14 | -2.4356510 | 0.227 | 0.671 | 1.021621e-09 |
| Tubb4b | 6.393222e-14 | 1.2912429 | 0.591 | 0.246 | 1.076363e-09 |
| Ybx1 | 2.379939e-13 | 1.1381948 | 0.879 | 0.666 | 4.006865e-09 |
| Mcm3 | 3.252660e-13 | 0.7390529 | 0.333 | 0.081 | 5.476178e-09 |
| Tyrobp | 5.182101e-13 | 0.9211215 | 0.712 | 0.282 | 8.724585e-09 |
| Cks1b | 7.362173e-13 | 0.9860740 | 0.455 | 0.153 | 1.239495e-08 |
| Anp32b | 1.484673e-12 | 1.1777789 | 0.742 | 0.485 | 2.499595e-08 |
| Actb | 1.544930e-12 | 0.7983446 | 1.000 | 0.996 | 2.601044e-08 |
| Mgl2 | 2.124704e-12 | 0.7859253 | 0.258 | 0.053 | 3.577152e-08 |
| Prdx1 | 2.152922e-12 | 0.8891464 | 0.939 | 0.609 | 3.624660e-08 |
| Slc25a5 | 2.910747e-12 | 1.1827808 | 0.788 | 0.551 | 4.900534e-08 |
| Tmpo | 3.331811e-12 | 1.1804920 | 0.621 | 0.297 | 5.609437e-08 |
| Npl | 4.127494e-12 | 0.9023949 | 0.273 | 0.060 | 6.949049e-08 |
| Clec4n | 4.785634e-12 | 0.8349504 | 0.409 | 0.119 | 8.057093e-08 |
| Ccl12 | 4.791061e-12 | 1.2250966 | 0.303 | 0.073 | 8.066230e-08 |
| 1810037I17Rik | 5.877583e-12 | 1.1257415 | 0.803 | 0.482 | 9.895499e-08 |
| Dck | 1.398086e-11 | 0.9412996 | 0.394 | 0.121 | 2.353817e-07 |
| Hint1 | 1.475023e-11 | 0.9495056 | 0.864 | 0.643 | 2.483348e-07 |
| Foxp1 | 3.507720e-11 | -1.5888010 | 0.197 | 0.602 | 5.905597e-07 |
| Cox5a | 3.572184e-11 | 1.2086200 | 0.788 | 0.575 | 6.014129e-07 |
| Pycard | 3.851142e-11 | 1.2514387 | 0.561 | 0.248 | 6.483783e-07 |
| Hmgb1 | 4.759003e-11 | 1.0910087 | 0.712 | 0.436 | 8.012258e-07 |
| Ntpcr | 9.674735e-11 | 1.0416145 | 0.394 | 0.134 | 1.628838e-06 |
